## Supplementary Figures and Tables for "Molecular estimation of neurodegeneration pseudotime in older brains"

Supplementary Figures and Tables for Mukherjee et al., *Molecular estimation of neurodegeneration pseudotime in older brains.*

**Supplemental Figures**

**Supplementary Figure 1 -** Comparison between trajectories inferred using different gene sub-set selection methods: i) Differential Expression with an FDR cut-off of 0.1, ii) High variance gene selection.

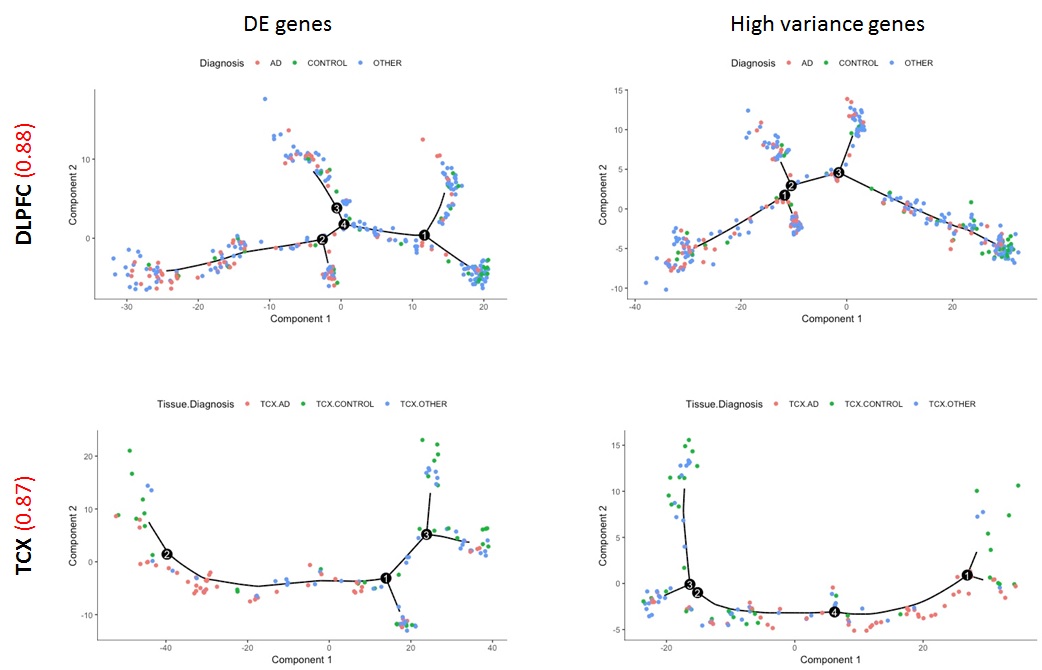

**Supplementary Figure 2 –** Trajectories with DE genes at FDR p-value <= 0.01 in TCX and DLFPC brain regions.

**Supplementary Figure 3 –** Patient trajectory maps for TCX data by adjustment A) Adjusted for PMI, B) Adjusted for first 10 PCs, C) Adjusted for RIN, D) Adjusted for RIN, PMI, and first 10 PCs.

DLPFC

TCX

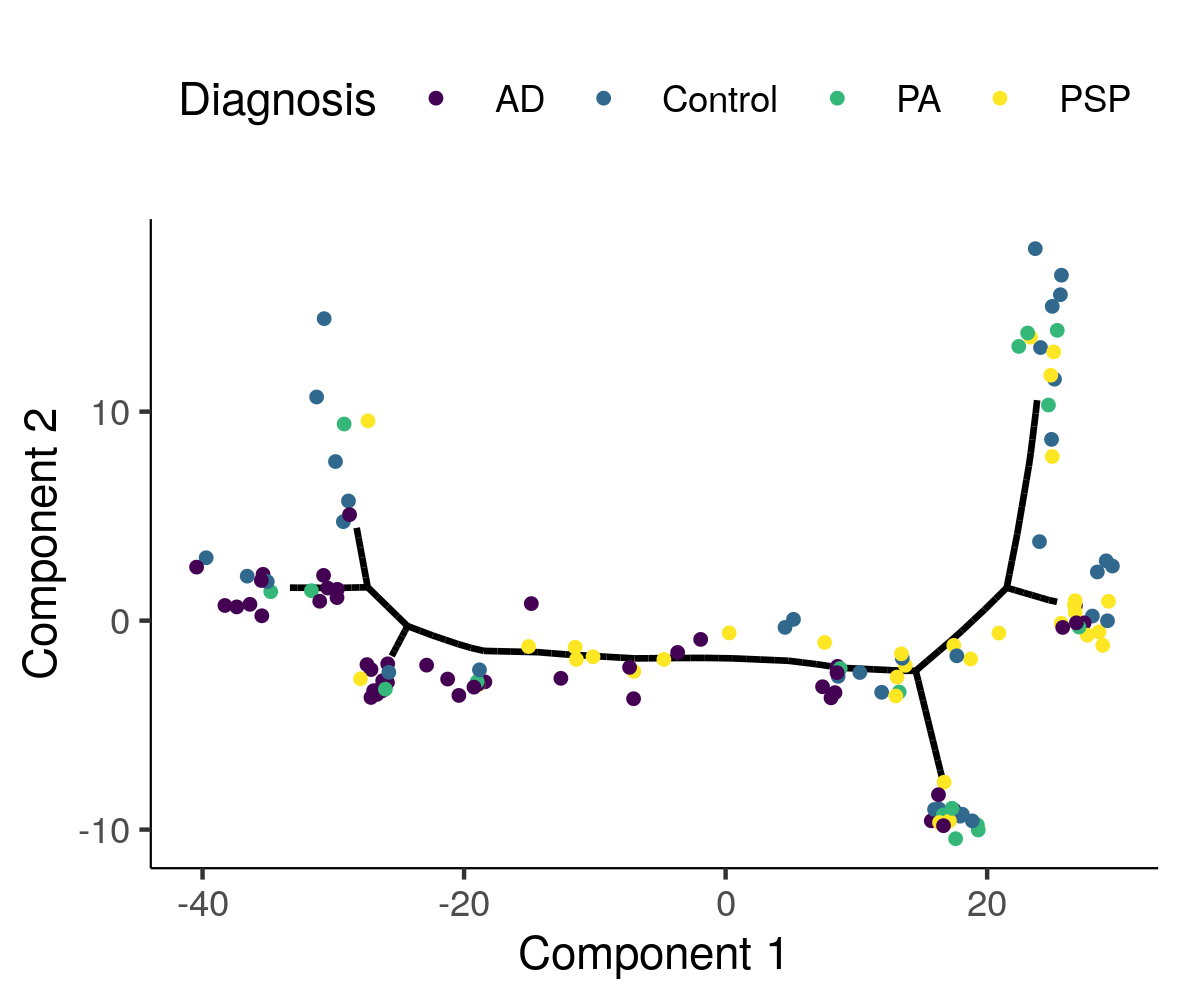

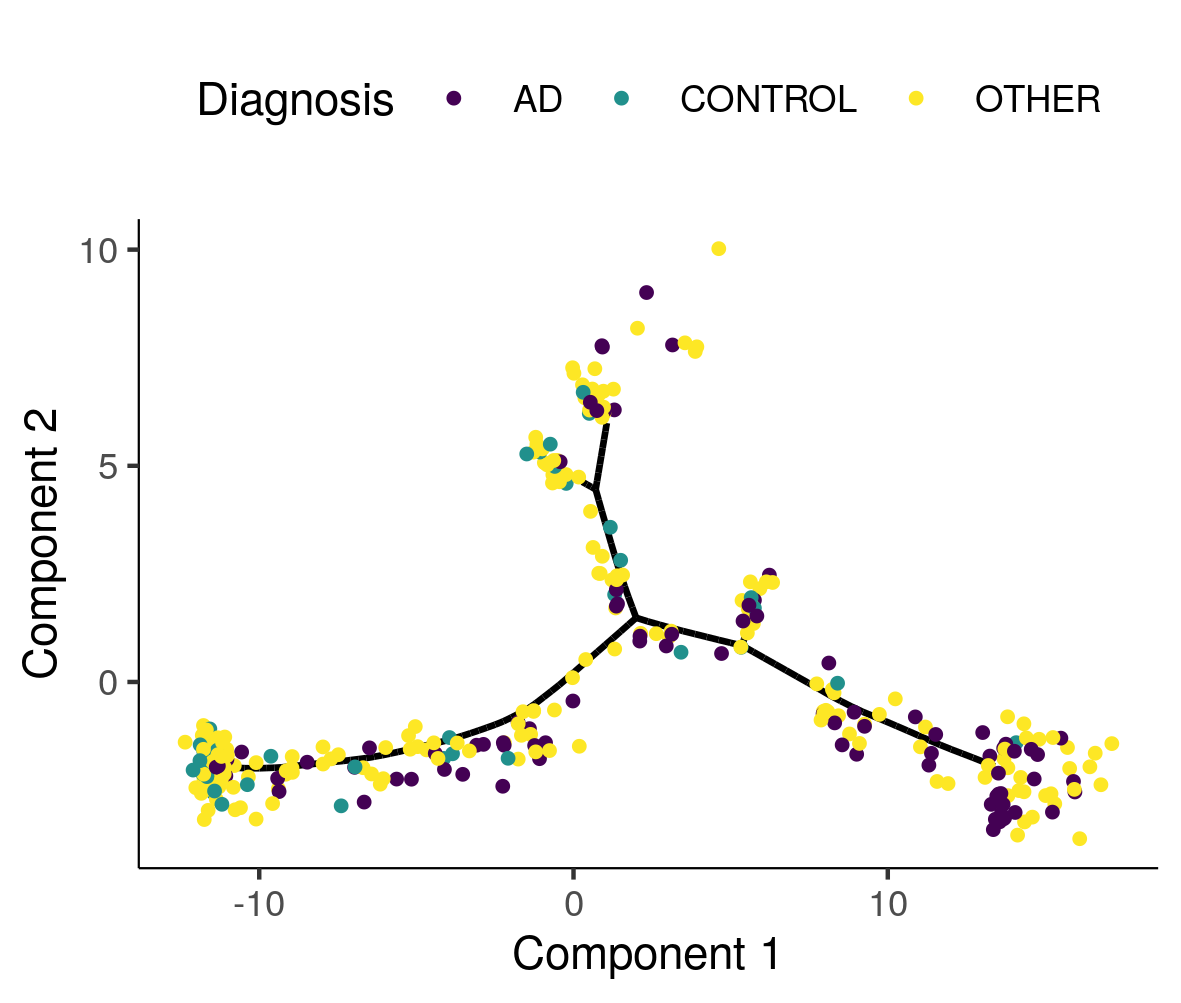

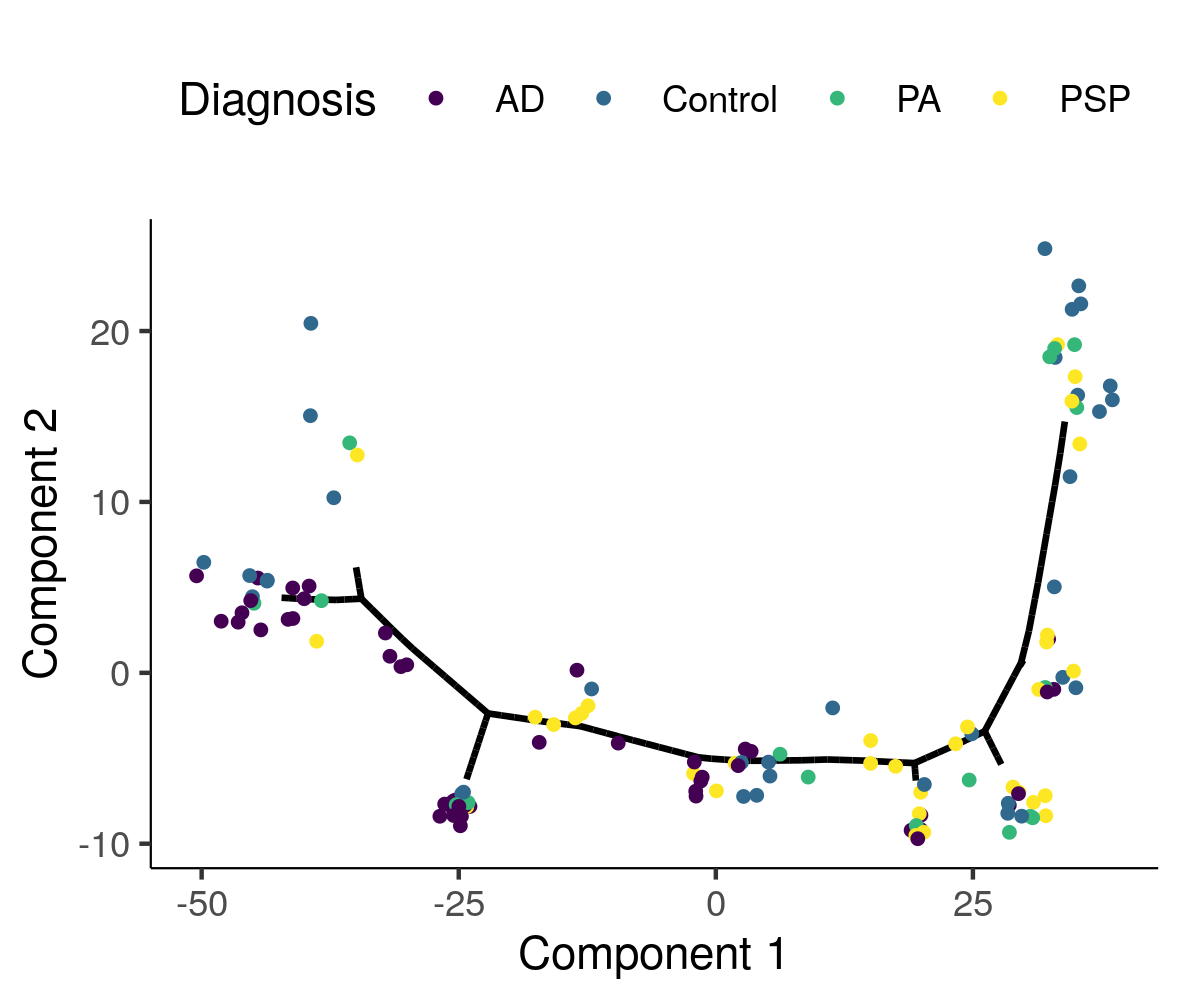

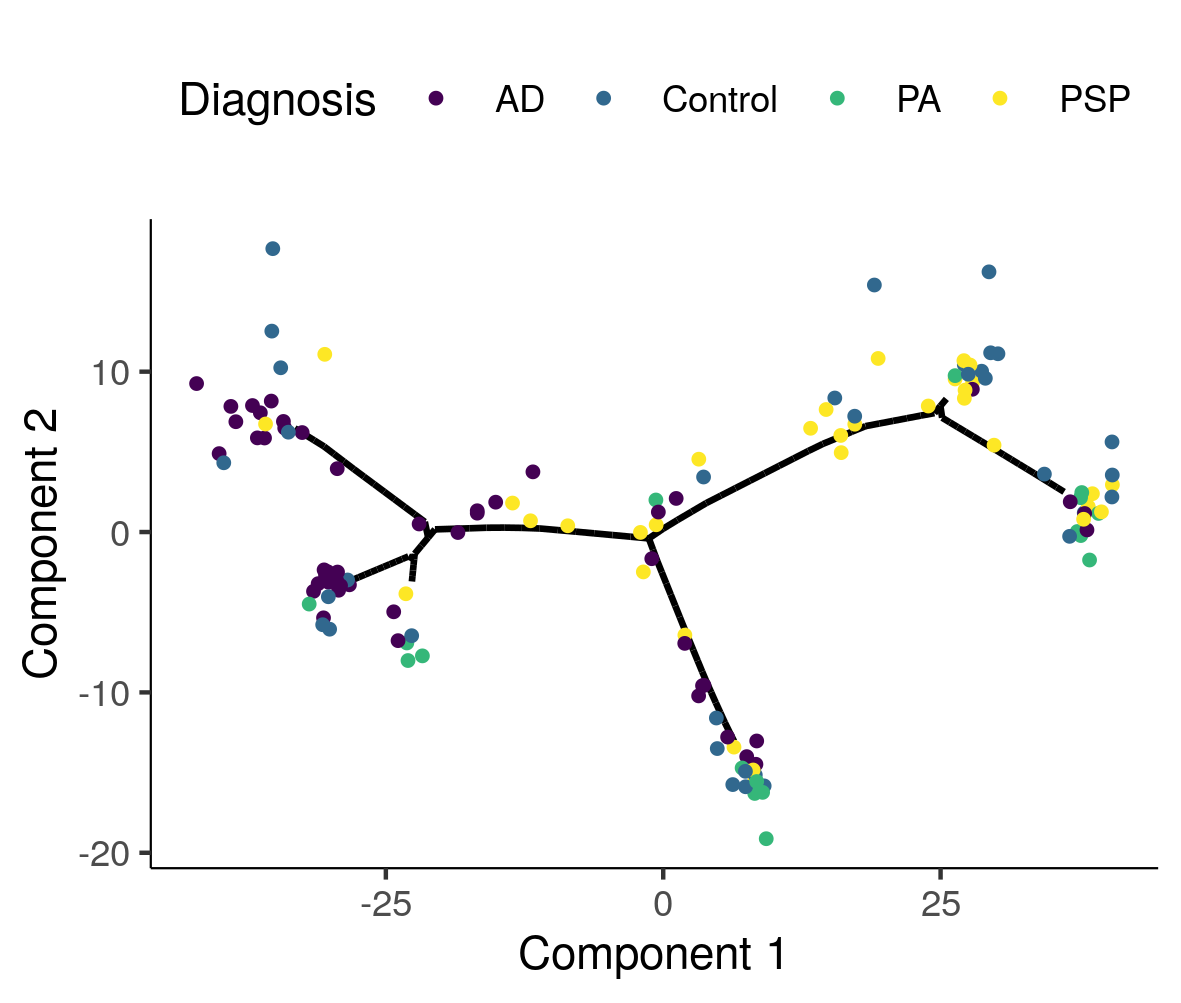

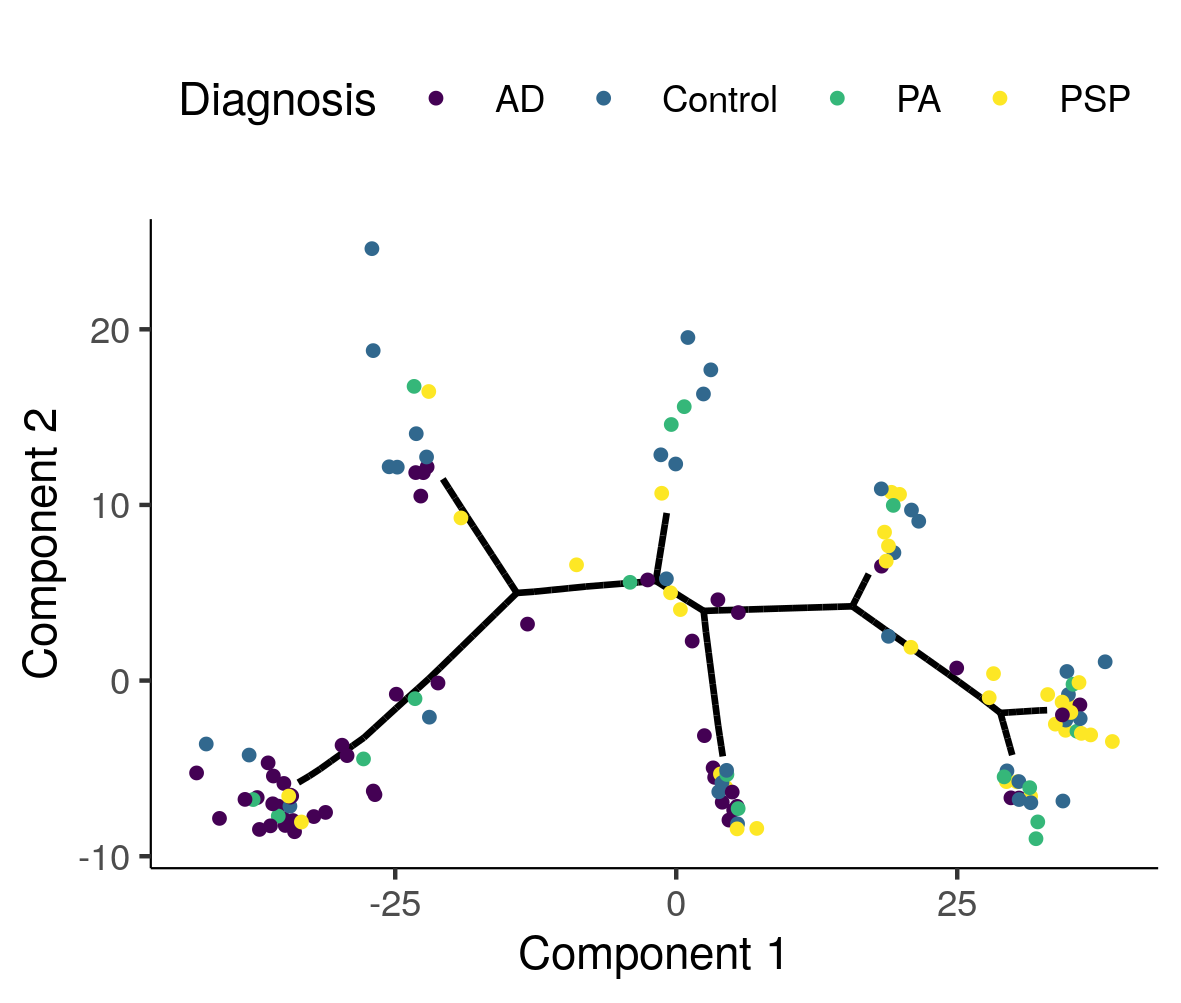

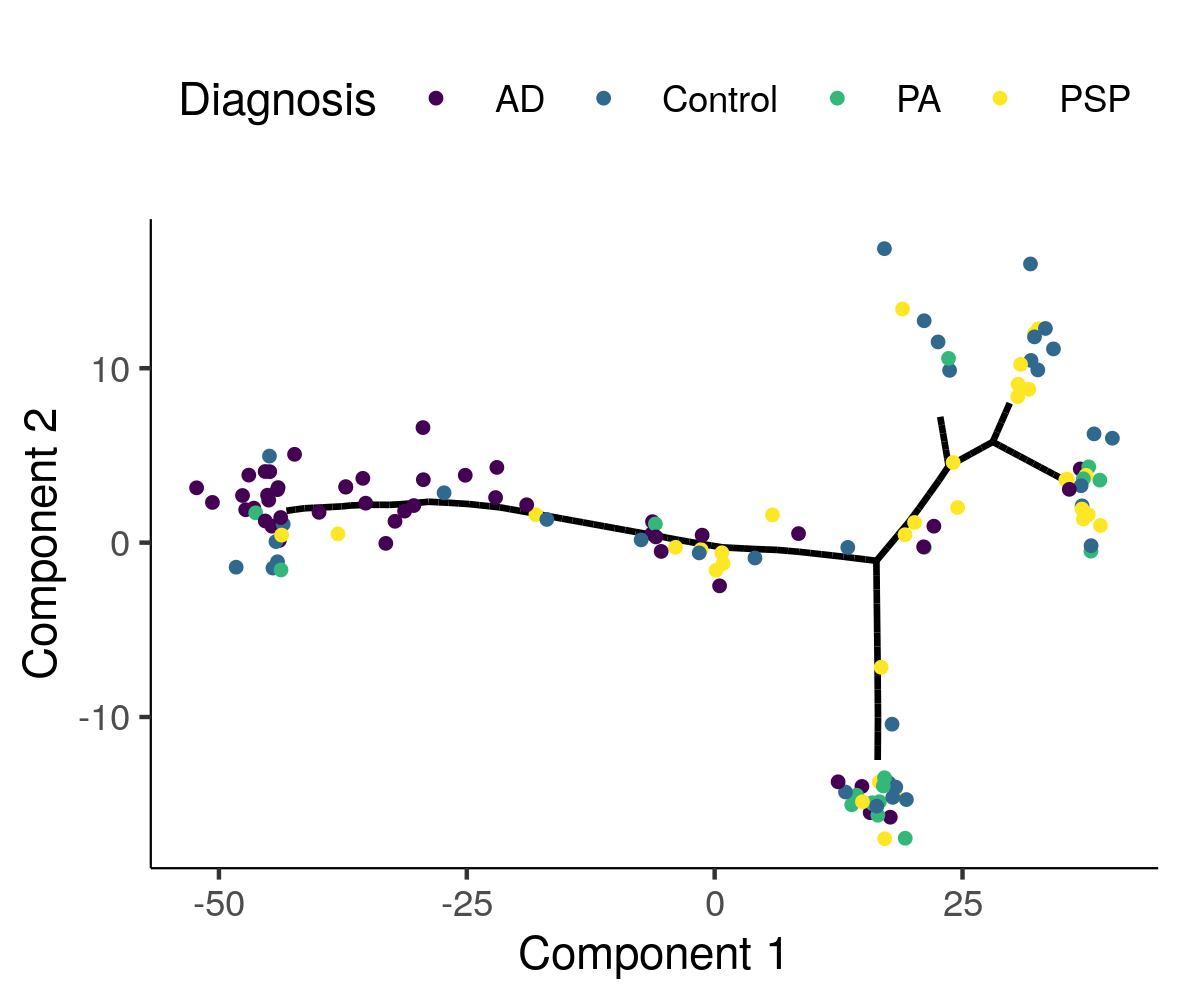

A

D

C

B

**Supplementary Figure 4 –** Patient trajectory maps for DLPFC data by adjustment A) Adjusted for PMI, B) Adjusted for first 10 PCs, C) Adjusted for RIN, D) Adjusted for RIN, PMI, and first 10 PCs.

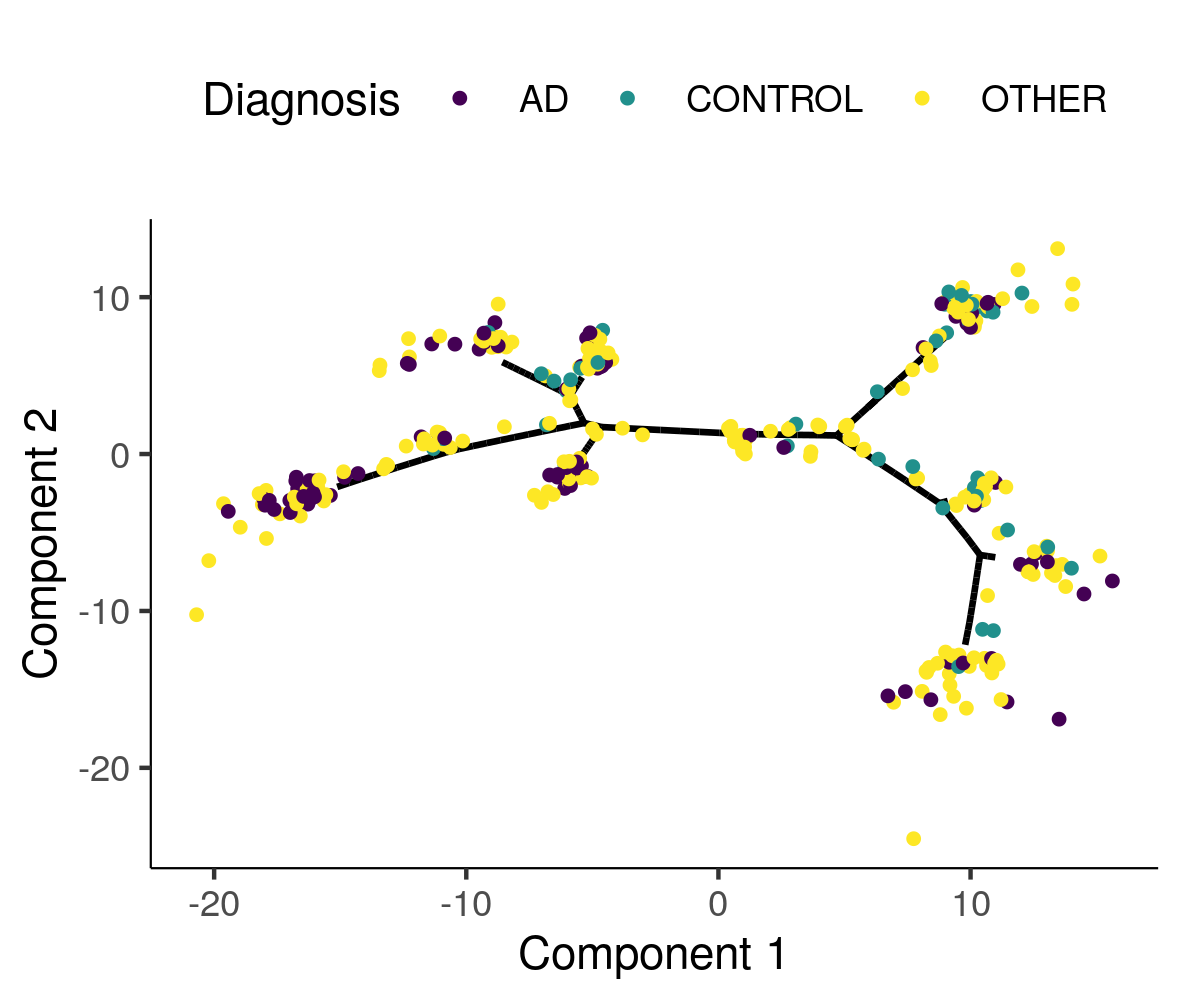

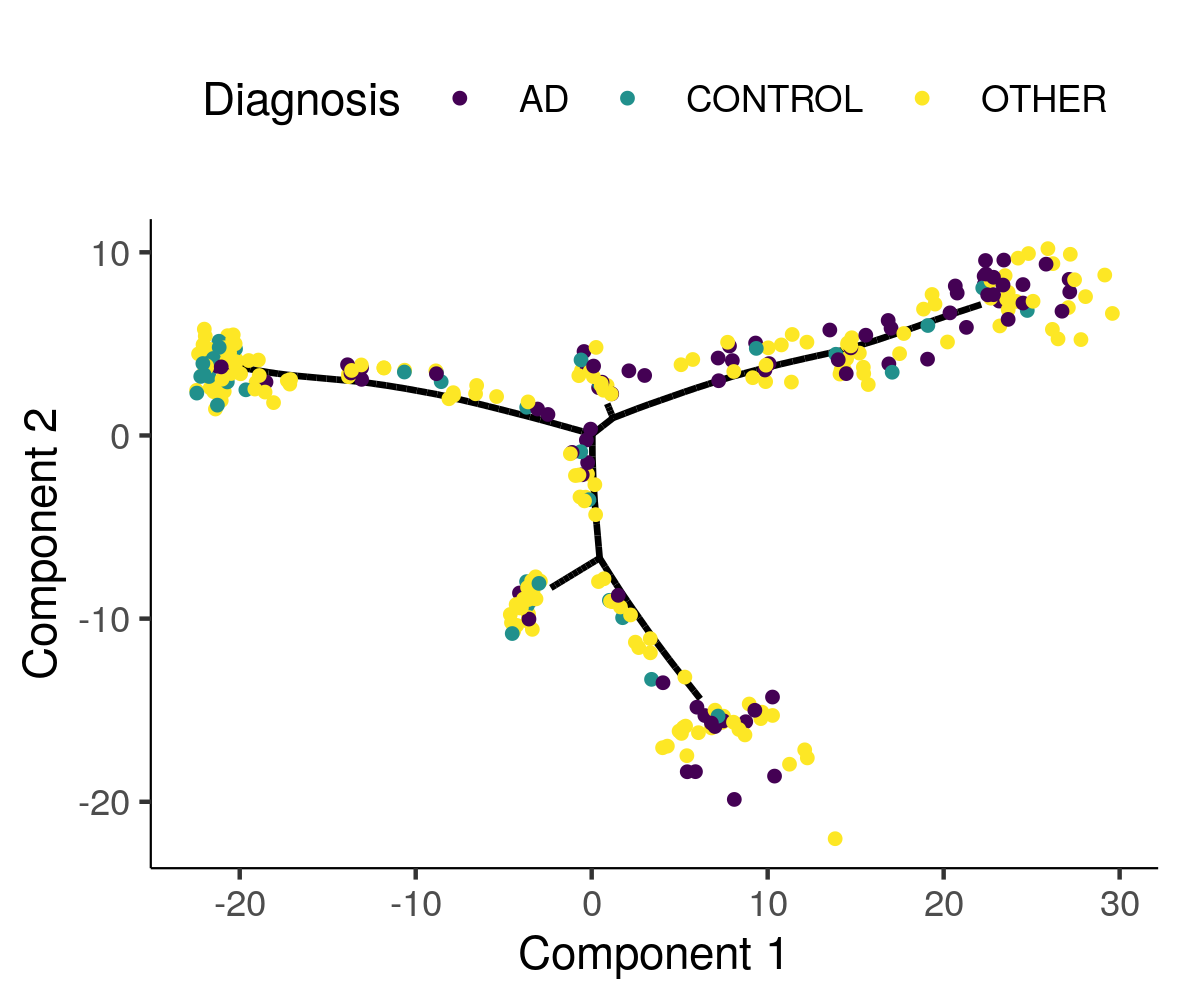

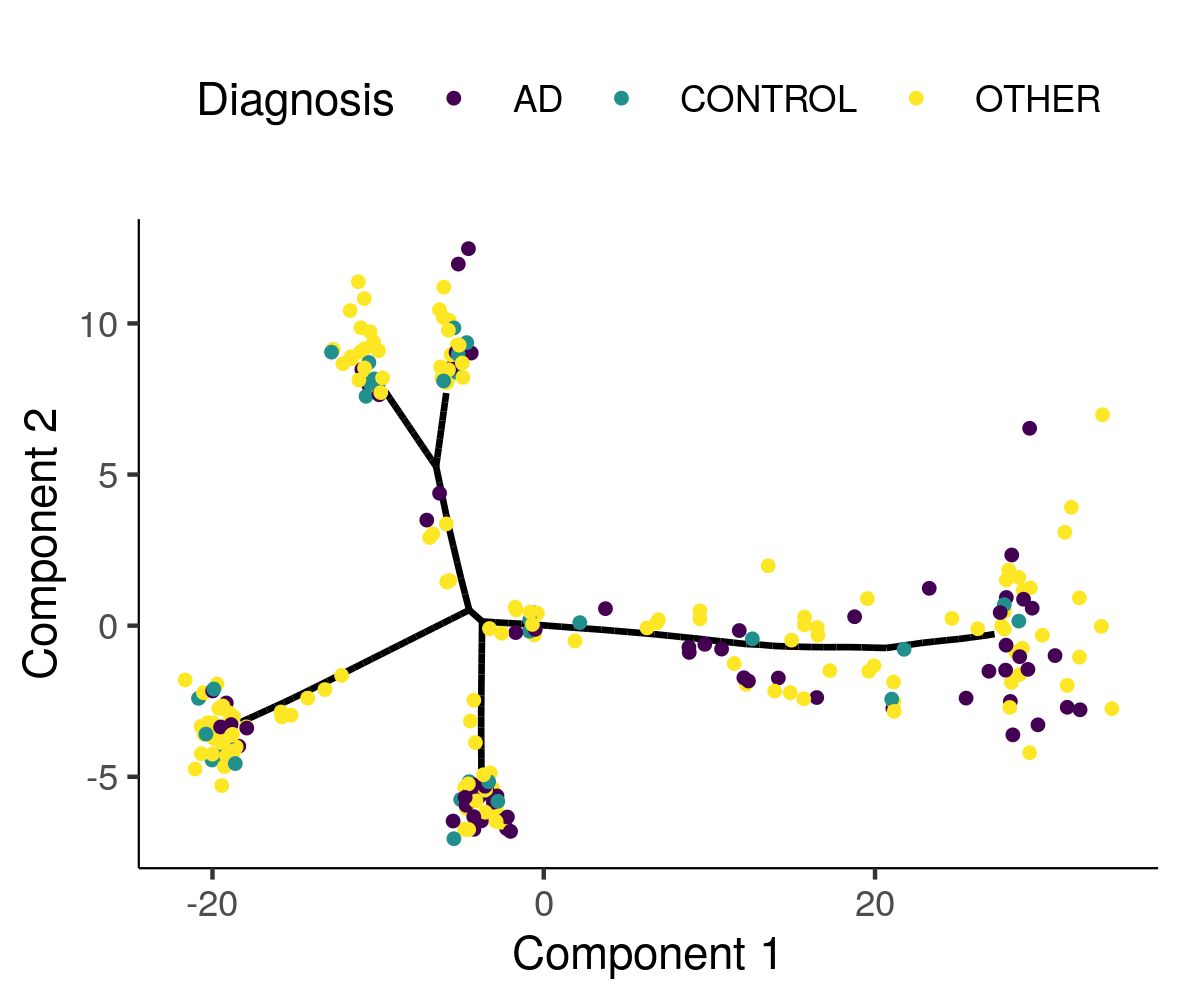

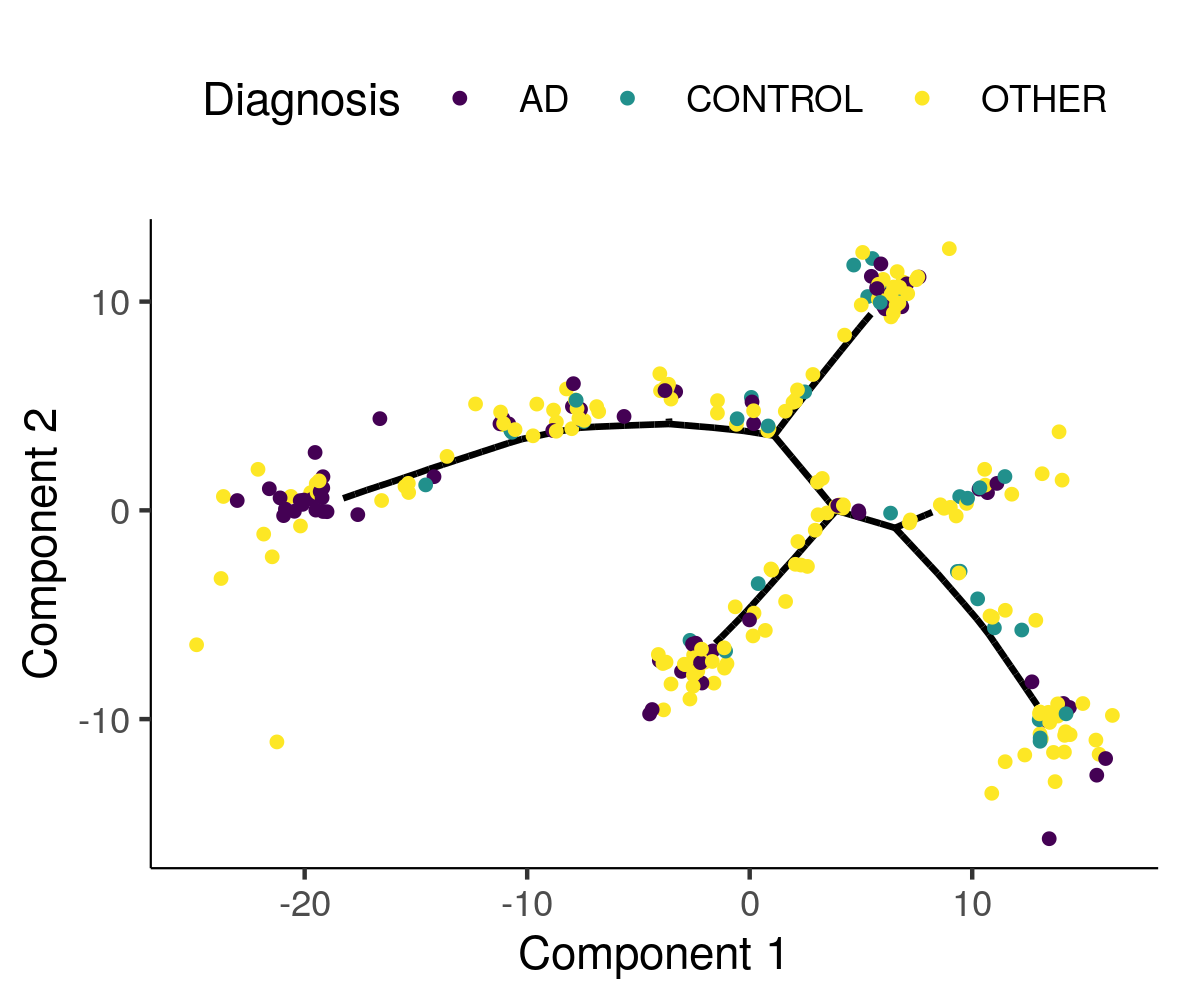

A

D

C

B

**Supplementary Figure 5 –** Absolute value of Pearson correlations between pseudotime estimated with all samples in both ROSMAP (DLPFC) and Mayo RNAseq (TCX), and pseudotime estimated with leave one out data-sets.

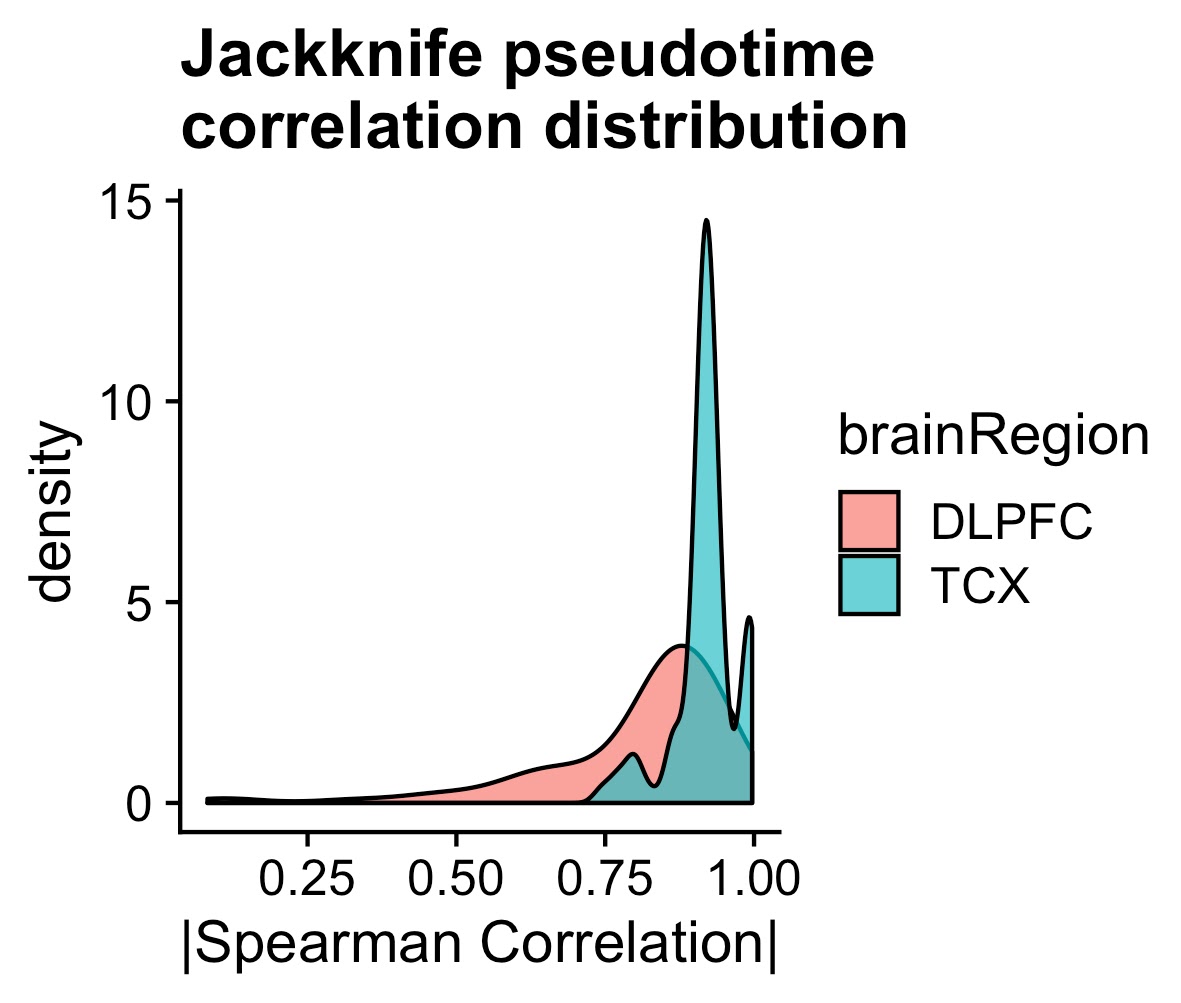

**Supplementary Figure 6 –** Pseudotime by AD case-control status for 218 independent samples from two independent studies. A) Adjusted for PMI, B) Adjusted for first 10 PCs, C) Adjusted for RIN, D) Adjusted for PMI, RIN, and first 10 PCs. Box plots have lower and upper hinges at the 25th and 75th percentiles and whiskers extending to at most 1.5xIQR (interquartile range).

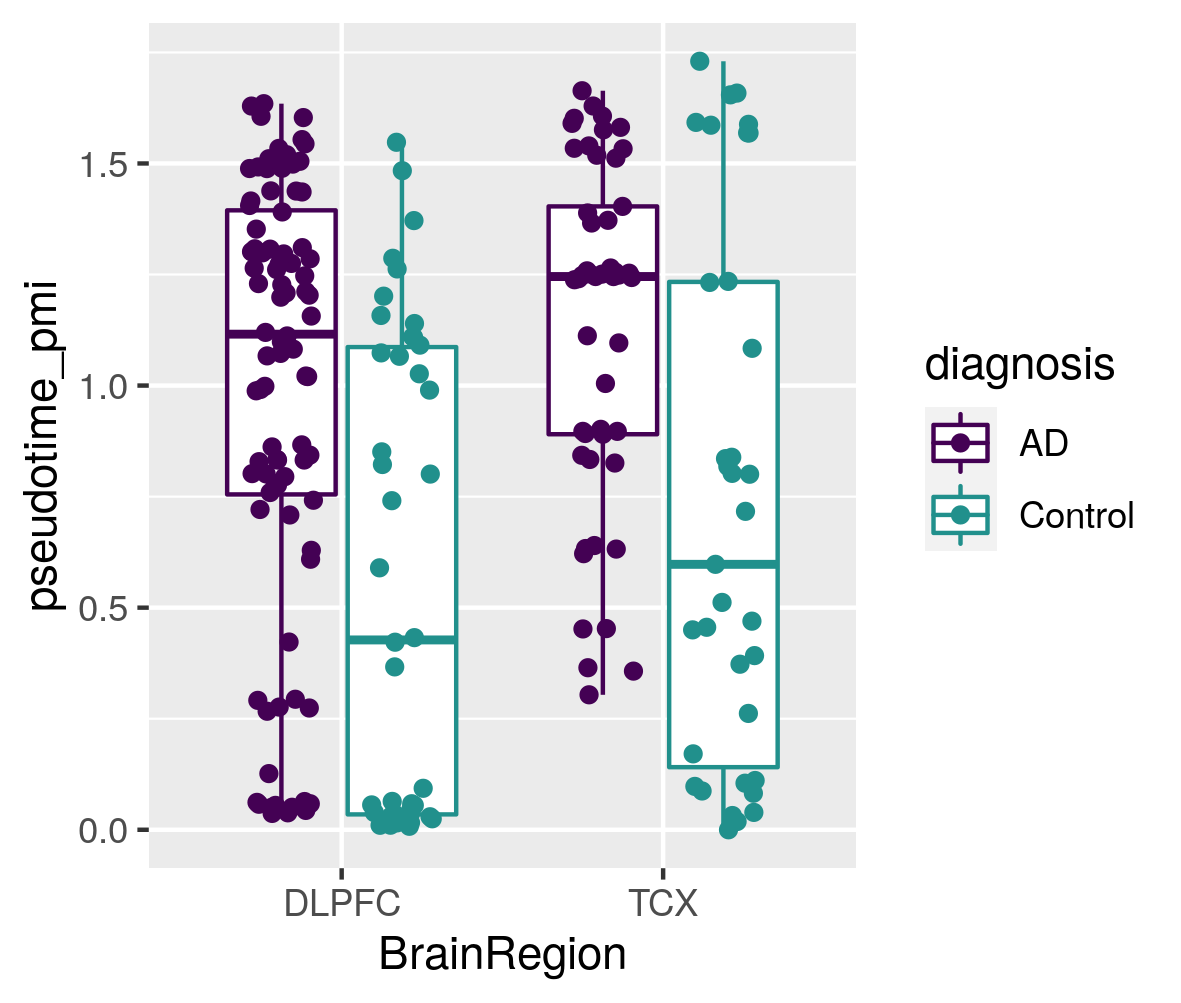

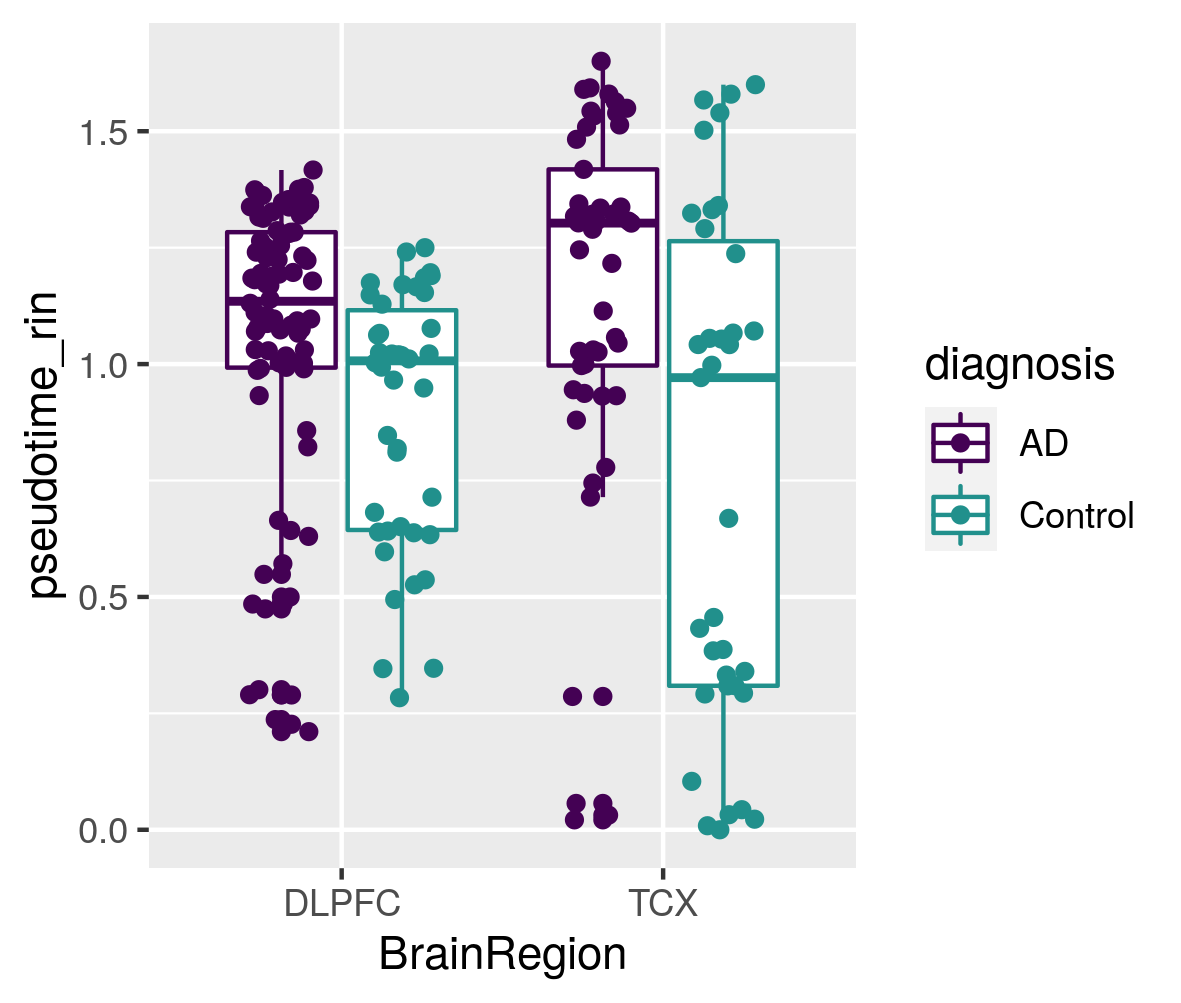

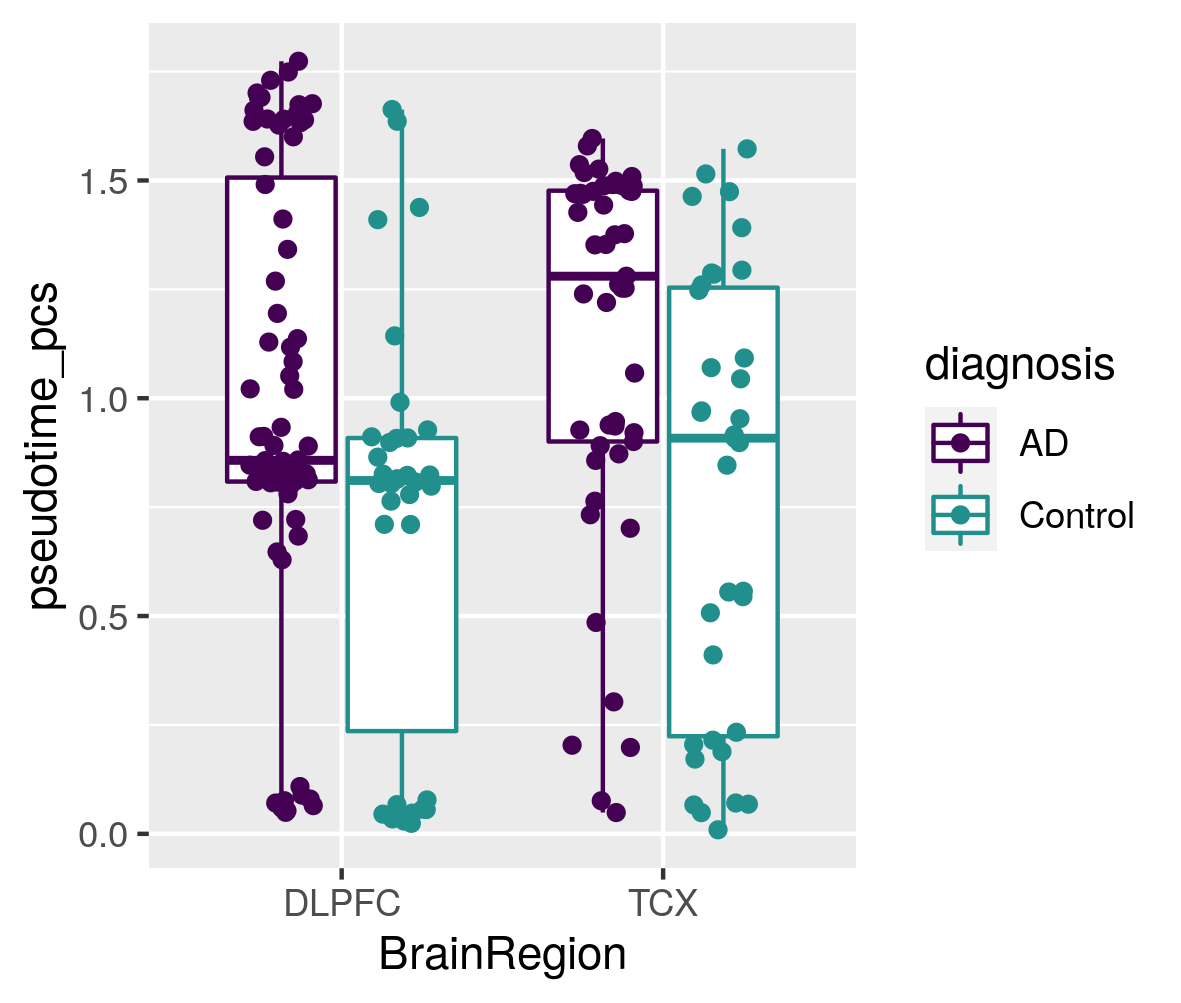

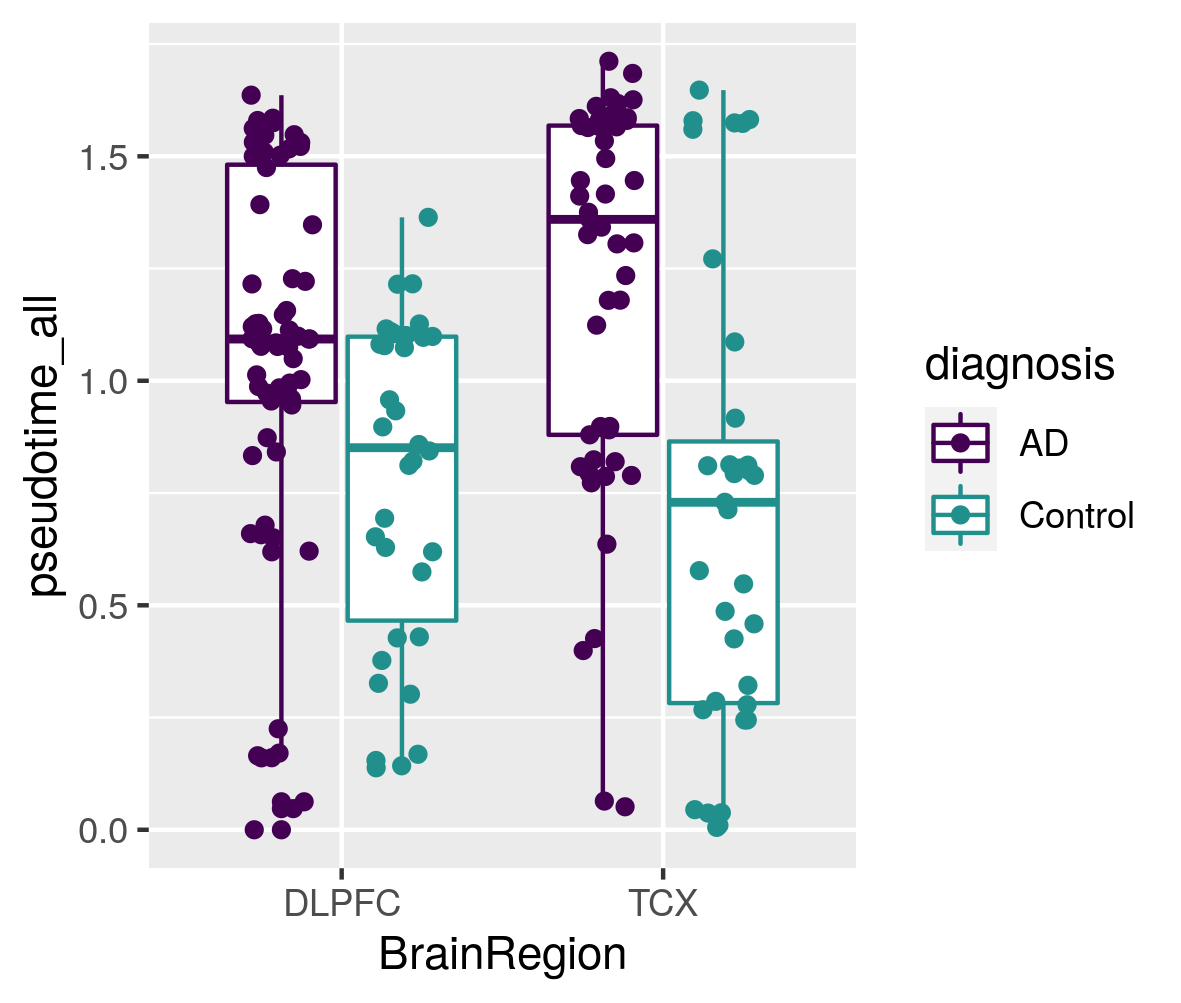

A

D

C

B

**Supplementary Figure 7 –** Lineage inference in the Mayo eGWAS expression array data-set for 186 independent samples from one independent study. A) Monocle2 inferred manifold, B) disease state as a function of disease pseudotime, C) APOE e4 dosage as a function of disease pseudotime, D) resistant individuals on the disease pseudotime manifold. Box plots have lower and upper hinges at the 25th and 75th percentiles and whiskers extending to at most 1.5xIQR (interquartile range).

**Supplementary Figure 8 -** Manifold learning infers disease states from RNA-seq samples, samples from males only for 143 independent samples from two independent studies. A) Estimated cell trajectory from A) TCX and B) DLPFC. C) Distribution of pseudotime for AD cases and controls for DLPFC and TCX. D) Distribution of expression correlation with pseudotime for both LOAD GWAS genes and non-LOAD GWAS genes. Box plots have lower and upper hinges at the 25th and 75th percentiles and whiskers extending to at most 1.5xIQR (interquartile range).

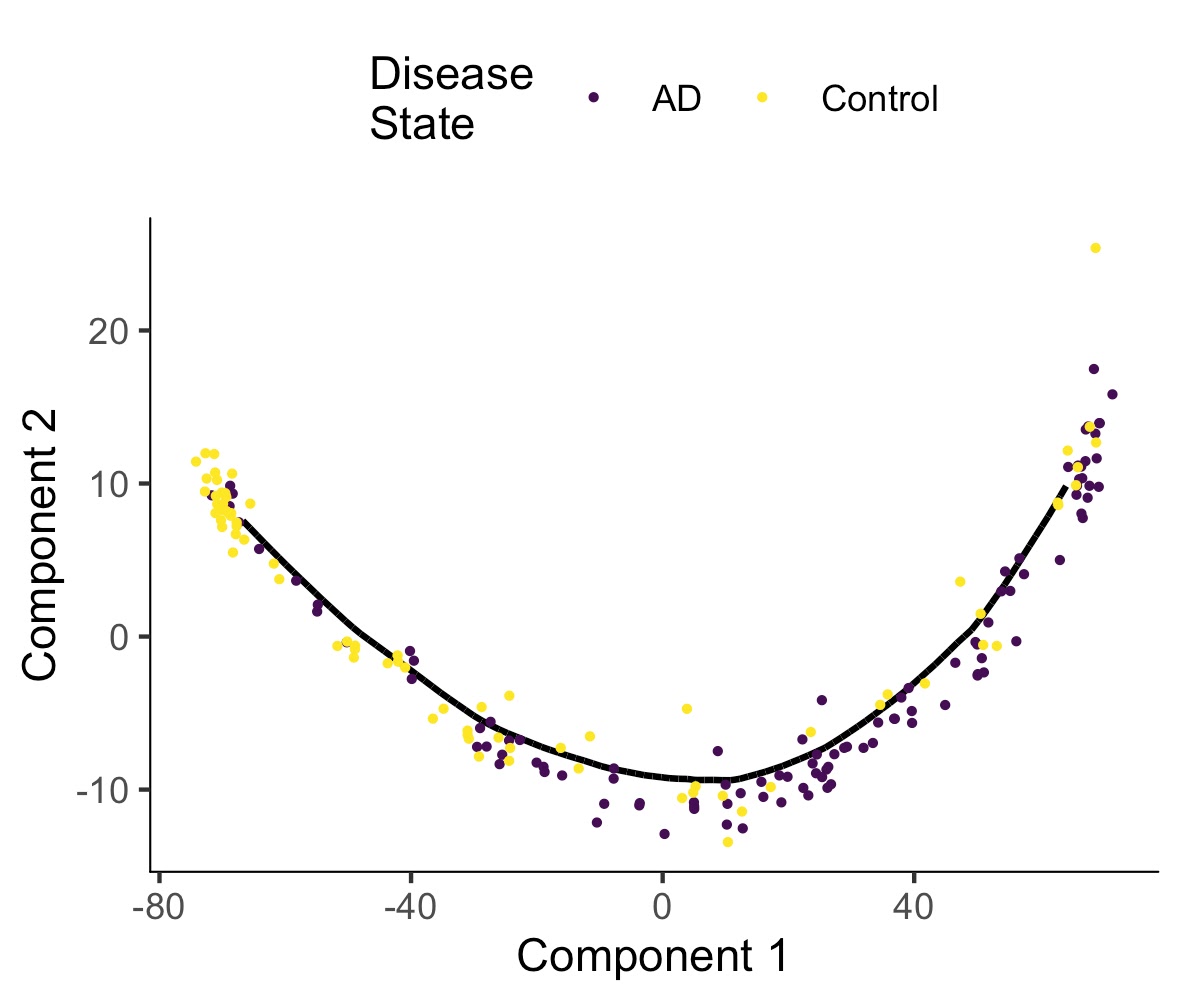

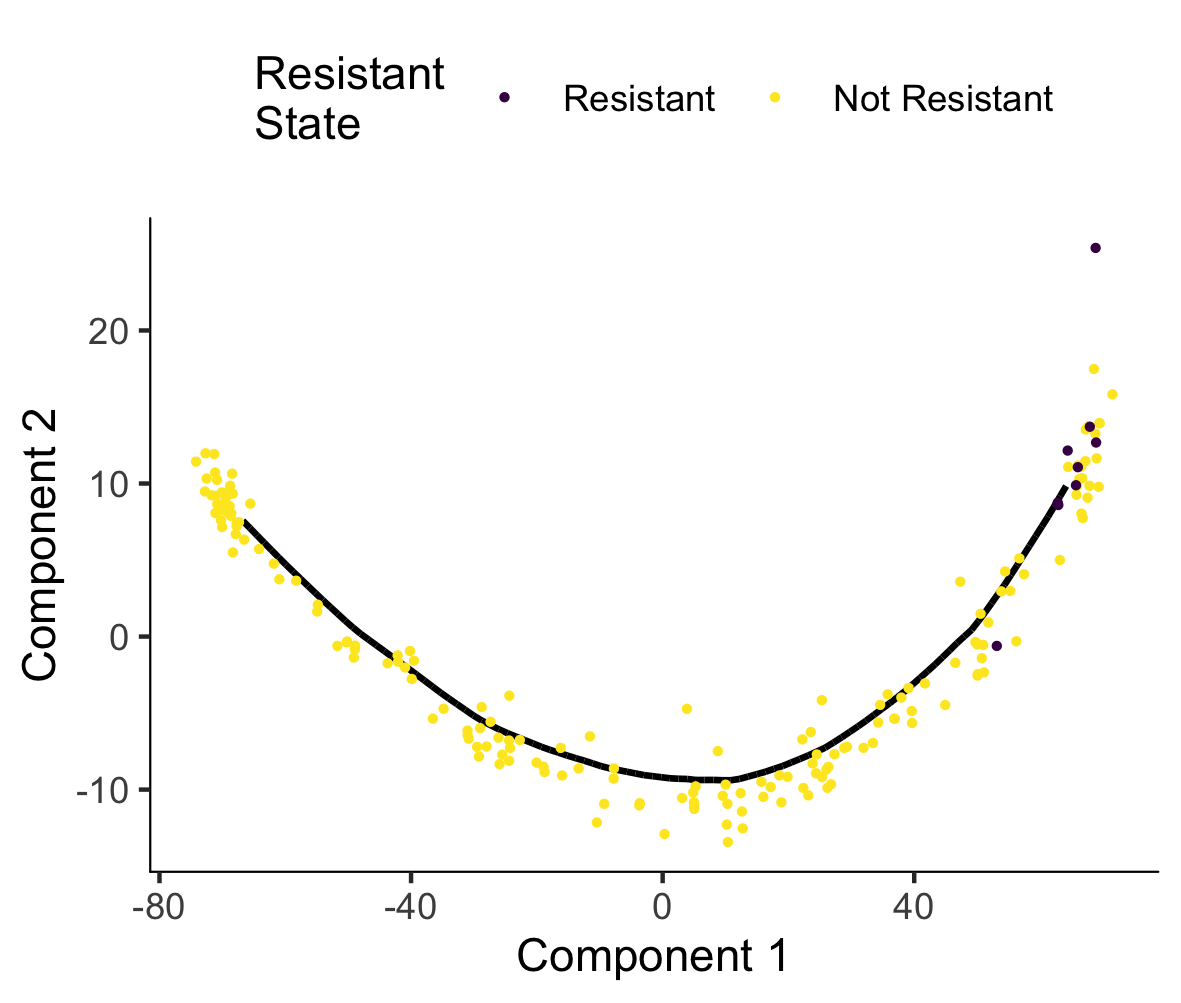

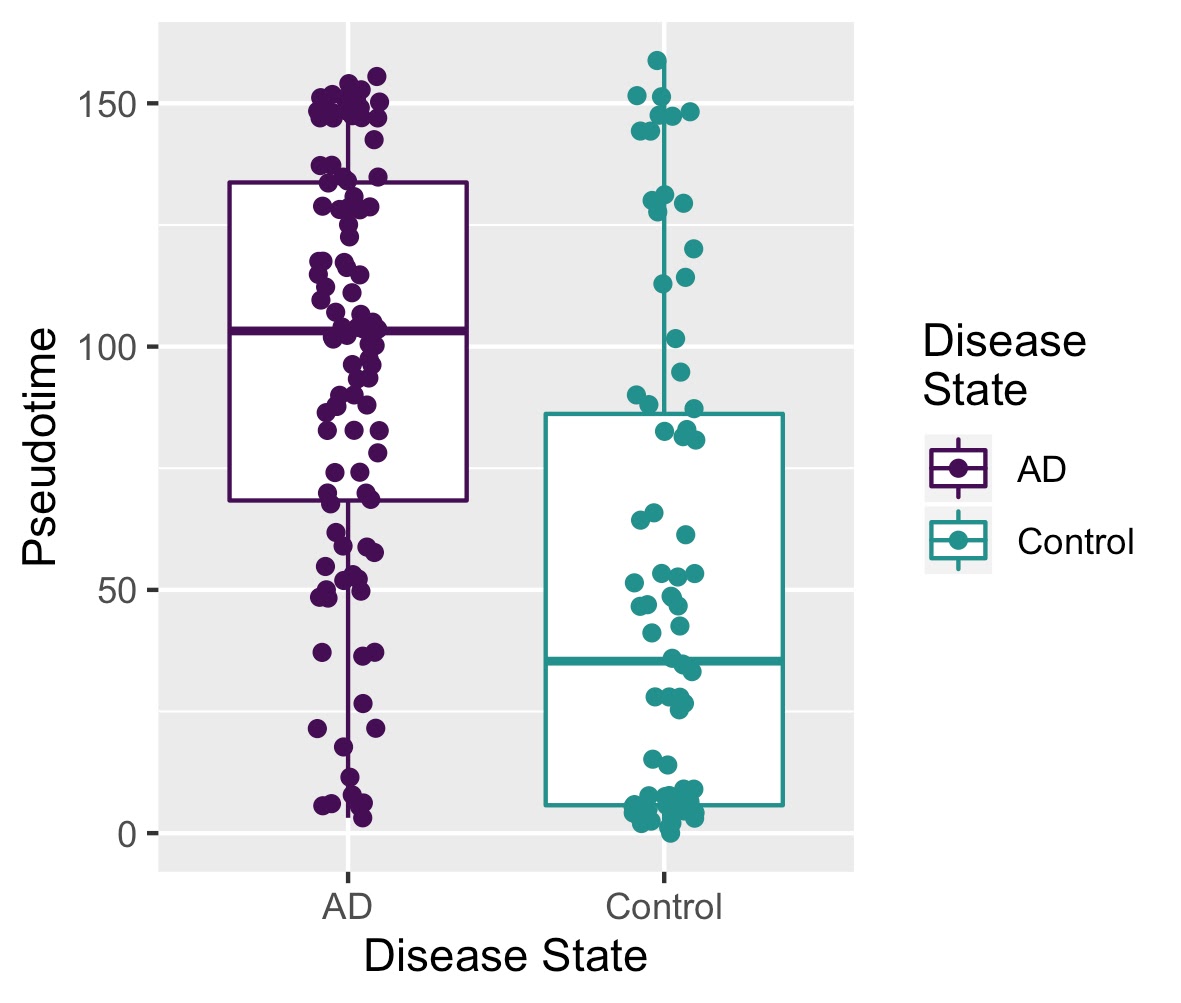

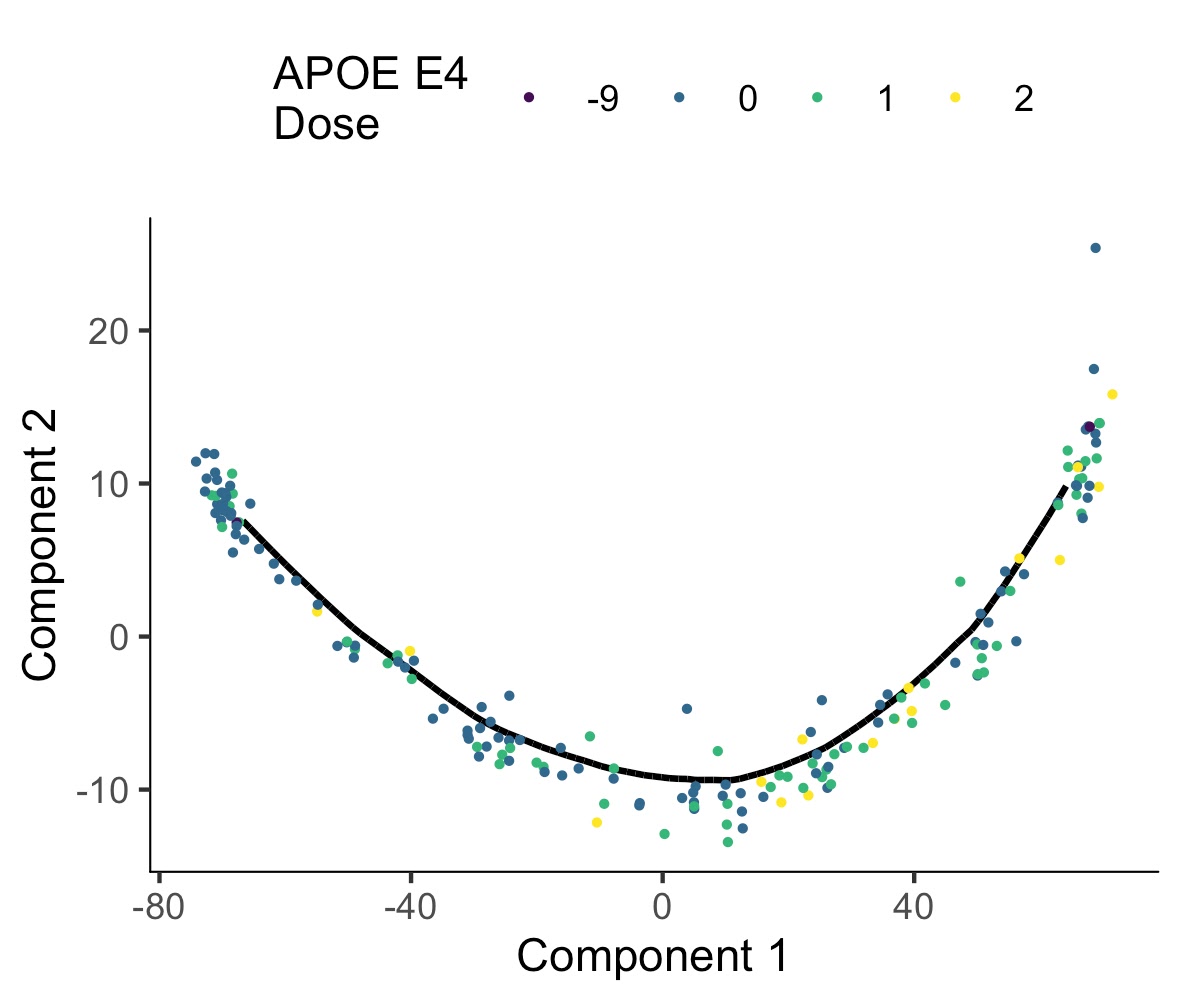

A

D

C

B

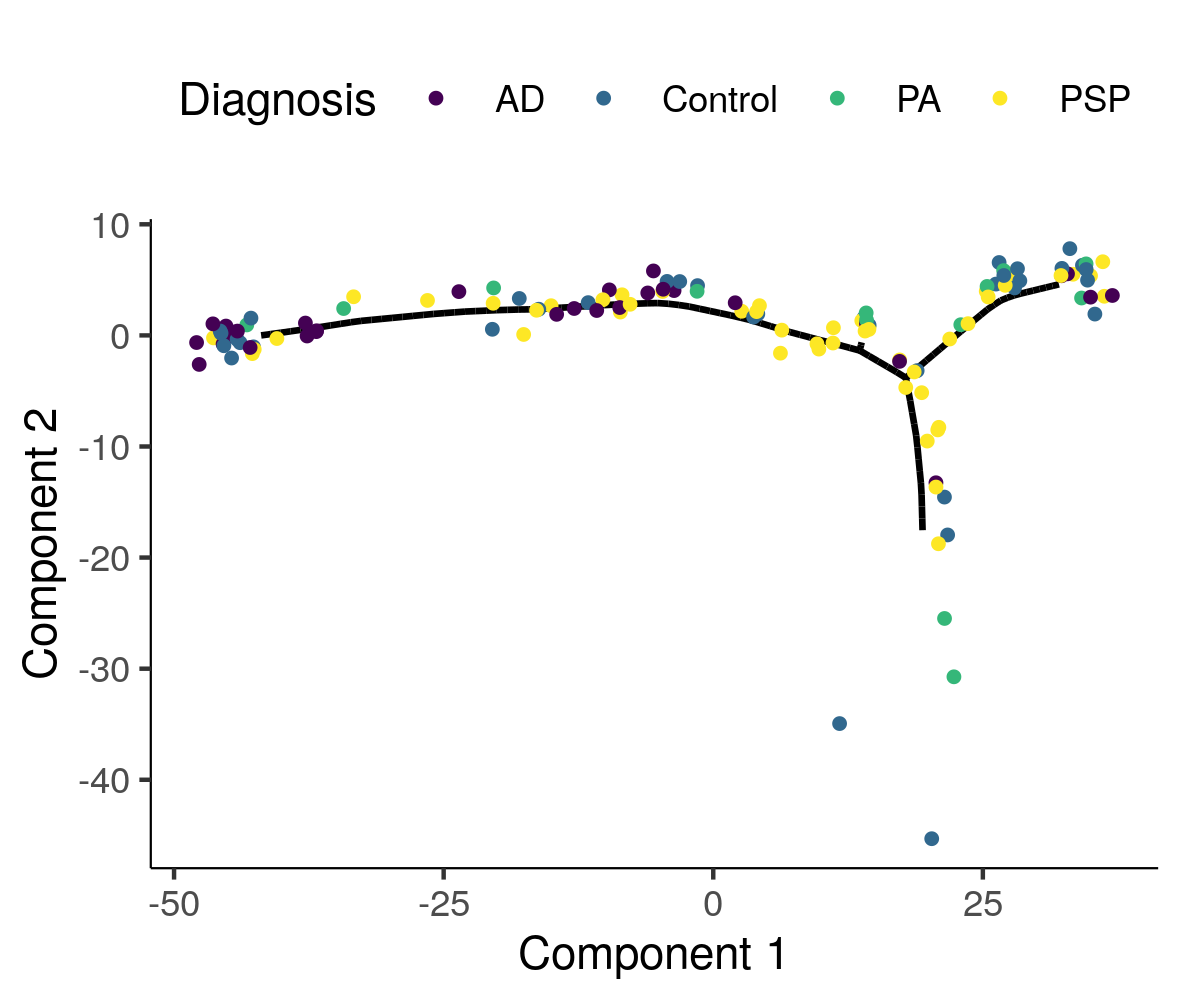

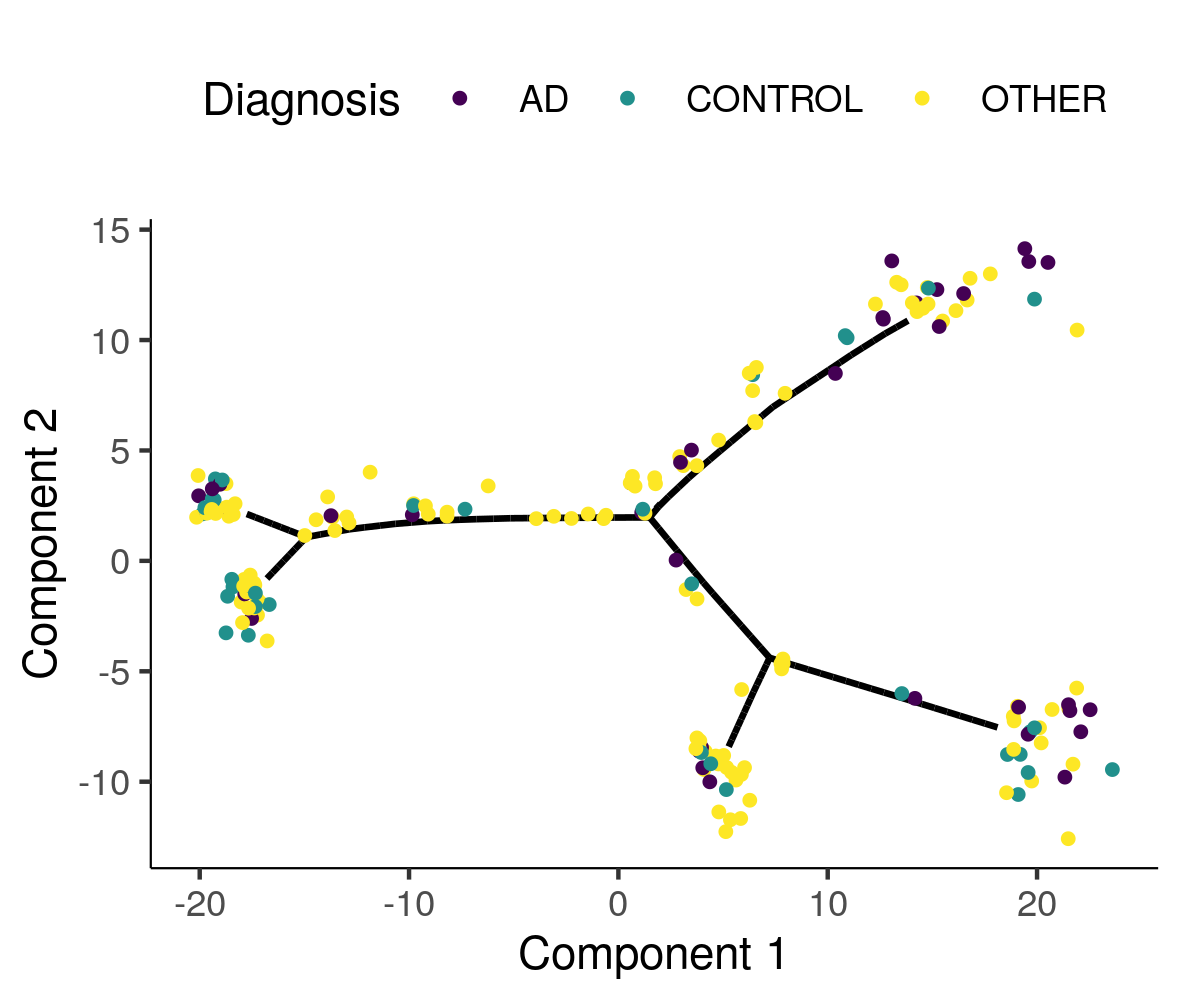

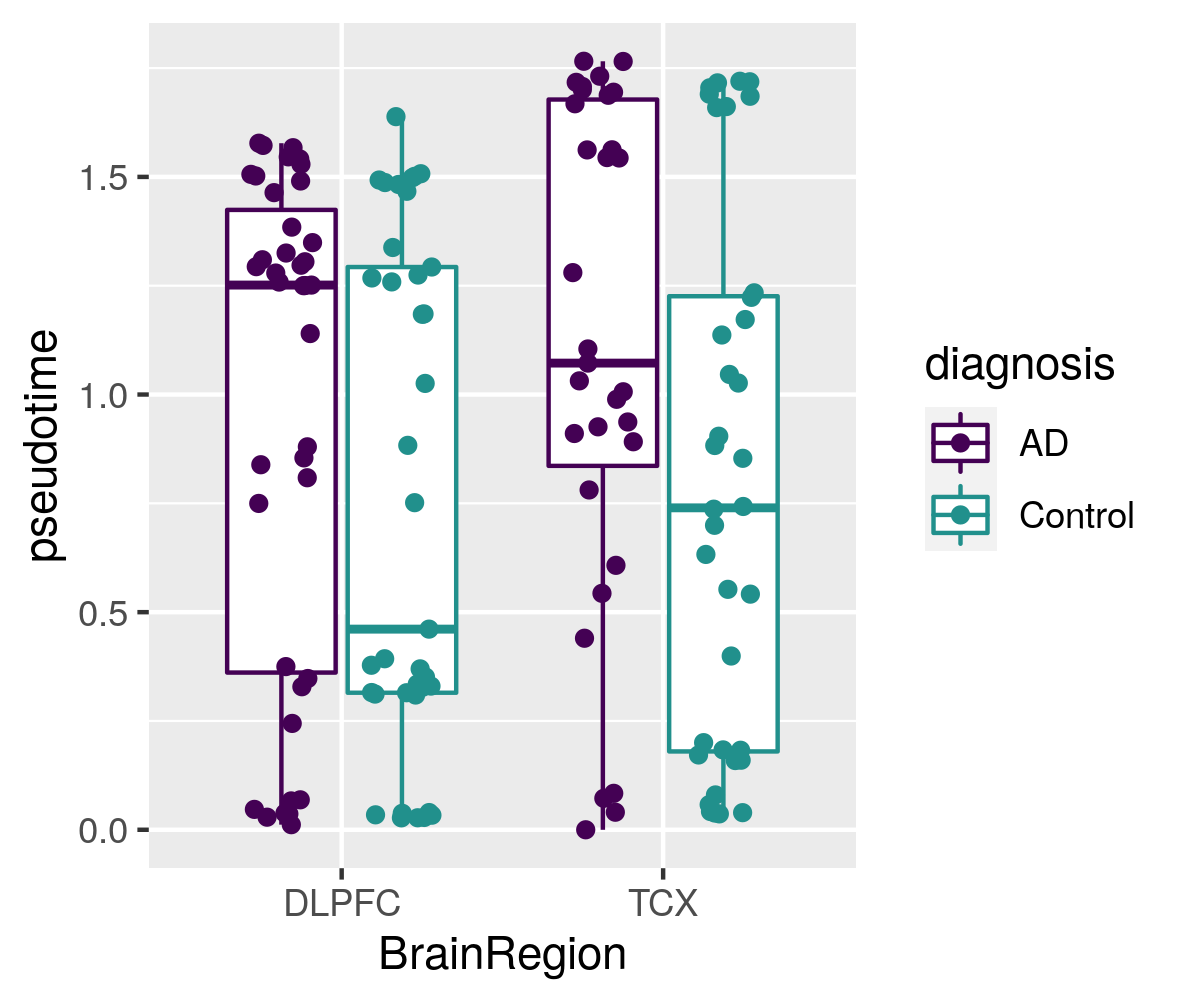

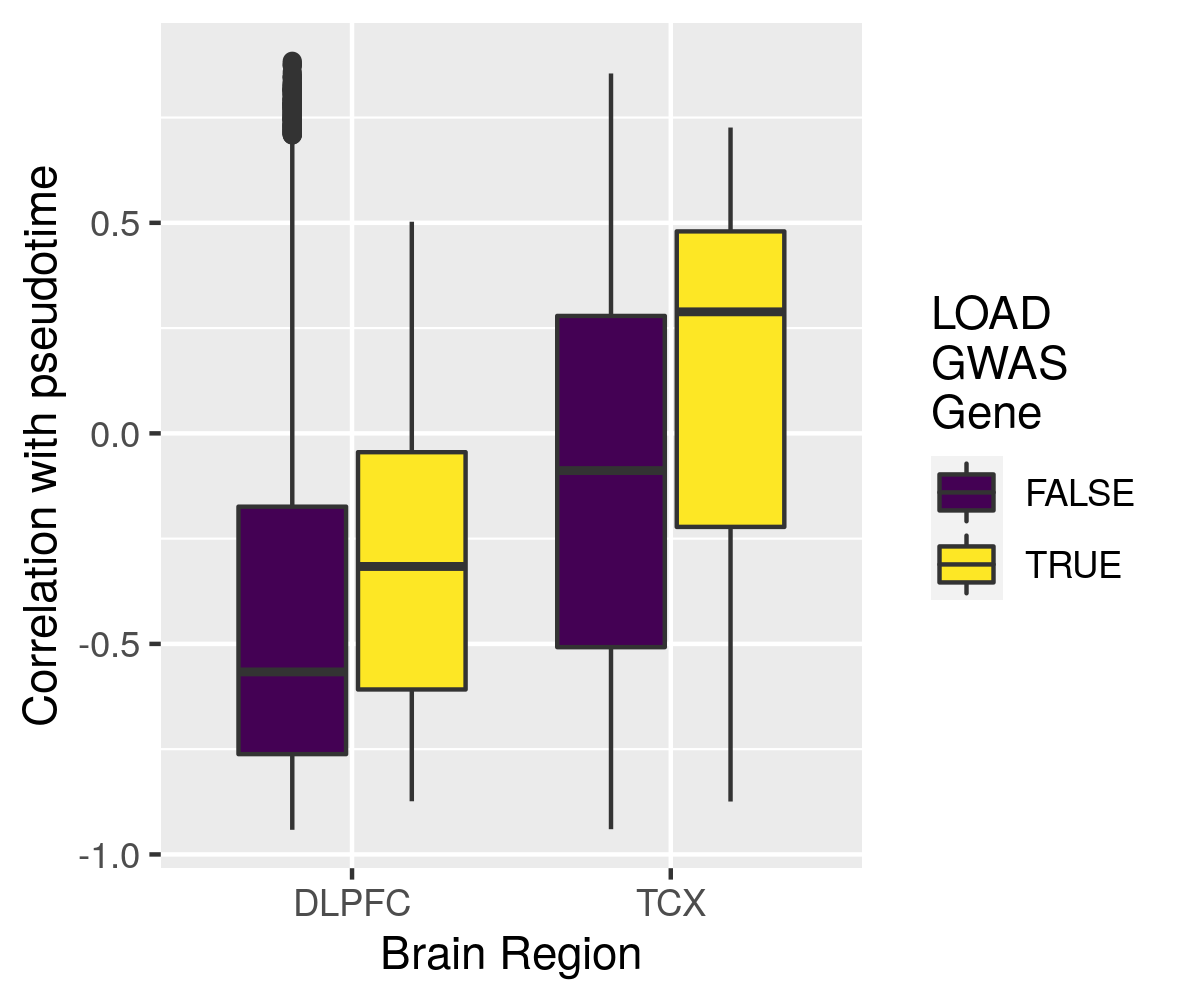

A

D

C

B

**Supplementary Figure 9 -** Manifold learning and measures of staging in LOAD in DLPFC samples, males only for 143 independent samples from two independent studies. A-C) Samples colored by three external measures of LOAD staging: Braak score, CERAD score, and cognitive diagnosis. D-F) Distribution of samples by inferred stage for different distinct stages in each of the three methods of measuring LOAD severity. Box plots have lower and upper hinges at the 25th and 75th percentiles and whiskers extending to at most 1.5xIQR (interquartile range).

F

E

D

A

B

C

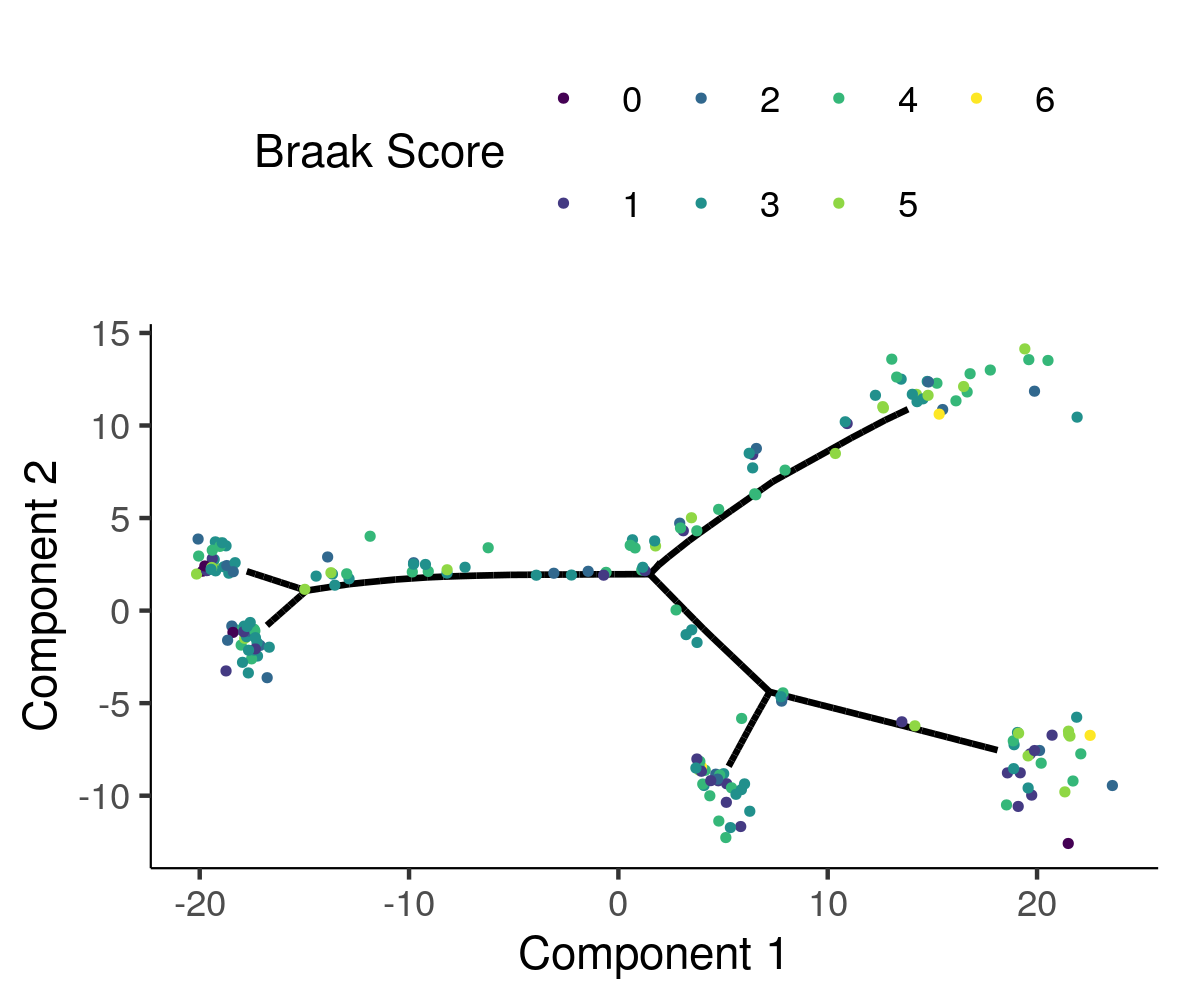

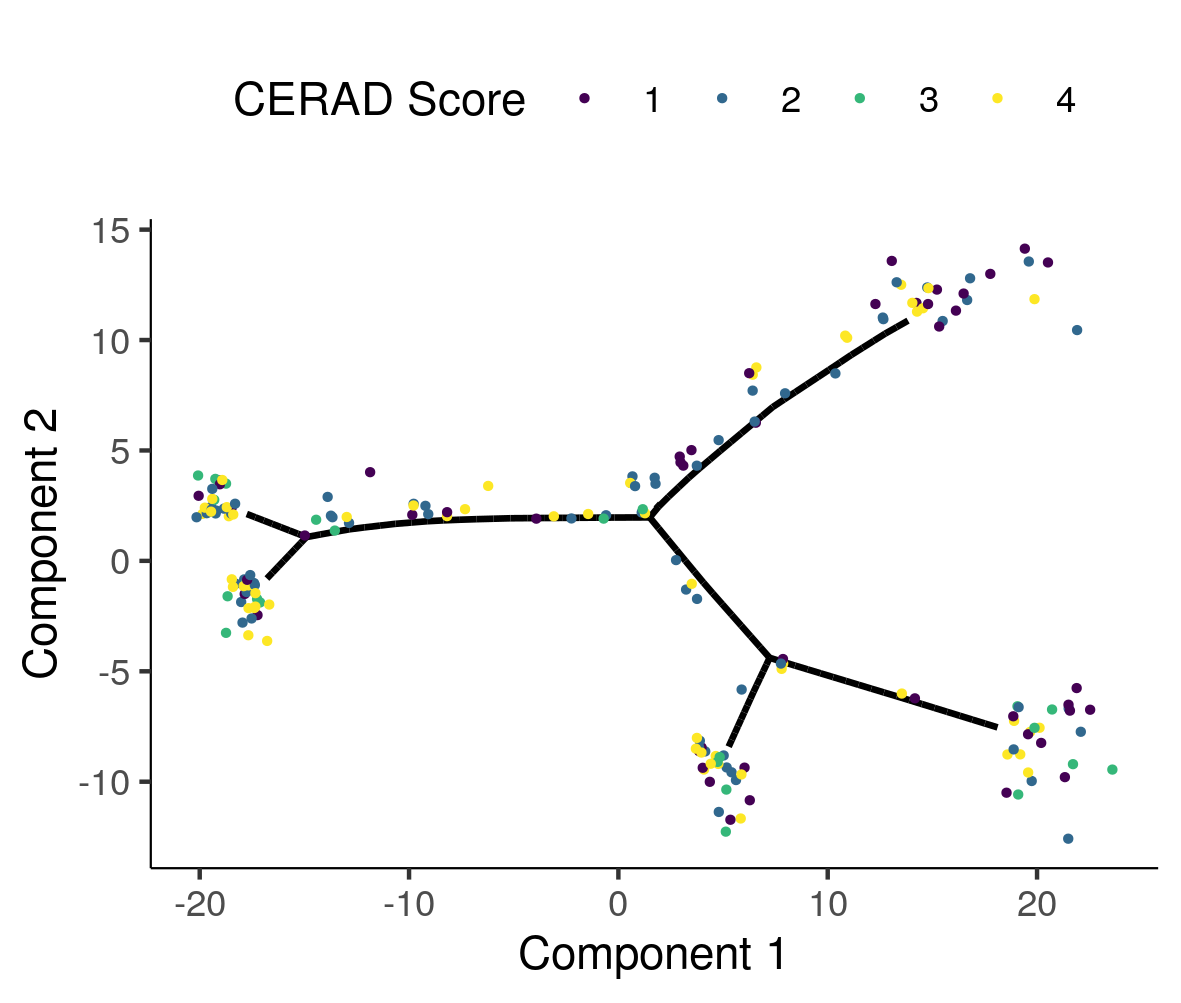

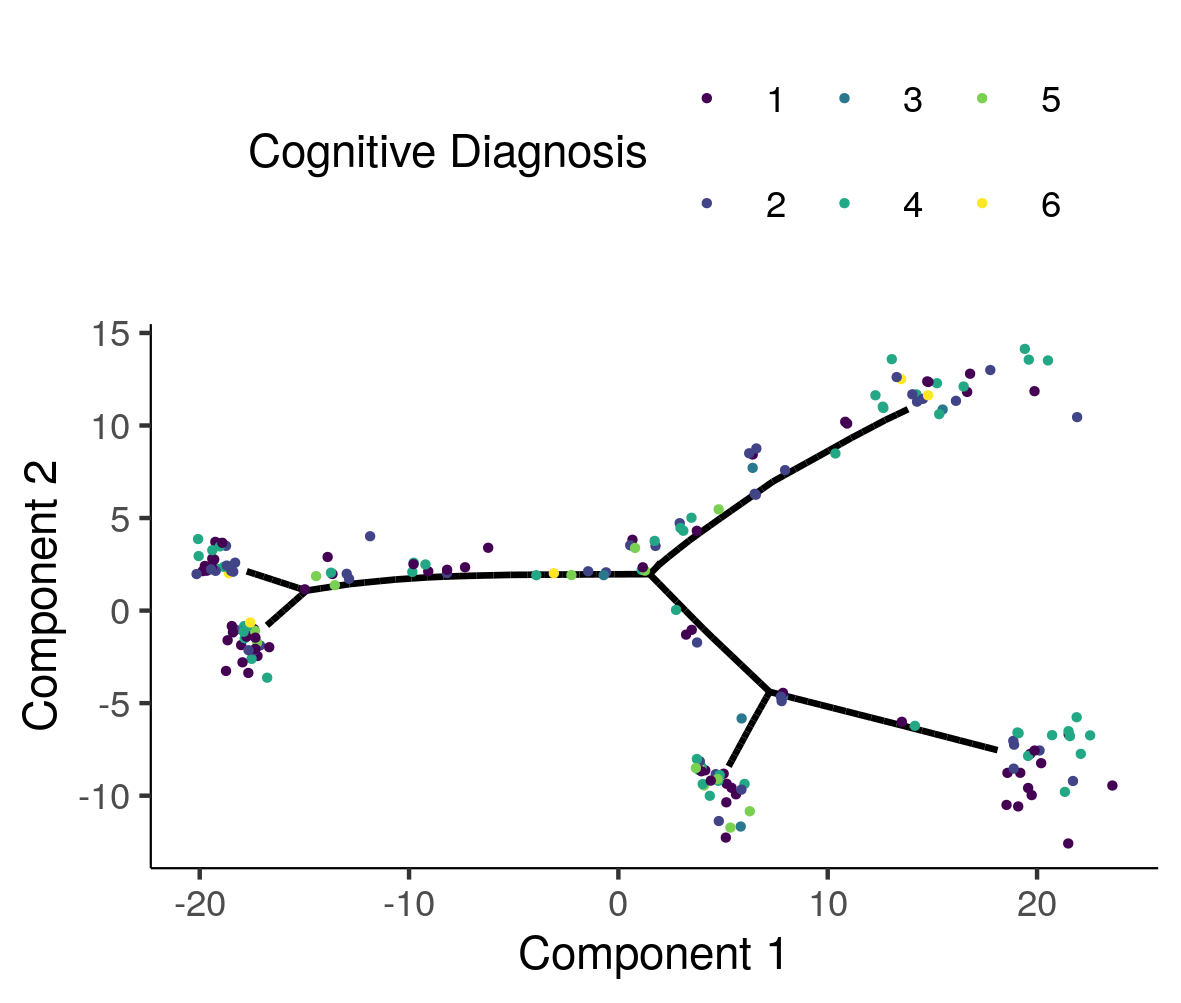

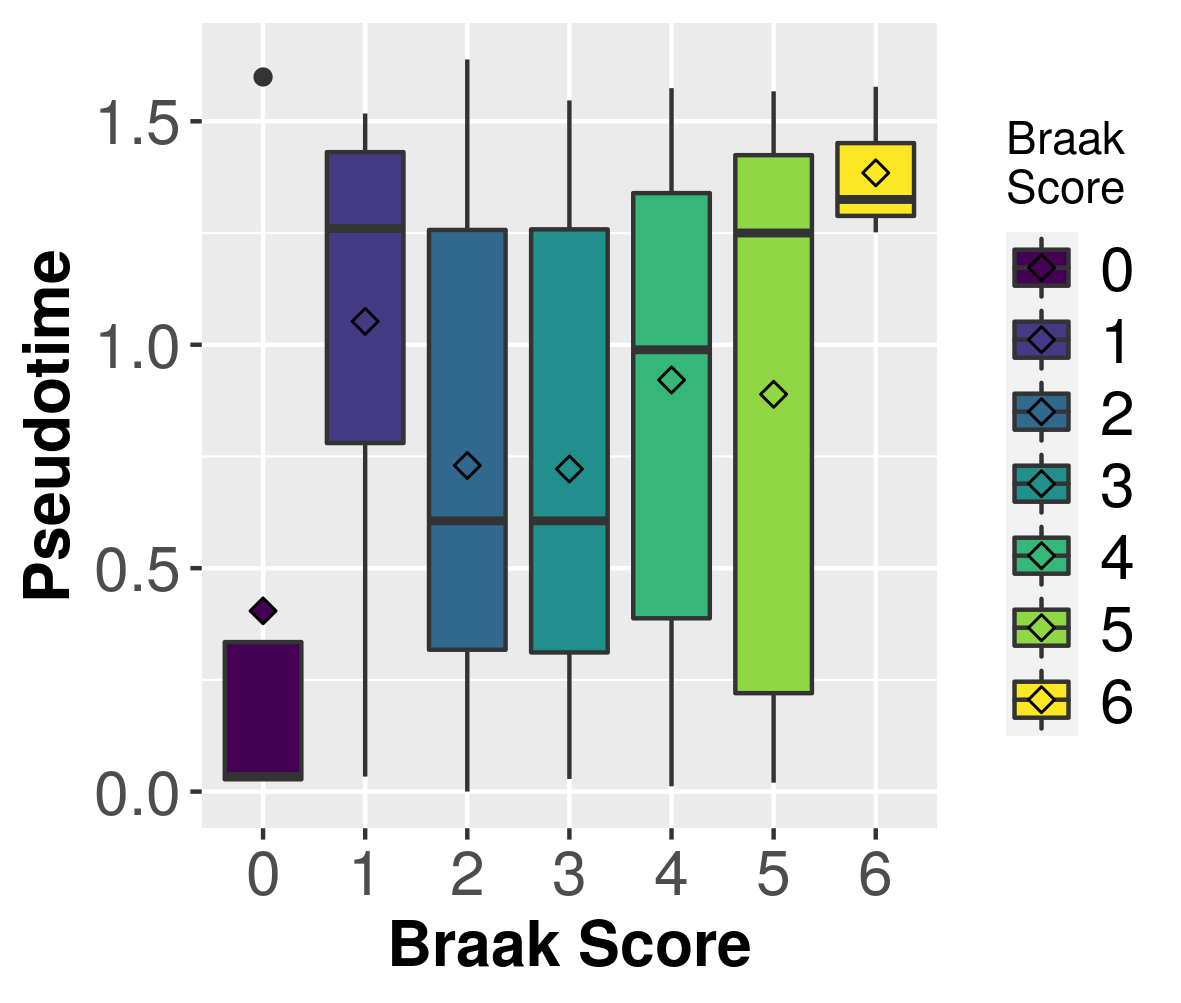

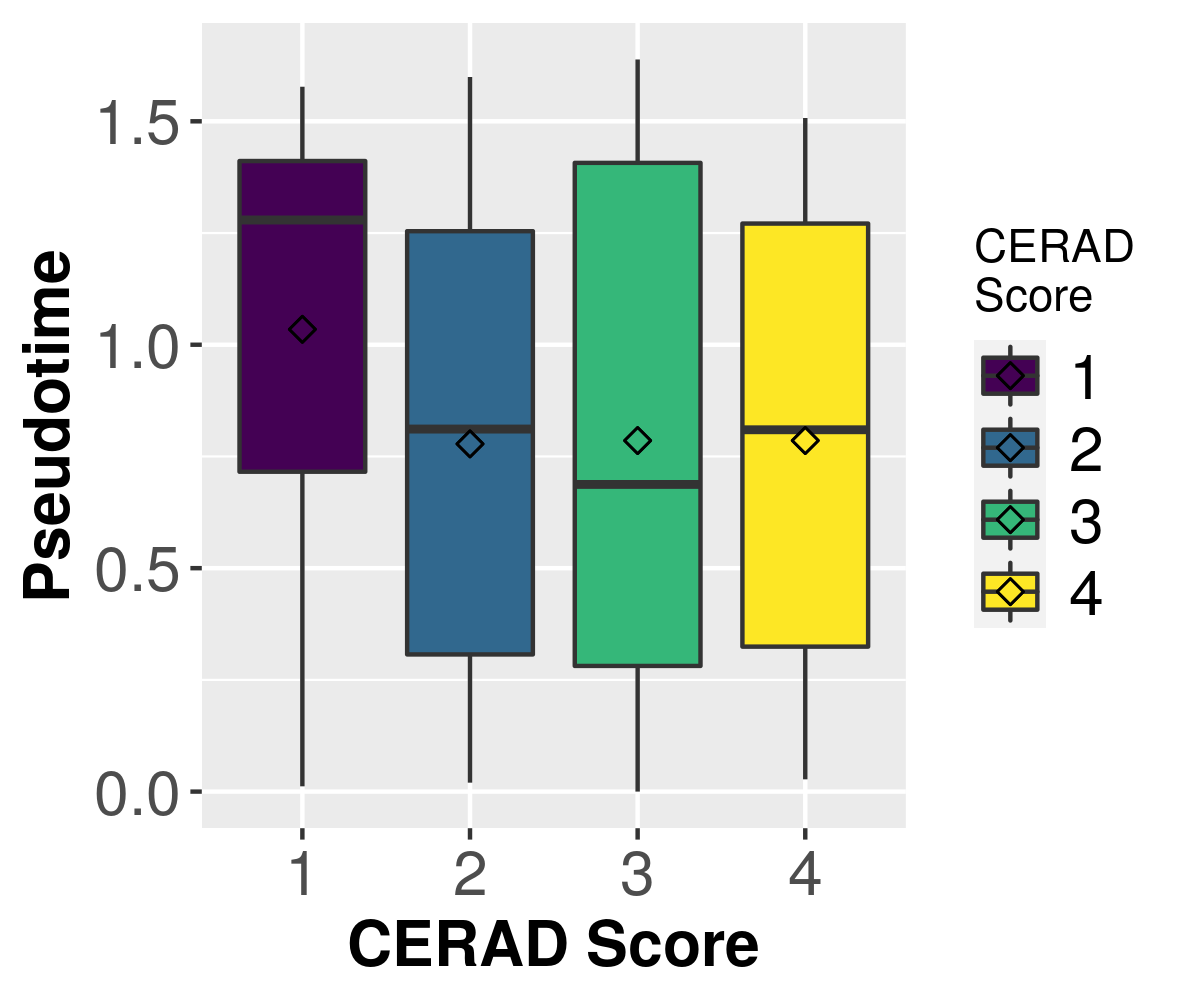

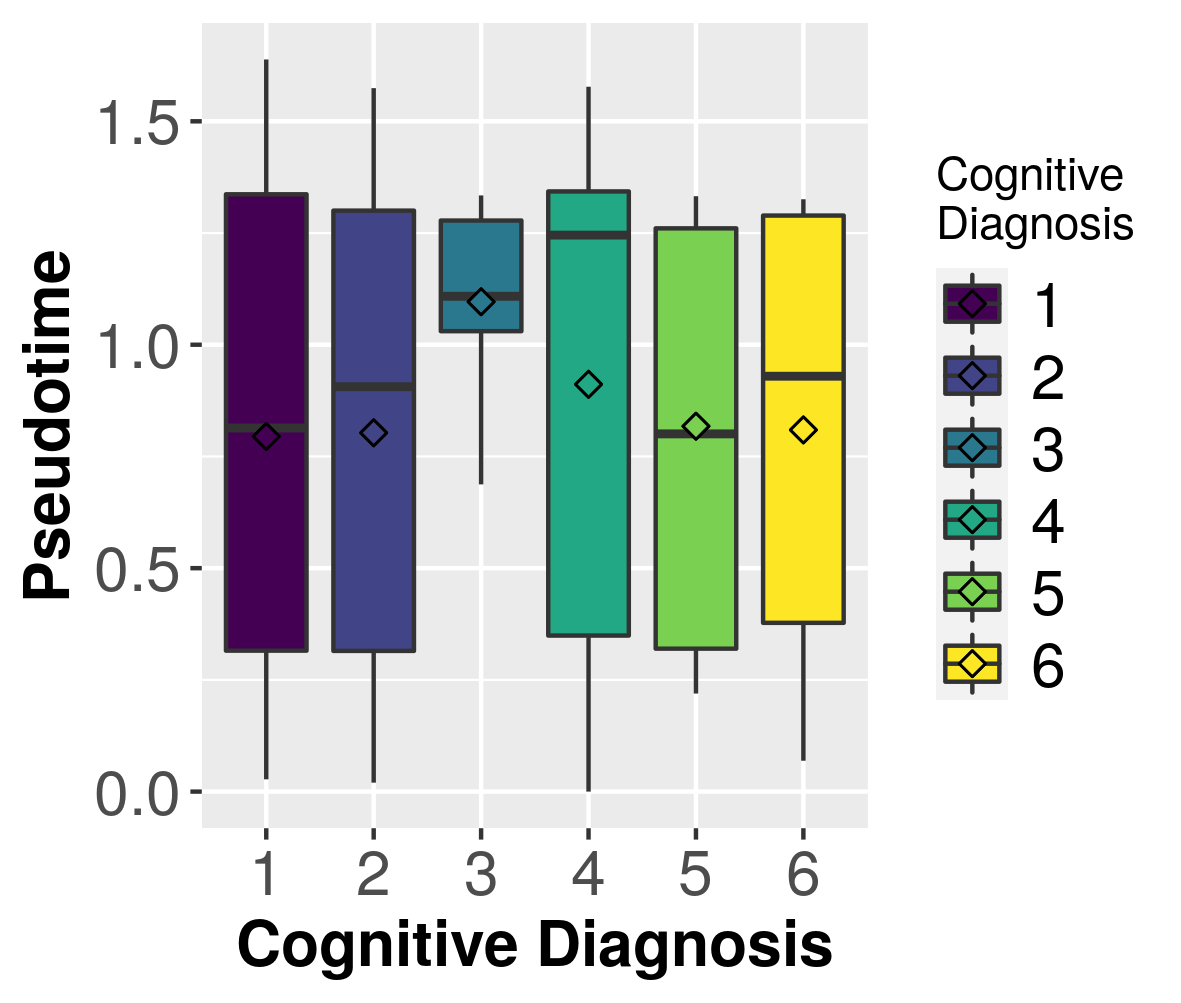

**Supplementary Figure 10 –** Manifold learning infers disease states from RNA-seq samples, samples from males and females combined for 361 independent samples from two independent studies. A) Estimated cell trajectory from A) TCX and B) DLPFC. C) Distribution of pseudotime for AD cases and controls for DLPFC and TCX. D) Distribution of expression correlation with pseudotime for both LOAD GWAS genes and non-LOAD GWAS genes. Box plots have lower and upper hinges at the 25th and 75th percentiles and whiskers extending to at most 1.5xIQR (interquartile range).

**Supplementary Figure 11 –** Manifold learning and measures of staging in LOAD in DLPFC samples, male and females combined for 537 independent samples from one study. A-C) Samples colored by three external measures of LOAD staging: Braak score, CERAD score, and cognitive diagnosis. D-F) Distribution of samples by inferred stage for different distinct stages in each of the three methods of measuring LOAD severity. Box plots have lower and upper hinges at the 25th and 75th percentiles and whiskers extending to at most 1.5xIQR (interquartile range).

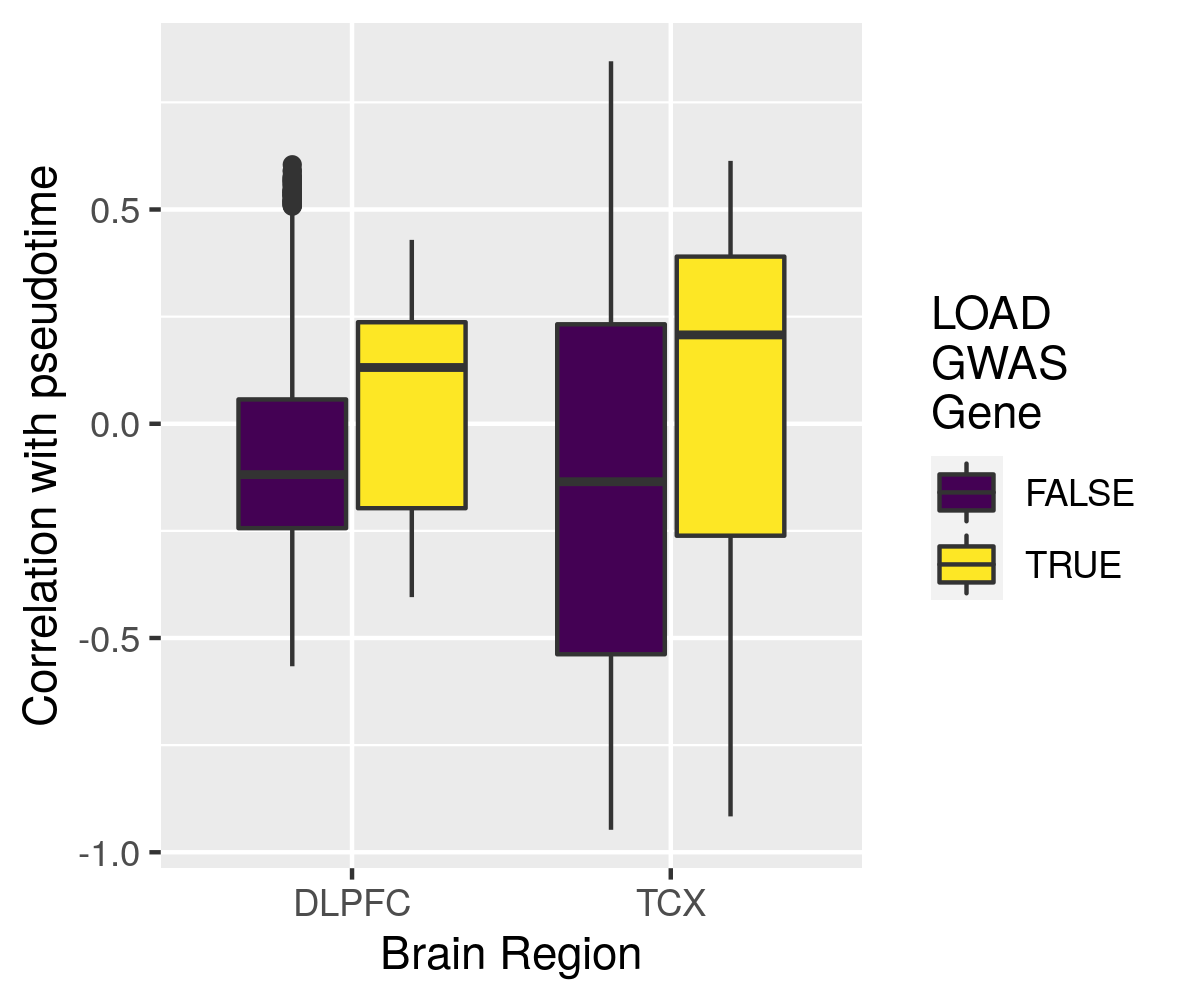

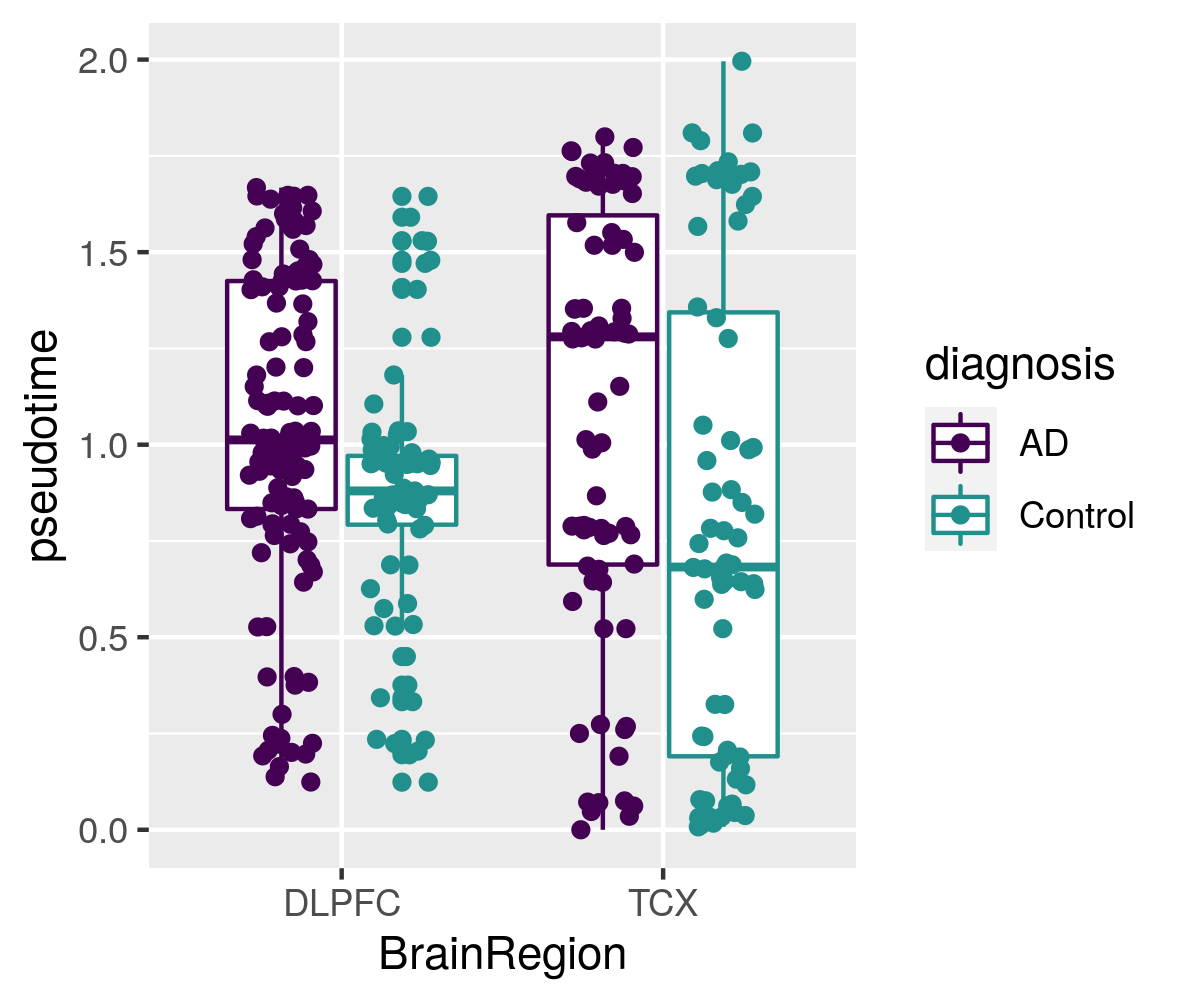

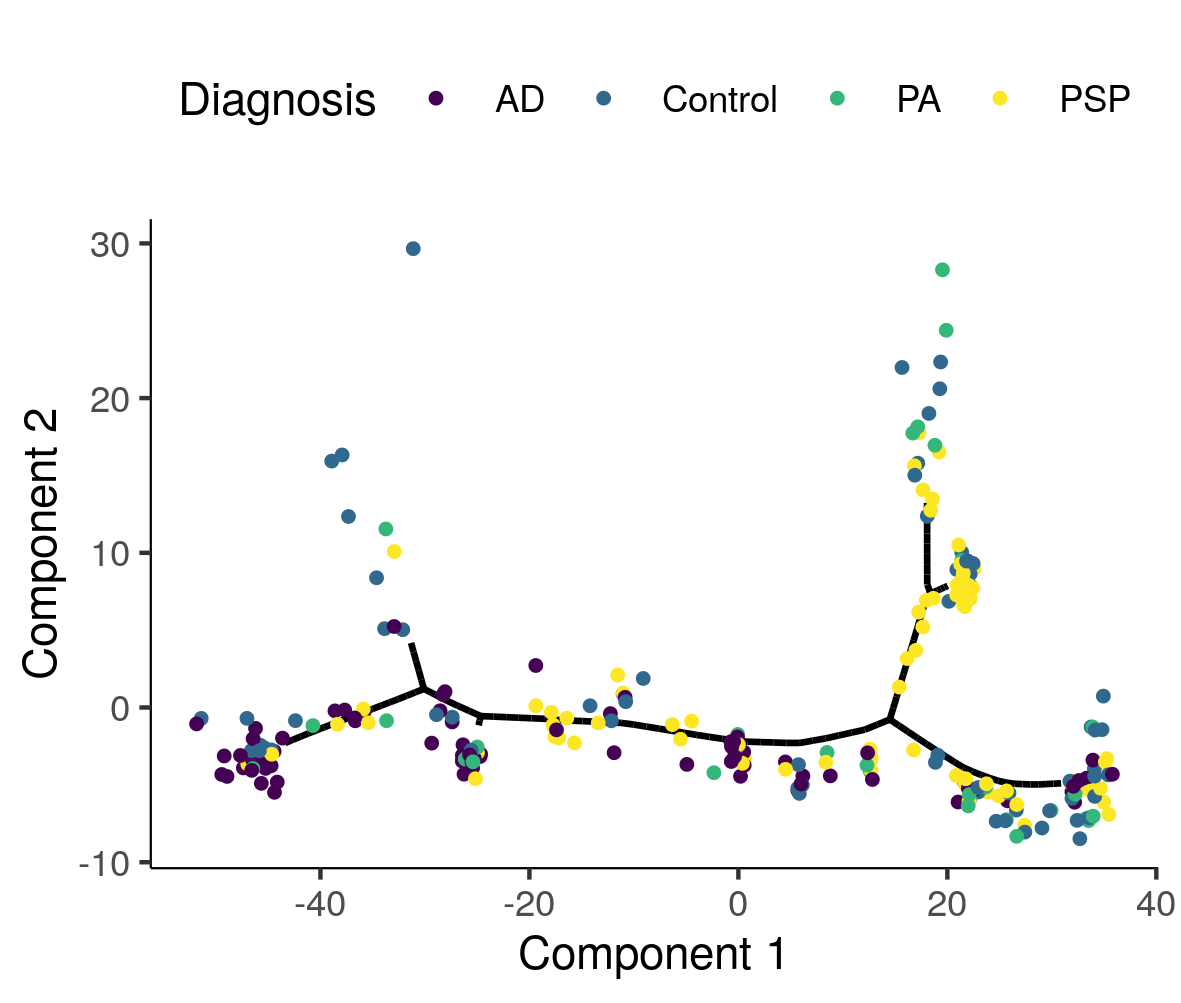

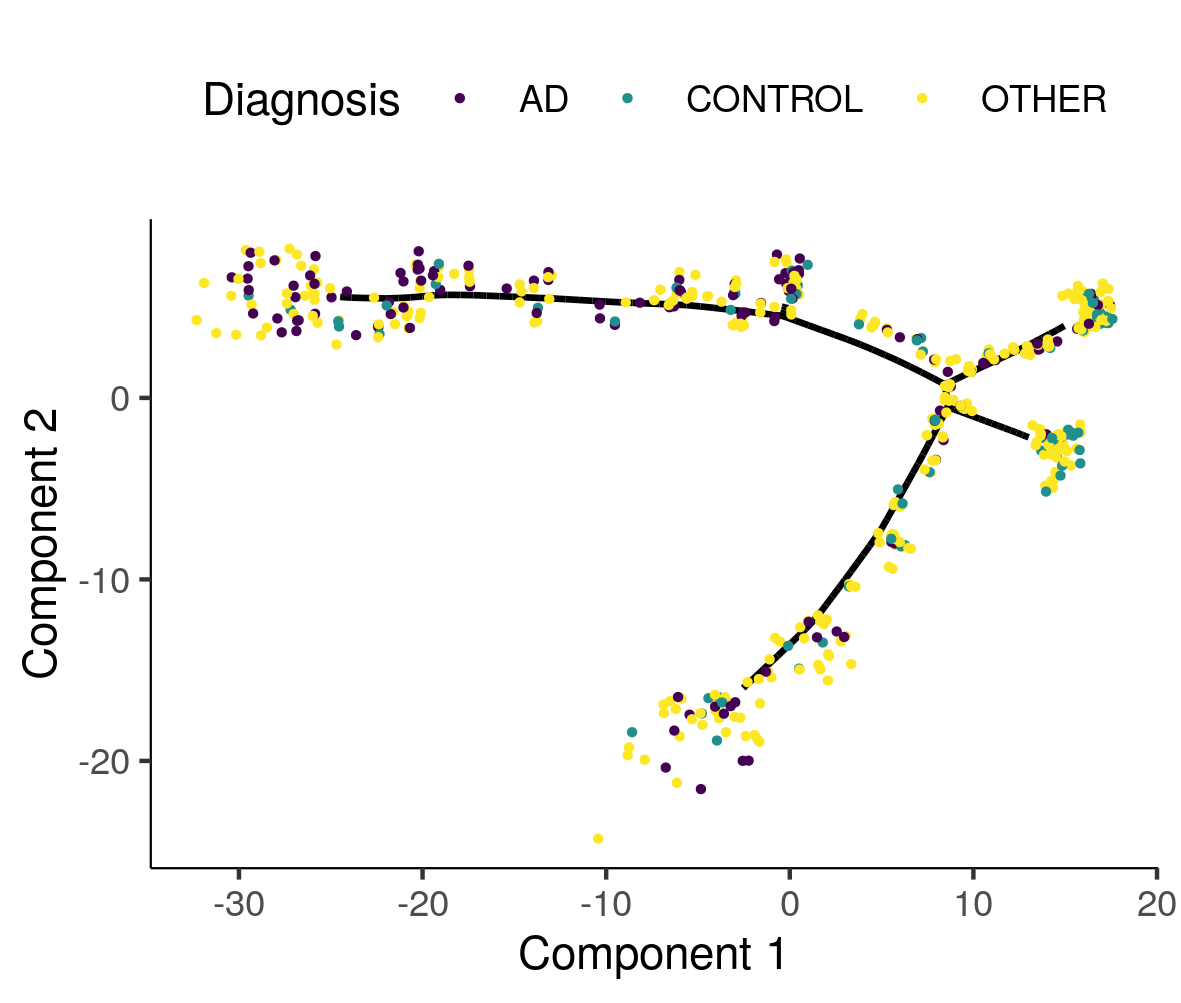

A

D

C

B

**Supplementary Figure 12 -** Correlations with pseudotime for IGAP GWAS genes for 17446 genes from two independent studies for A) PMI adjusted pseudotimes, B) top 10 PC adjusted pseudotime, C) RIN adjusted pseudotimes, and D) PMI, PC, and RIN adjusted pseudotimes. Box plots have lower and upper hinges at the 25th and 75th percentiles and whiskers extending to at most 1.5xIQR (interquartile range).

C

B

A

D

E

F

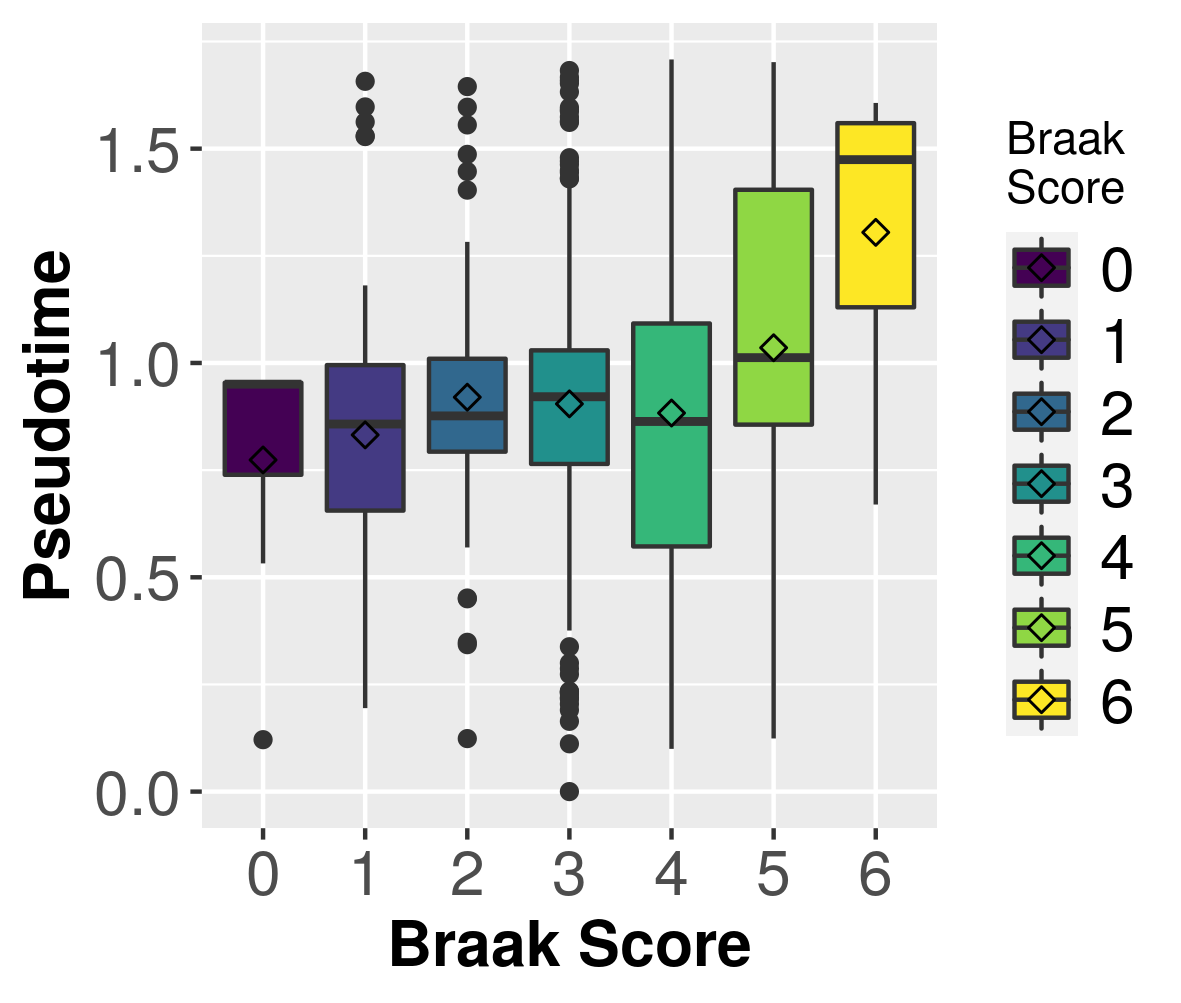

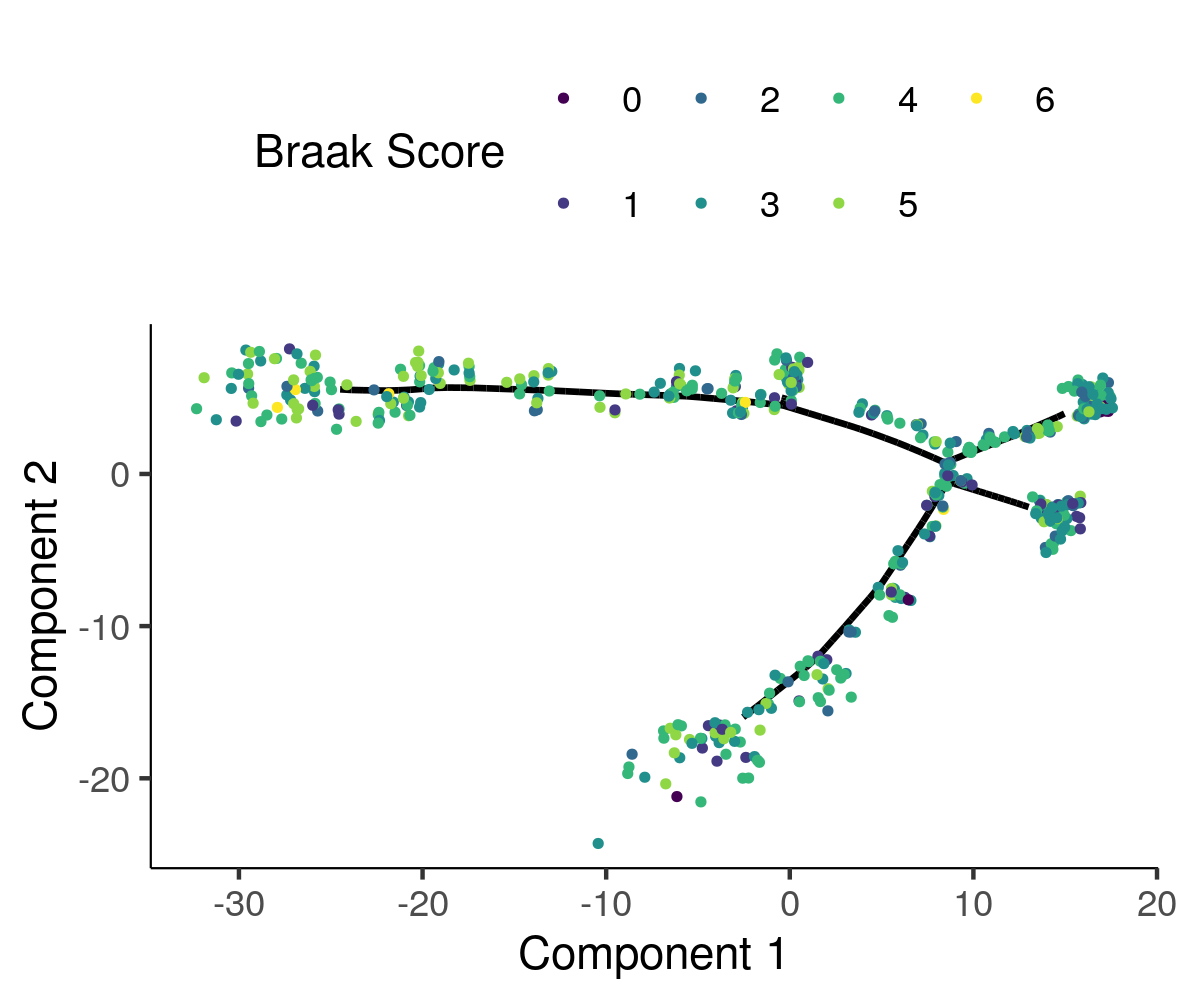

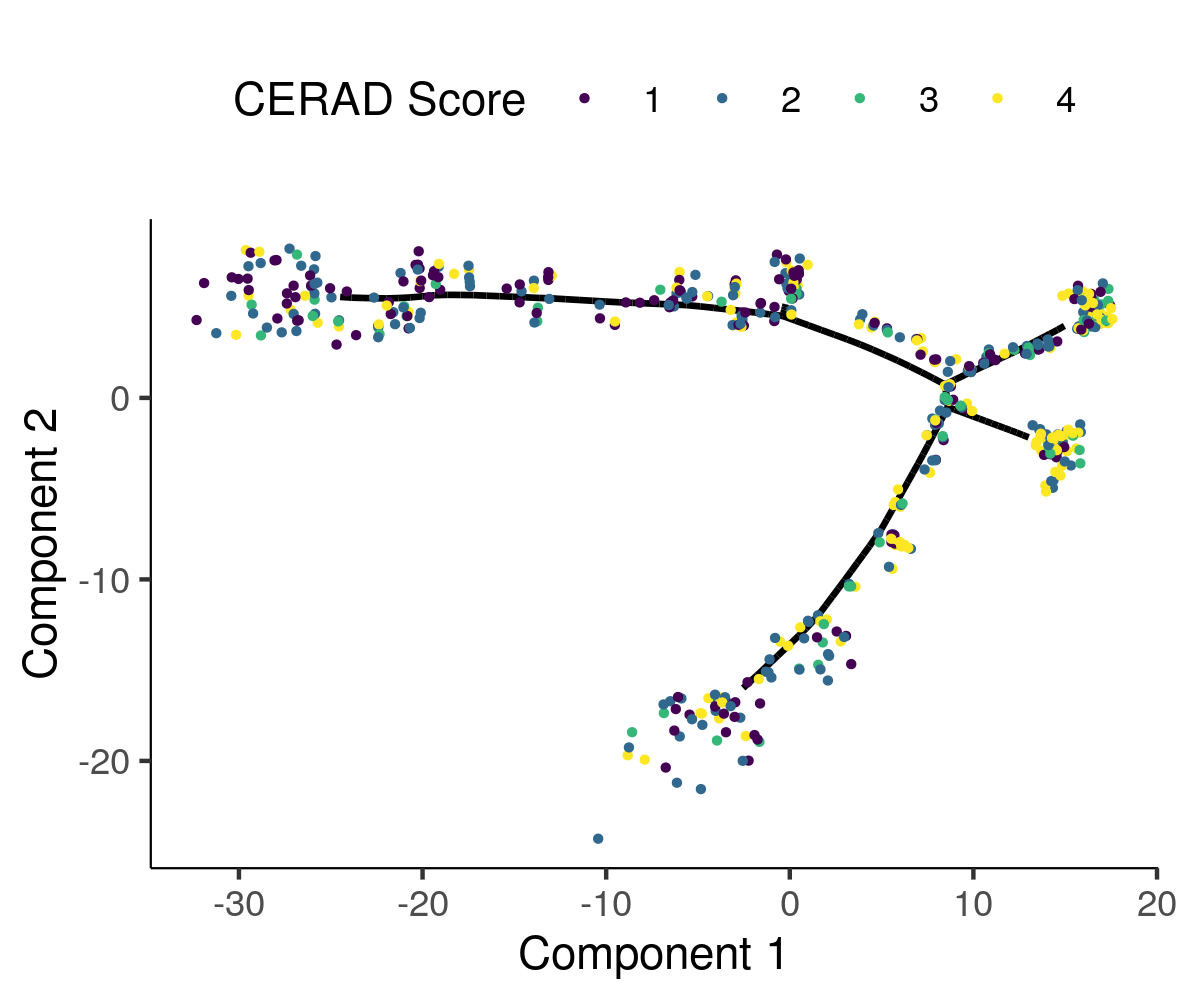

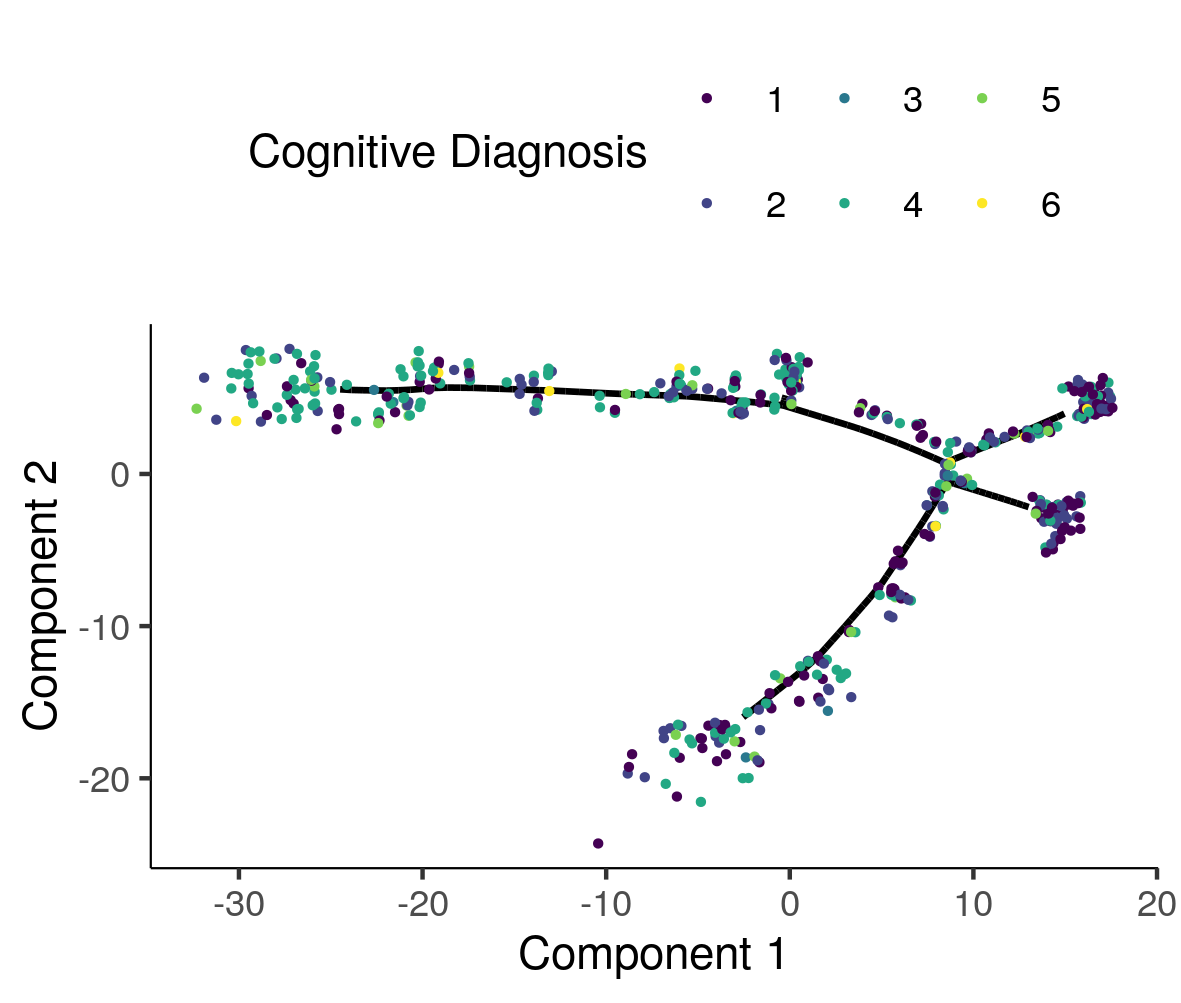

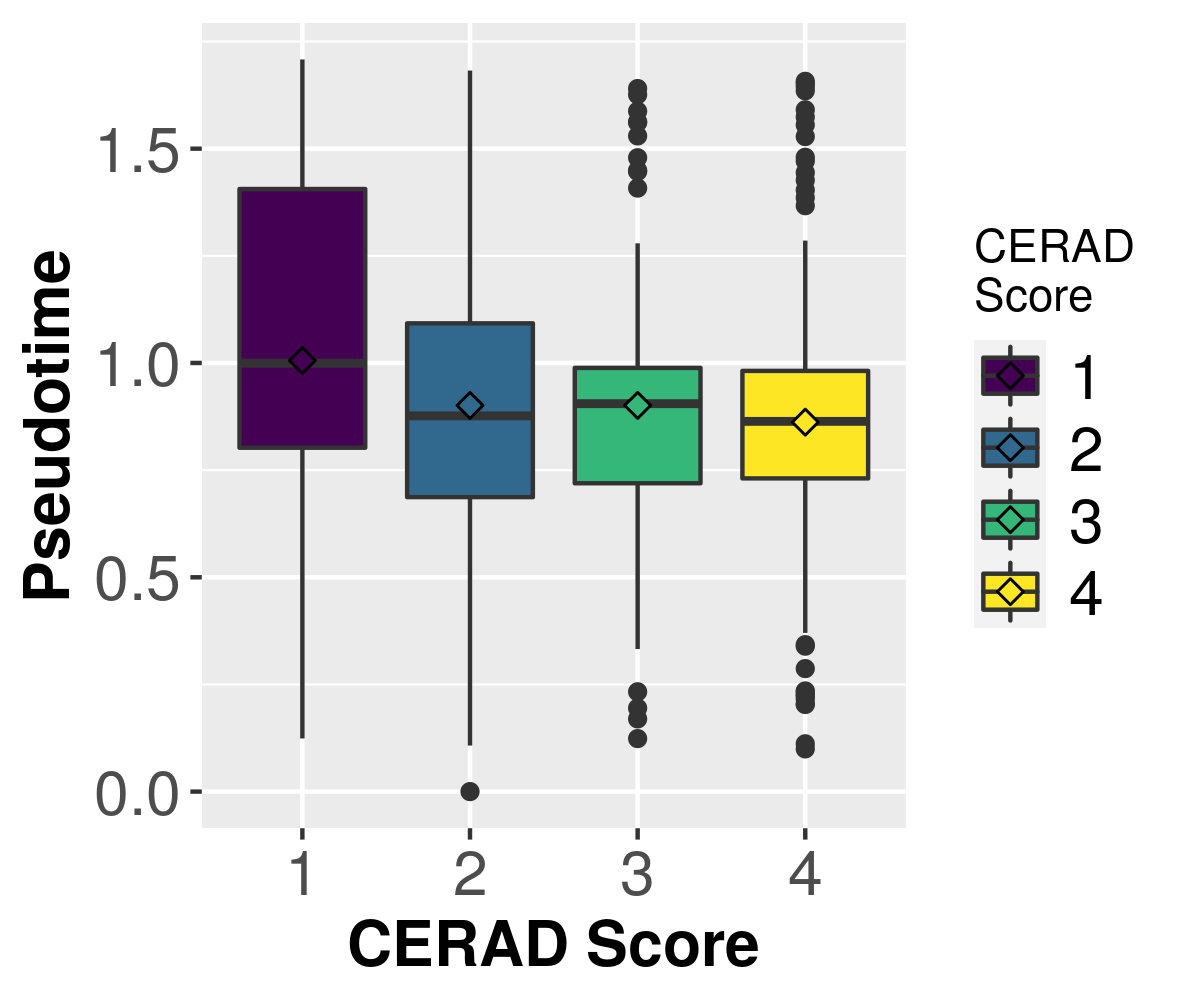

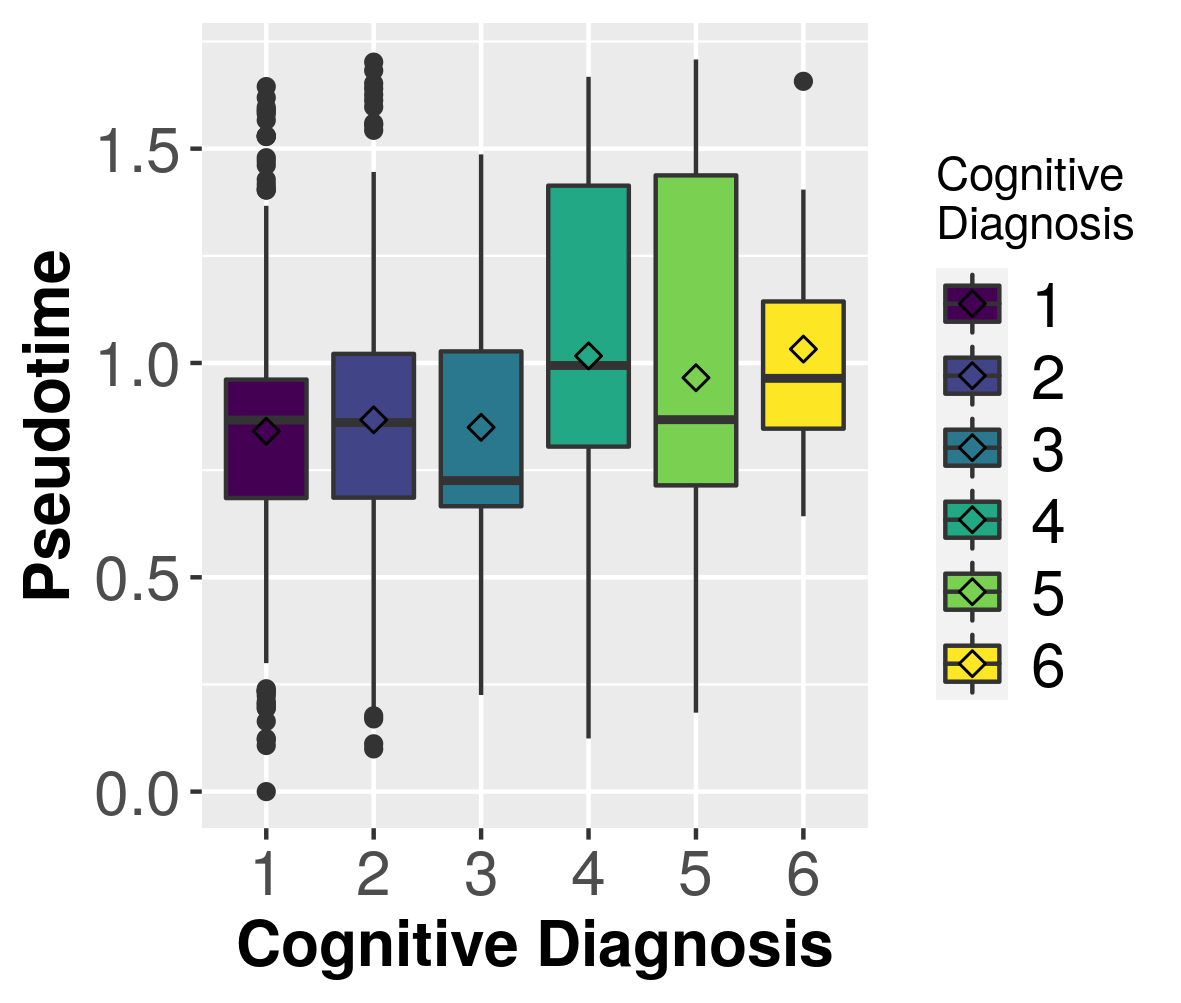

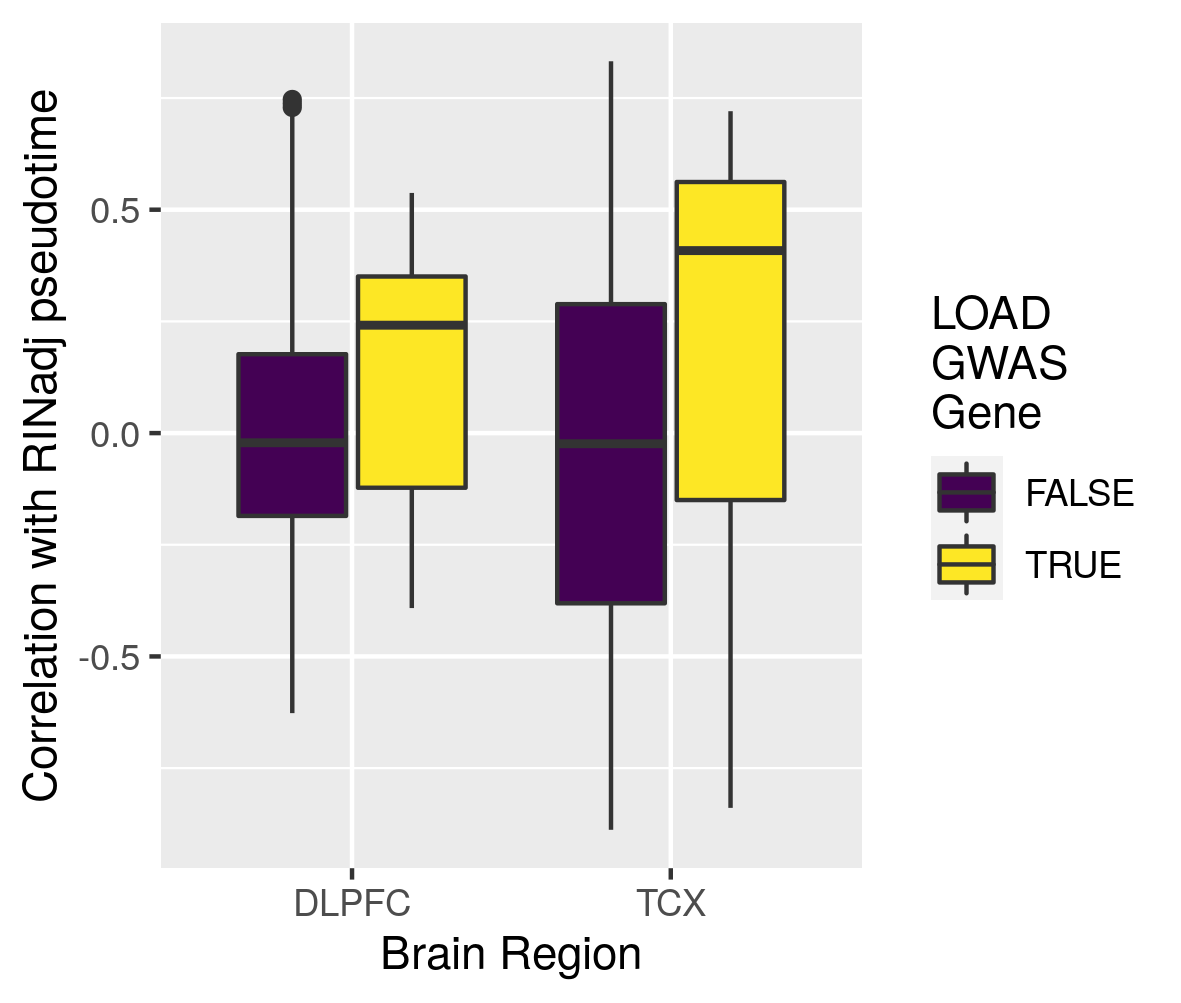

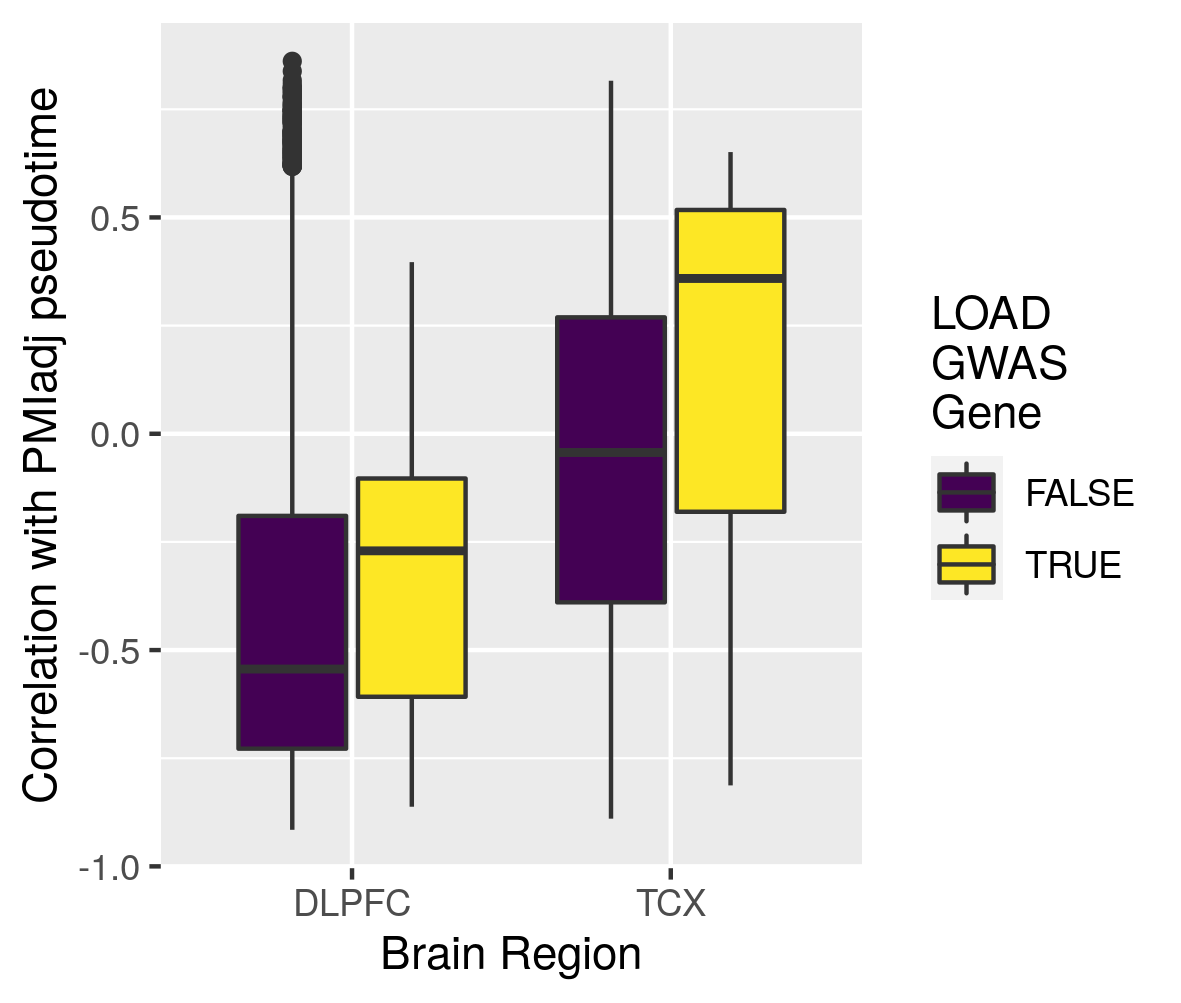

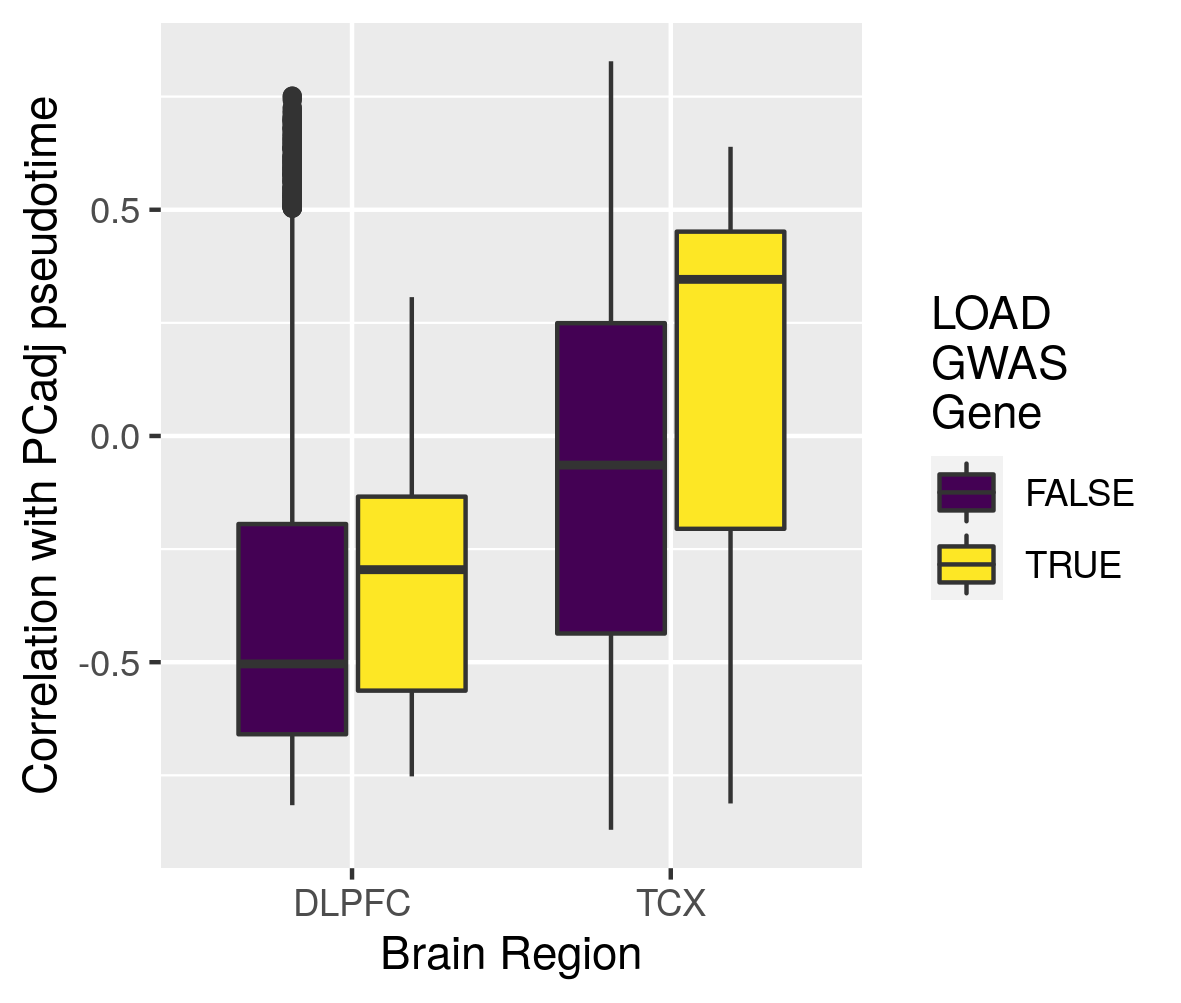

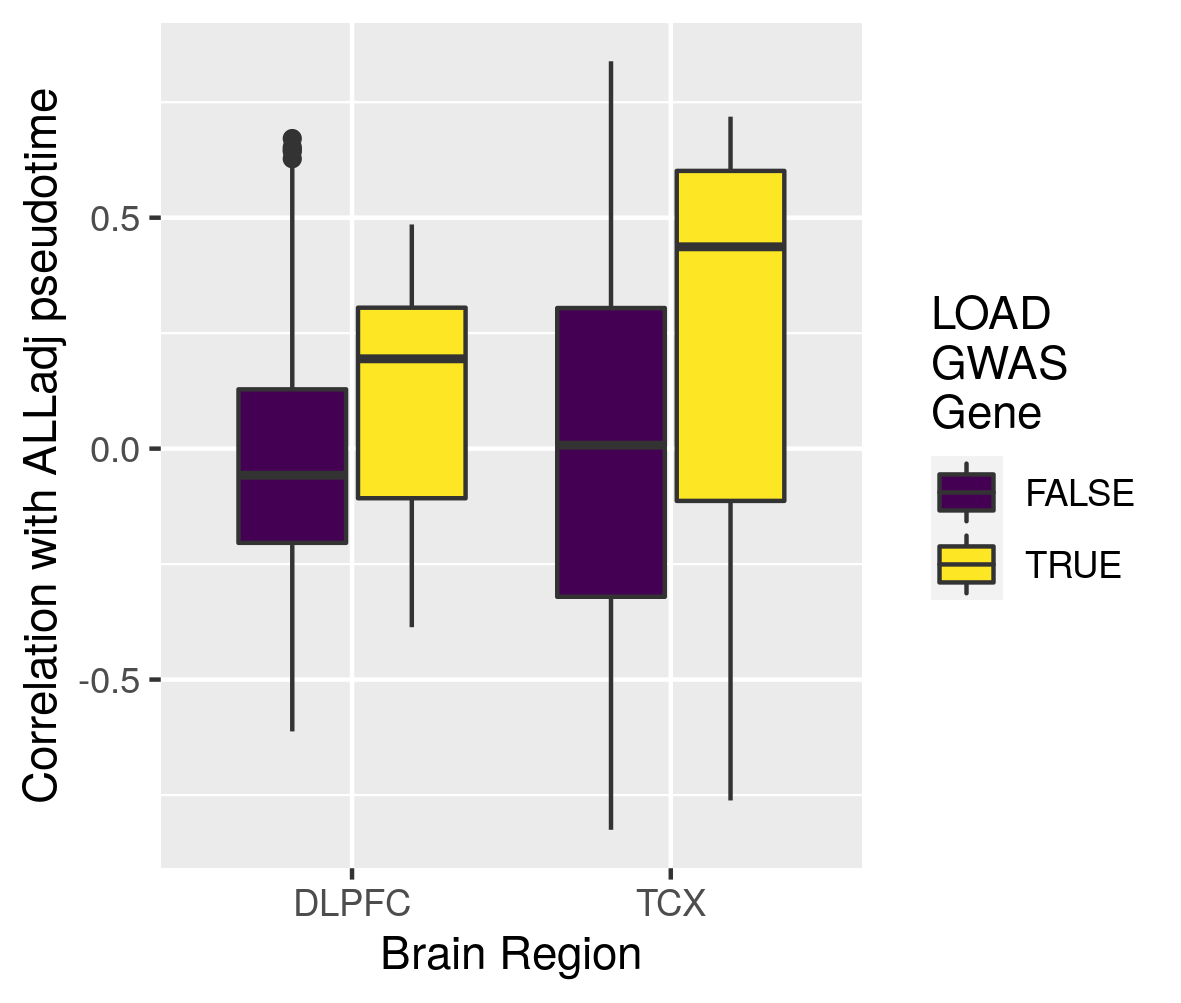

A

D

C

B

**Supplementary Figure 13 –** Lineages analyses adjusted for Braak score in DLPFC for 338 independent samples from one study. A) Lineage adjusted for Braak, B) Diagnosis as a function of Braak adjusted pseudotime, C) Cognitive diagnosis on Braak adjusted lineage, D) Cognitive diagnosis as a function of Braak adjusted pseudotime, E) correlation between IGAP GWAS genes and Braak adjusted pseudotime for 17446 genes from one study. Box plots have lower and upper hinges at the 25th and 75th percentiles and whiskers extending to at most 1.5xIQR (interquartile range).

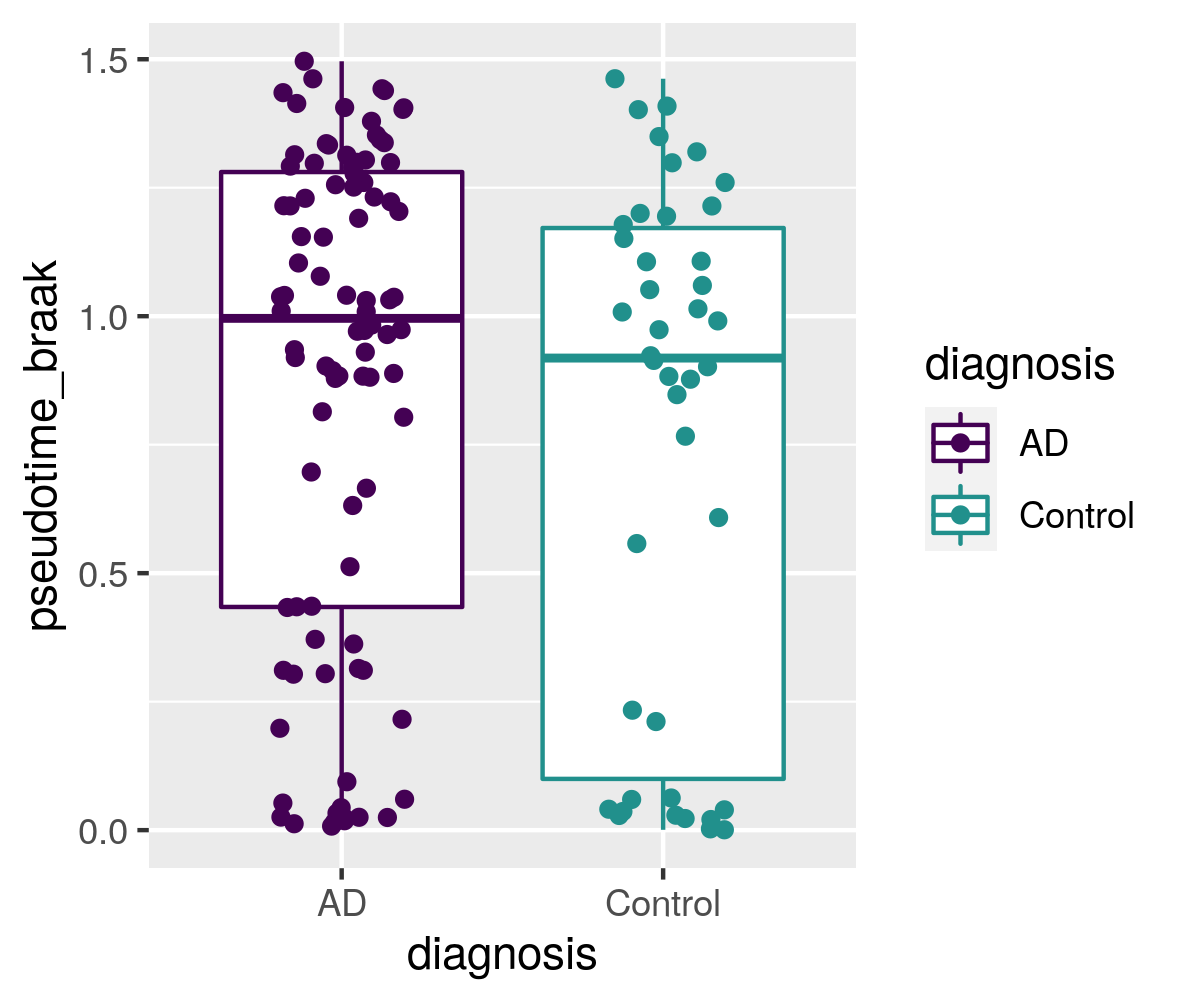

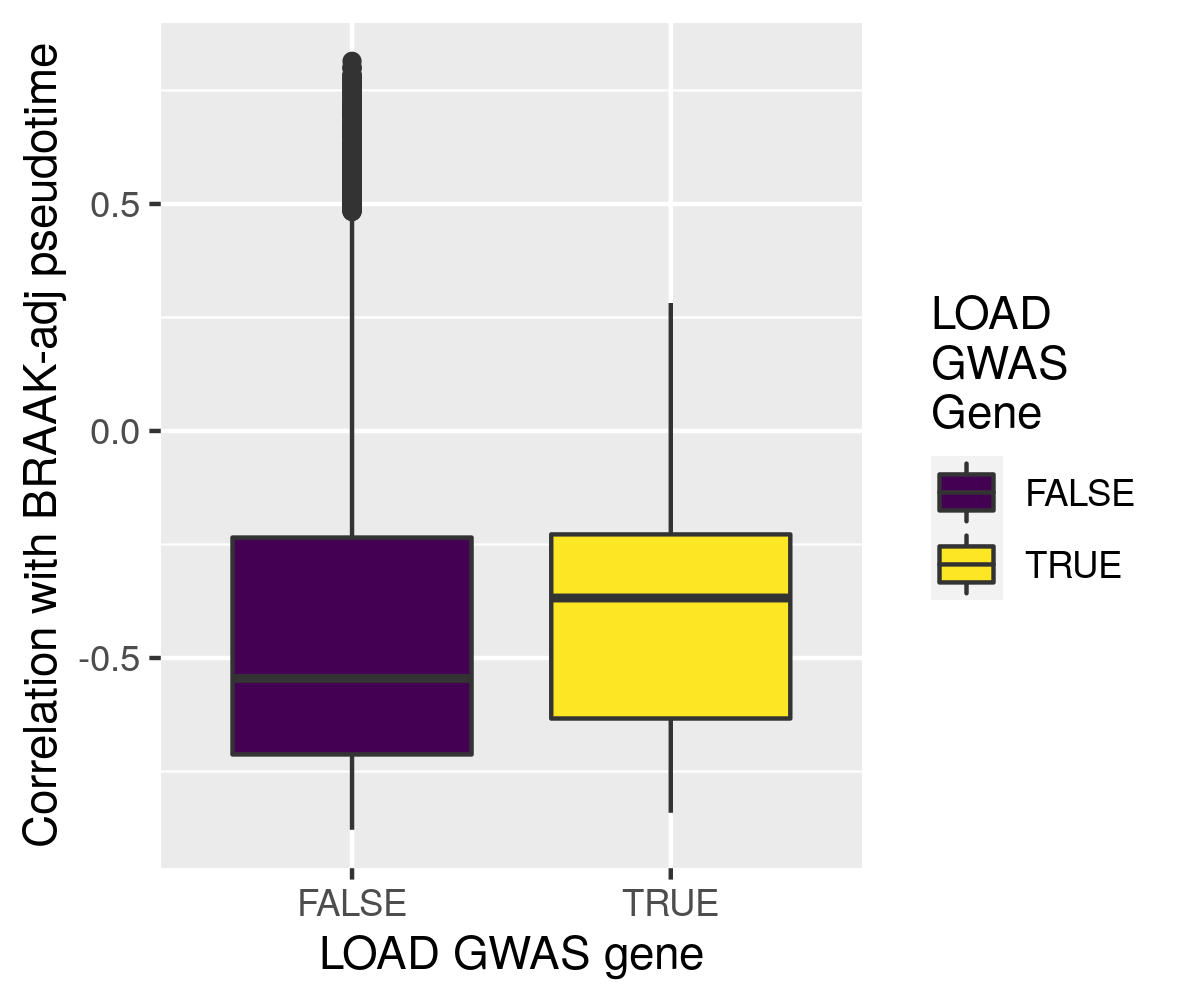

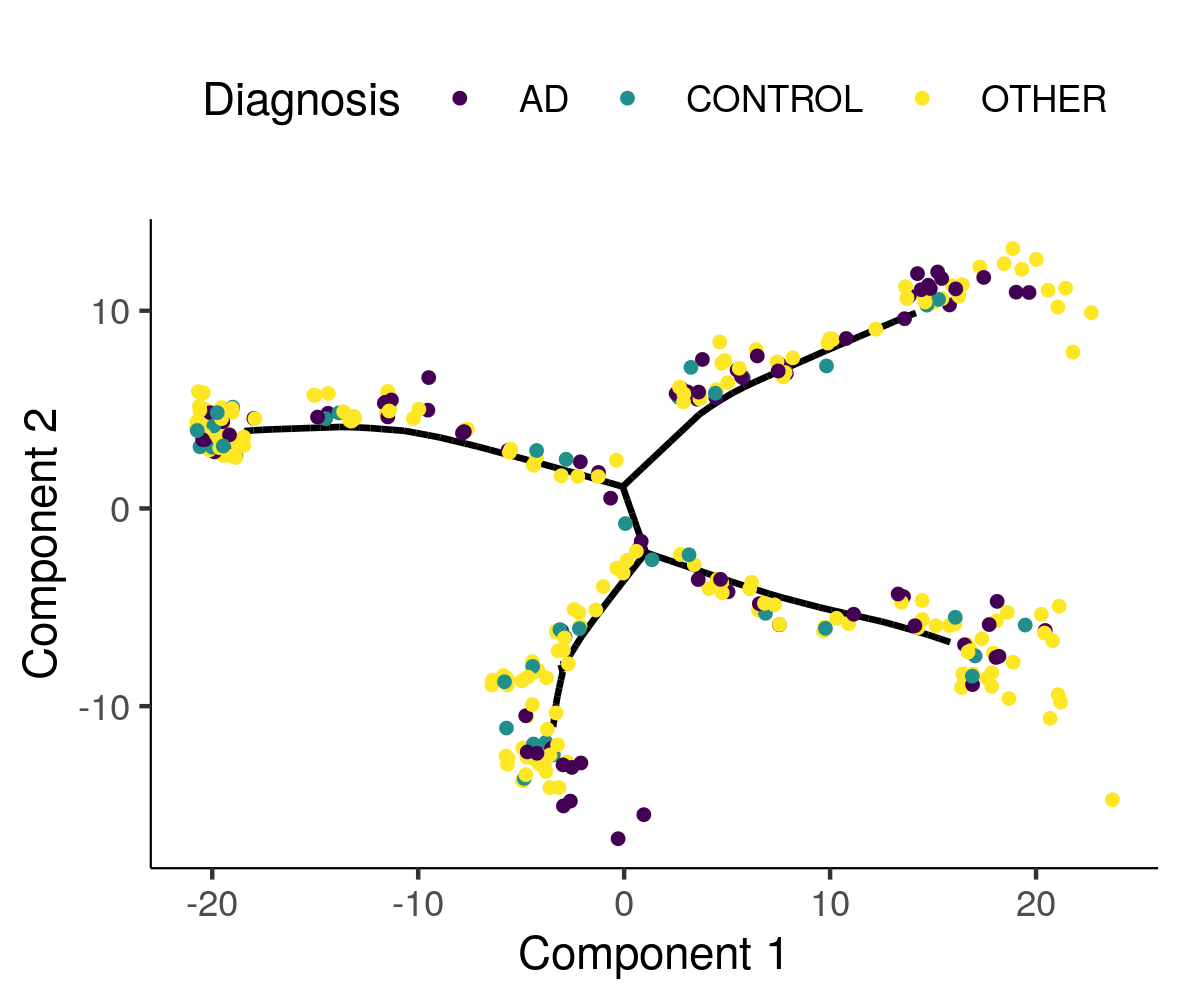

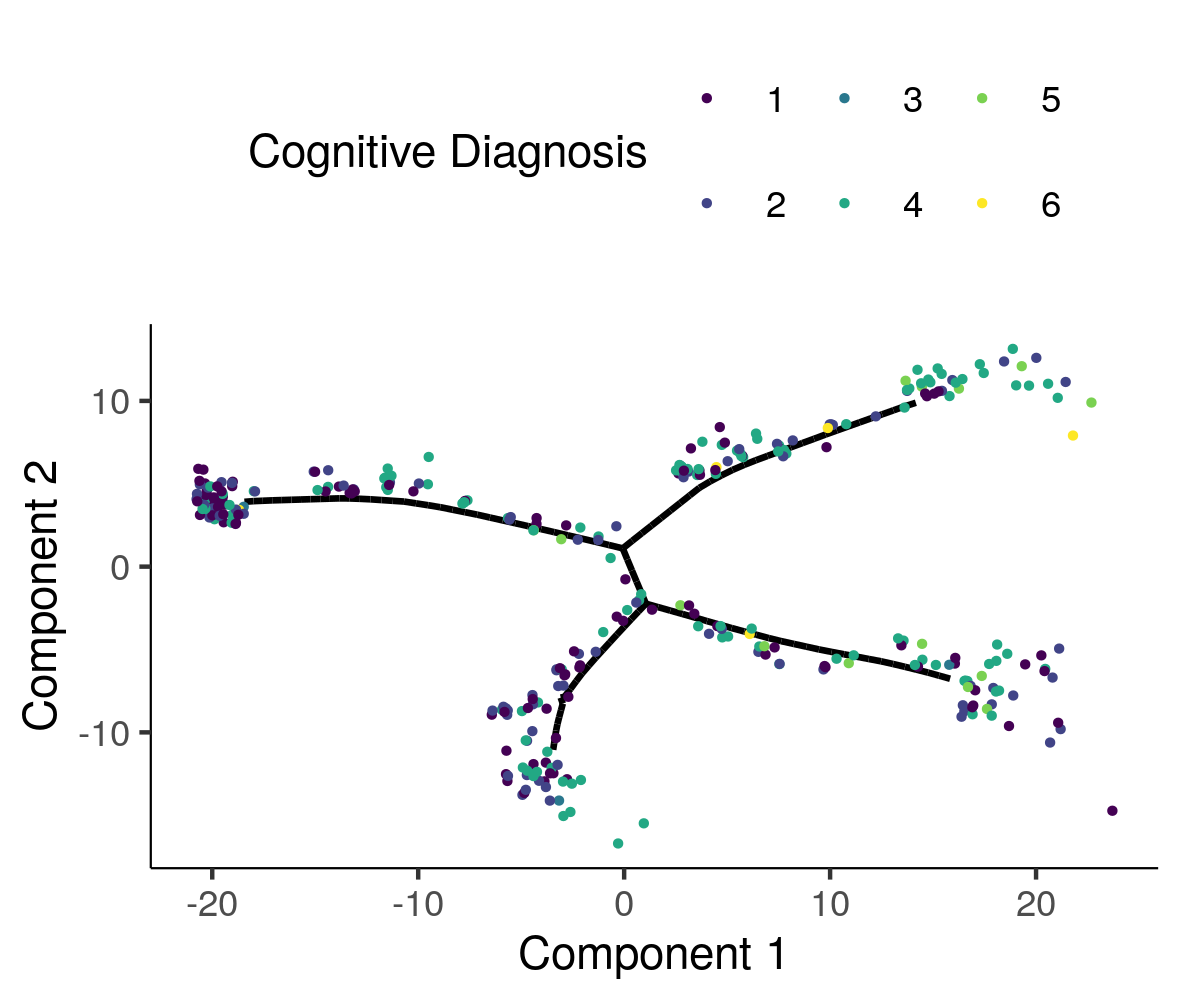

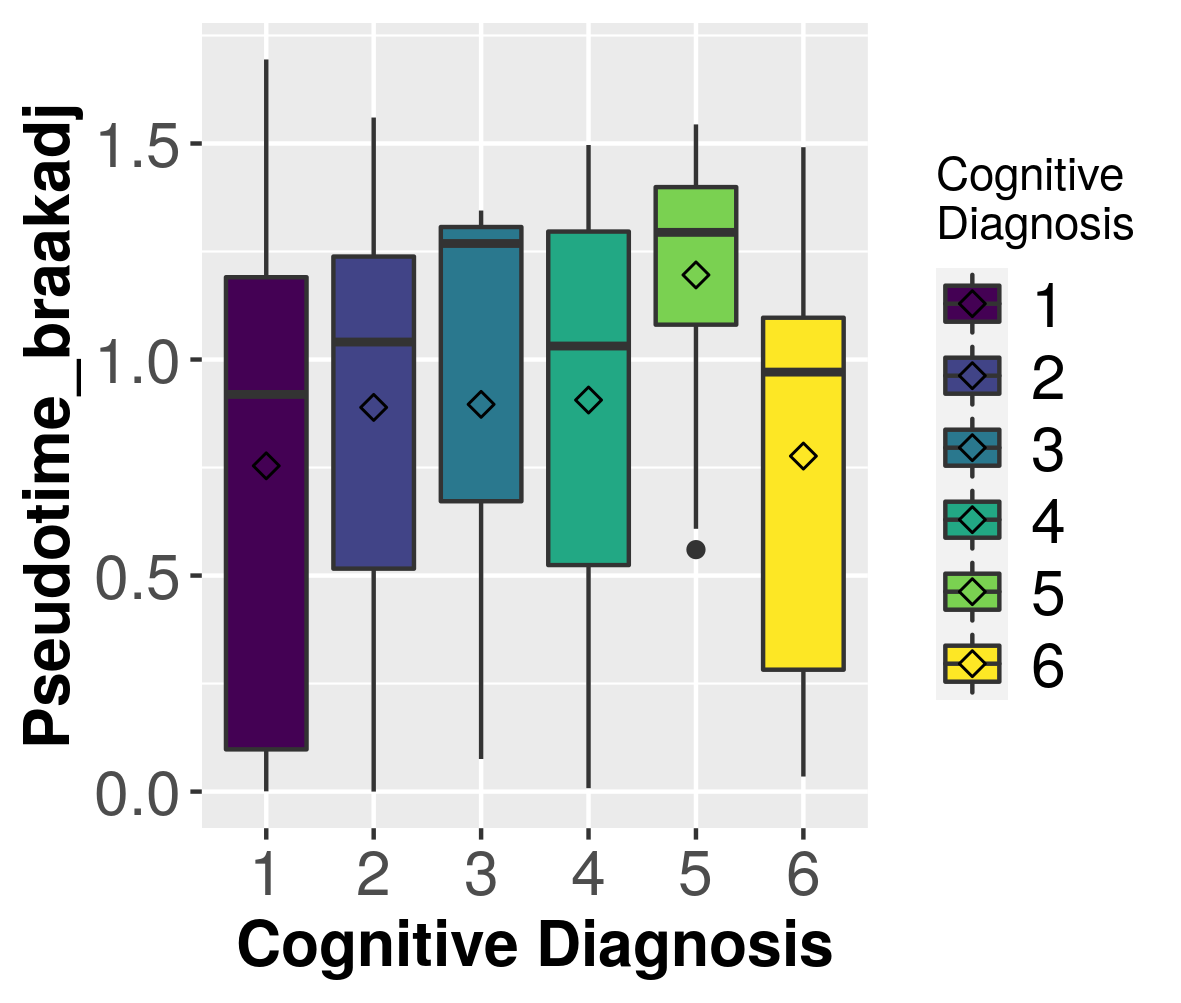

A

D

C

B

E

**Supplementary Figure 14 -** Manifold learning and measures of staging in LOAD in TCX samples for 76 independent samples from one study. A) Samples colored by two external measures of LOAD staging: Braak score and Thal amyloid. B) Distribution of samples by inferred stage for different distinct stages in each of the two methods of measuring LOAD severity. Box plots have lower and upper hinges at the 25th and 75th percentiles and whiskers extending to at most 1.5xIQR (interquartile range).

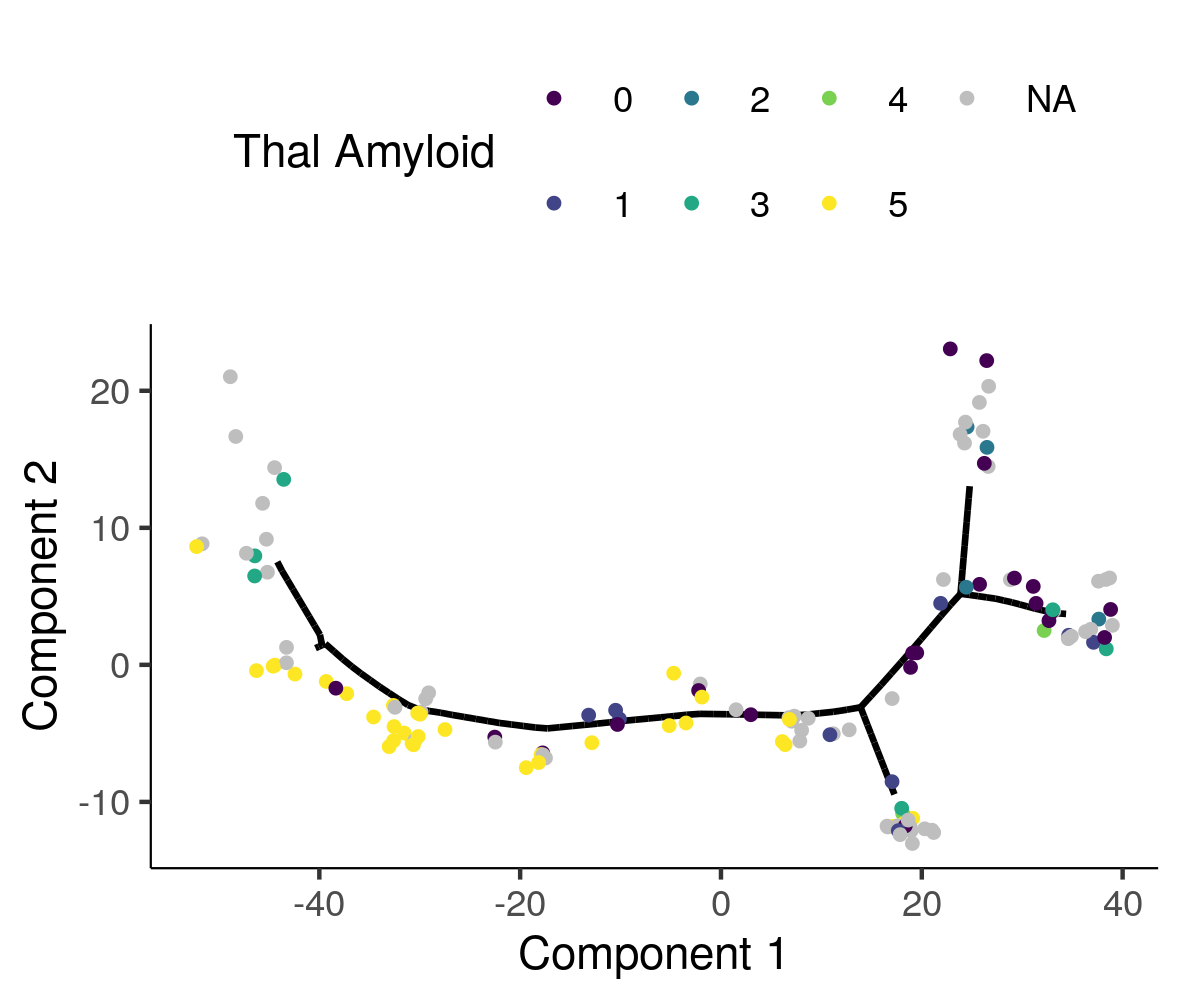

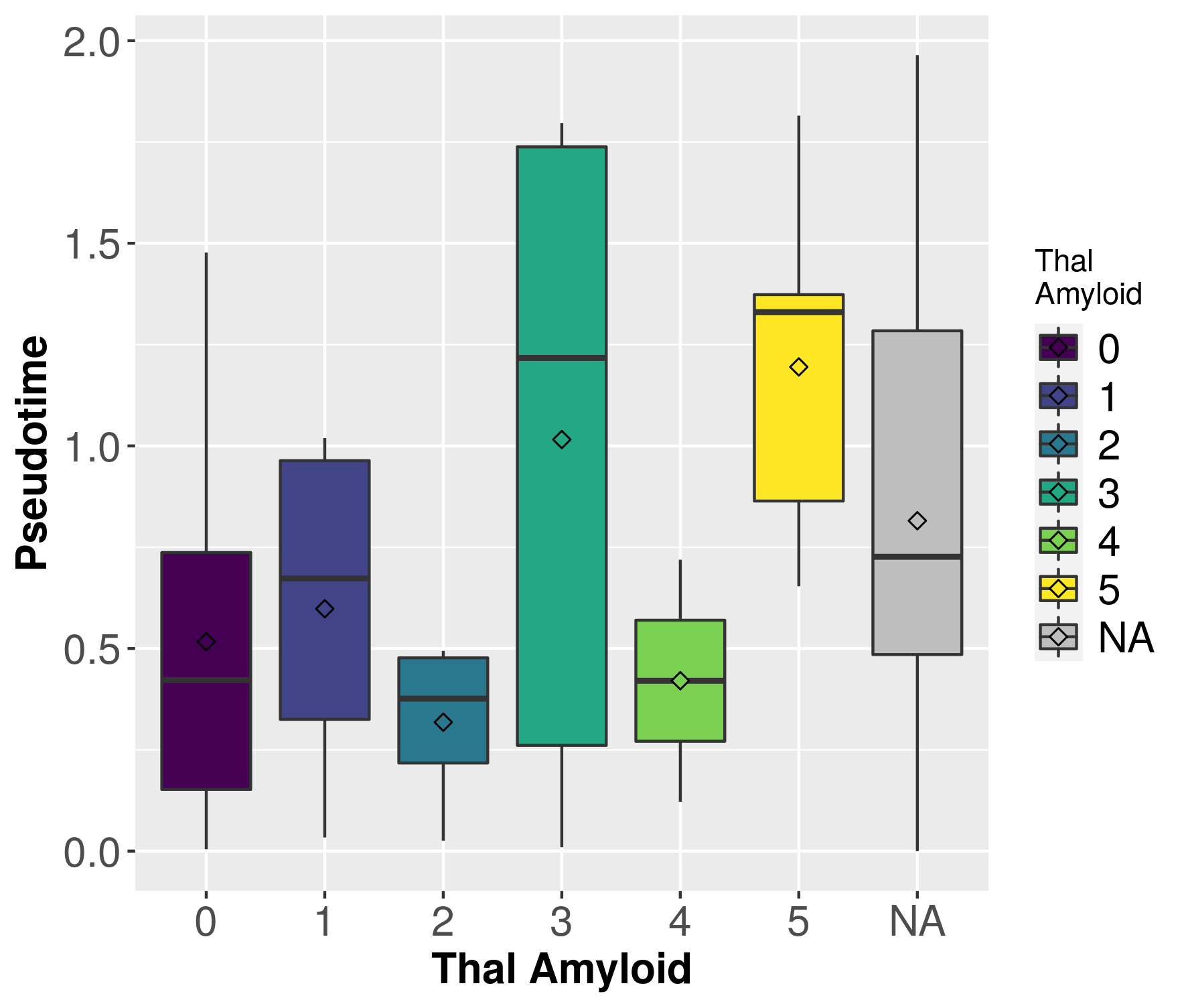

A

B

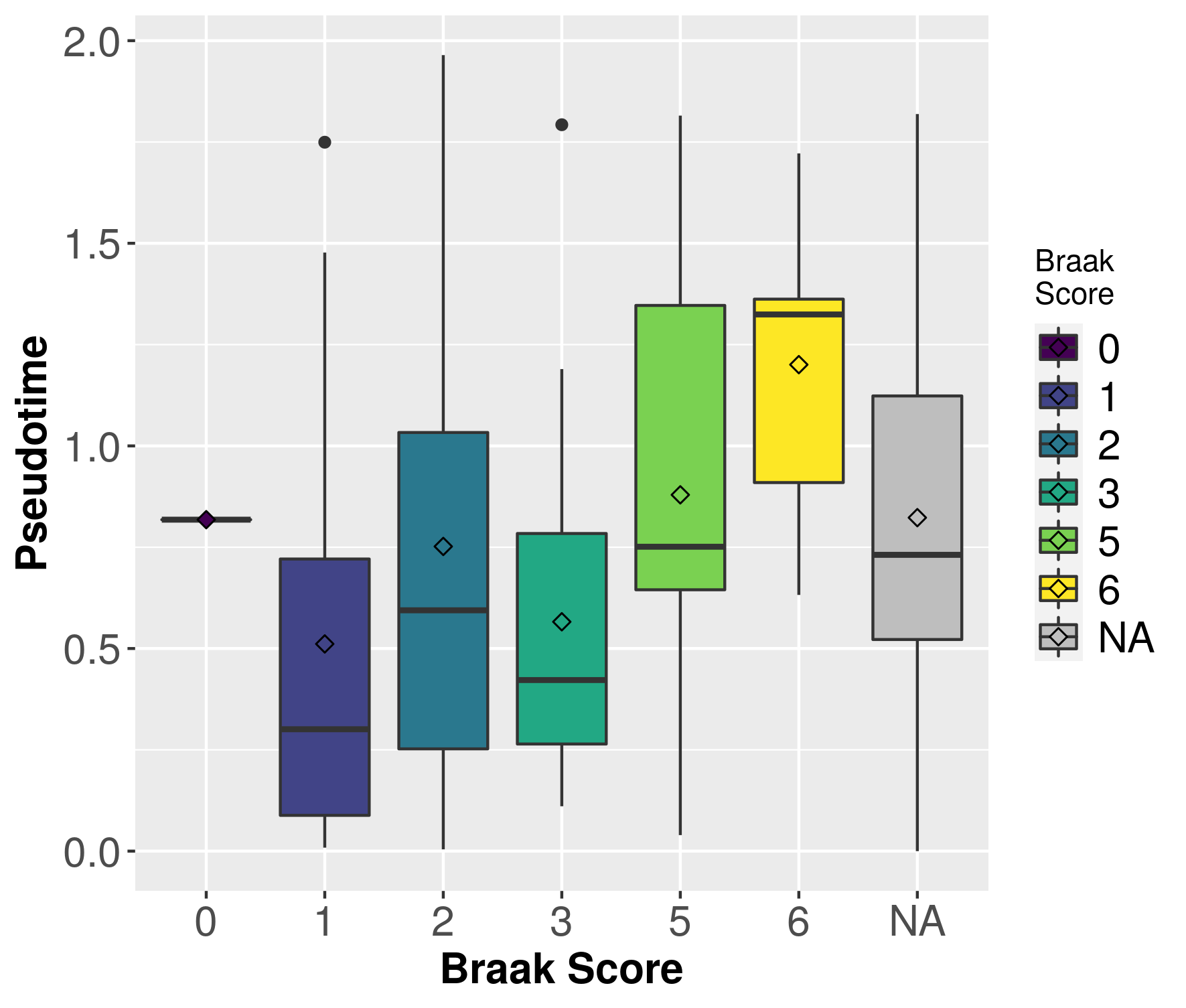

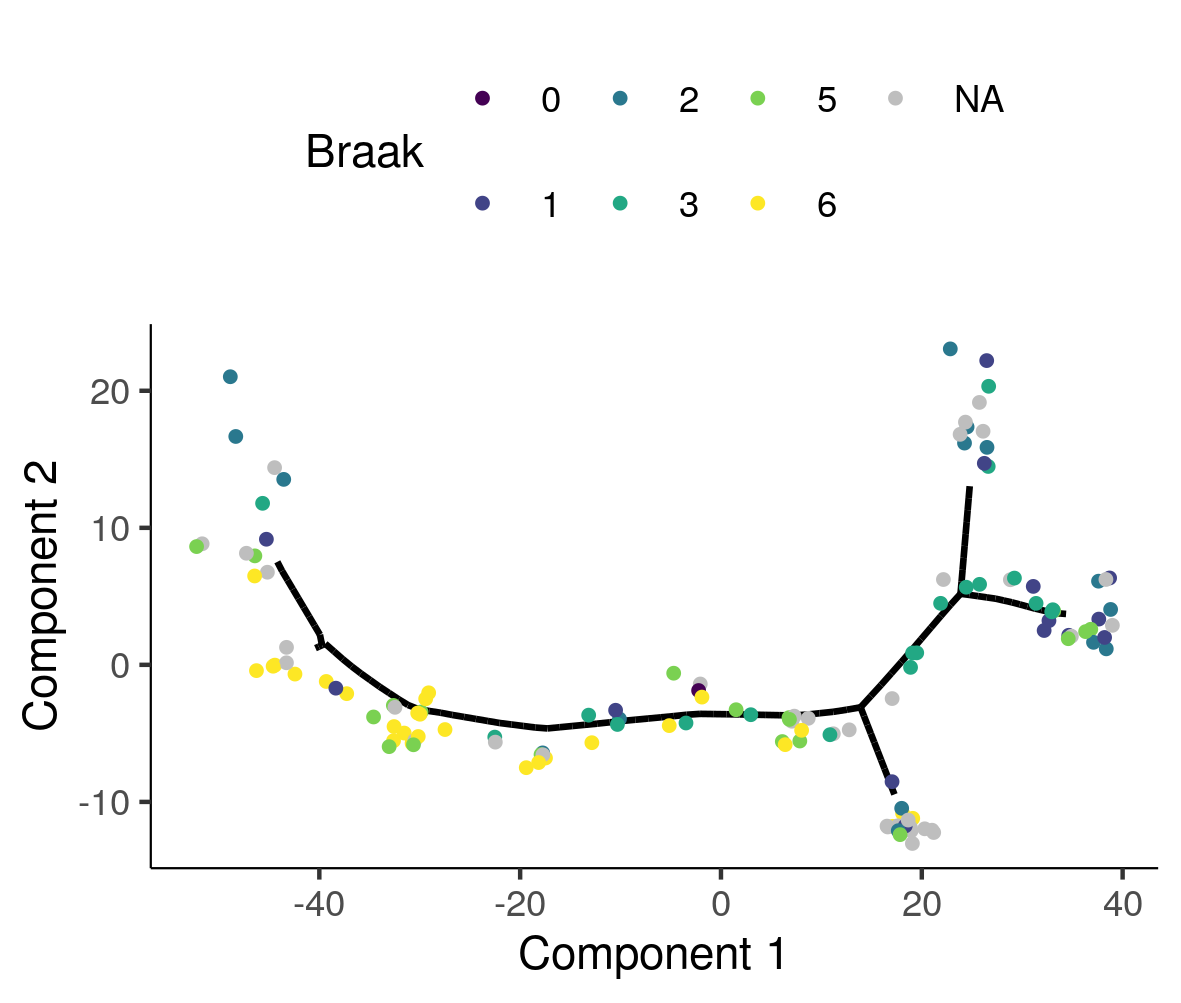

**Supplementary Figure 15 –** Pearson correlation between Pseuodtime and principal component 1 (A), 2 (B), tSNE component 1 (C), 2 (D), and UMAP component 1 (E), and 2 (F) for ROS/MAP (DLPFC).

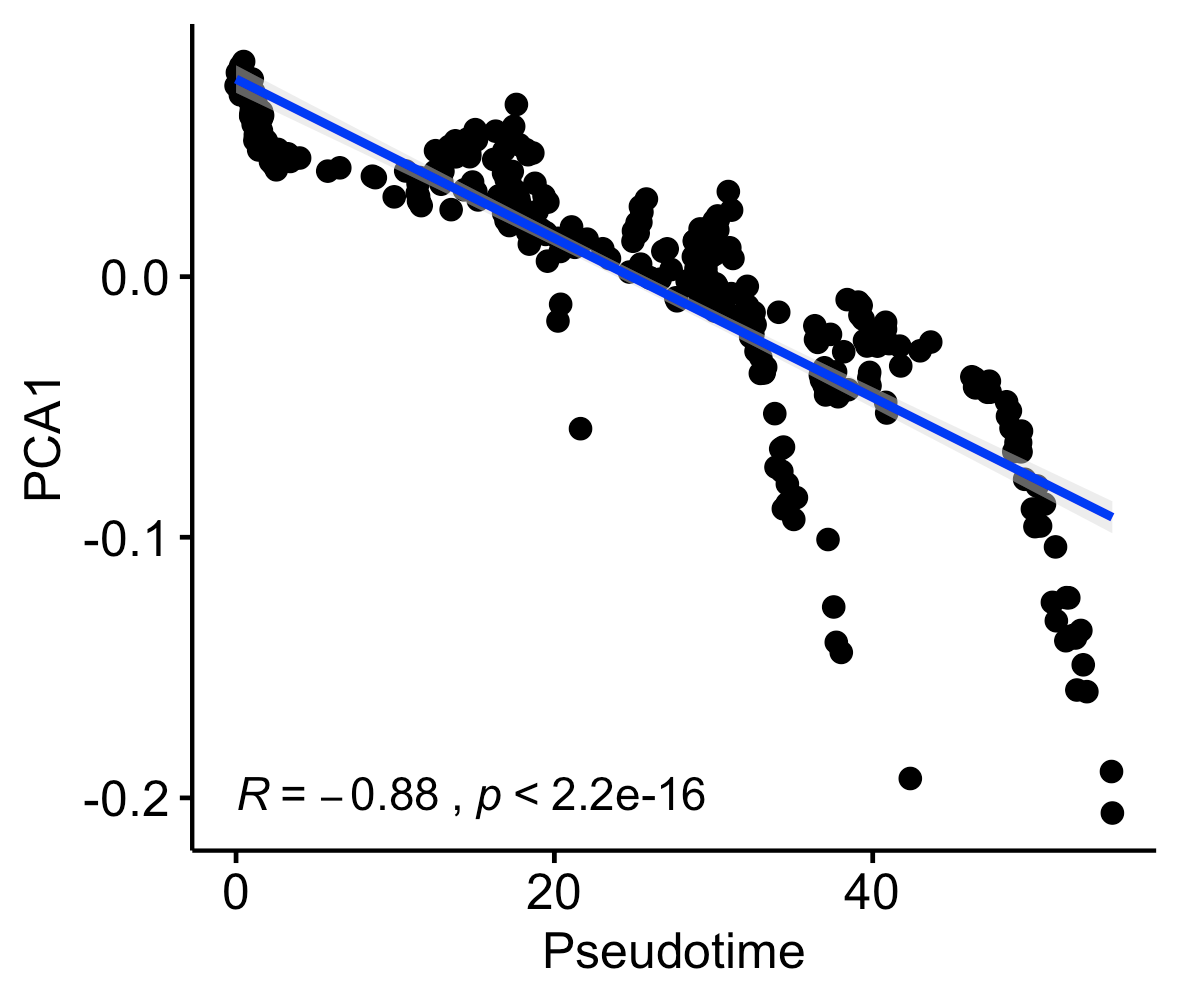

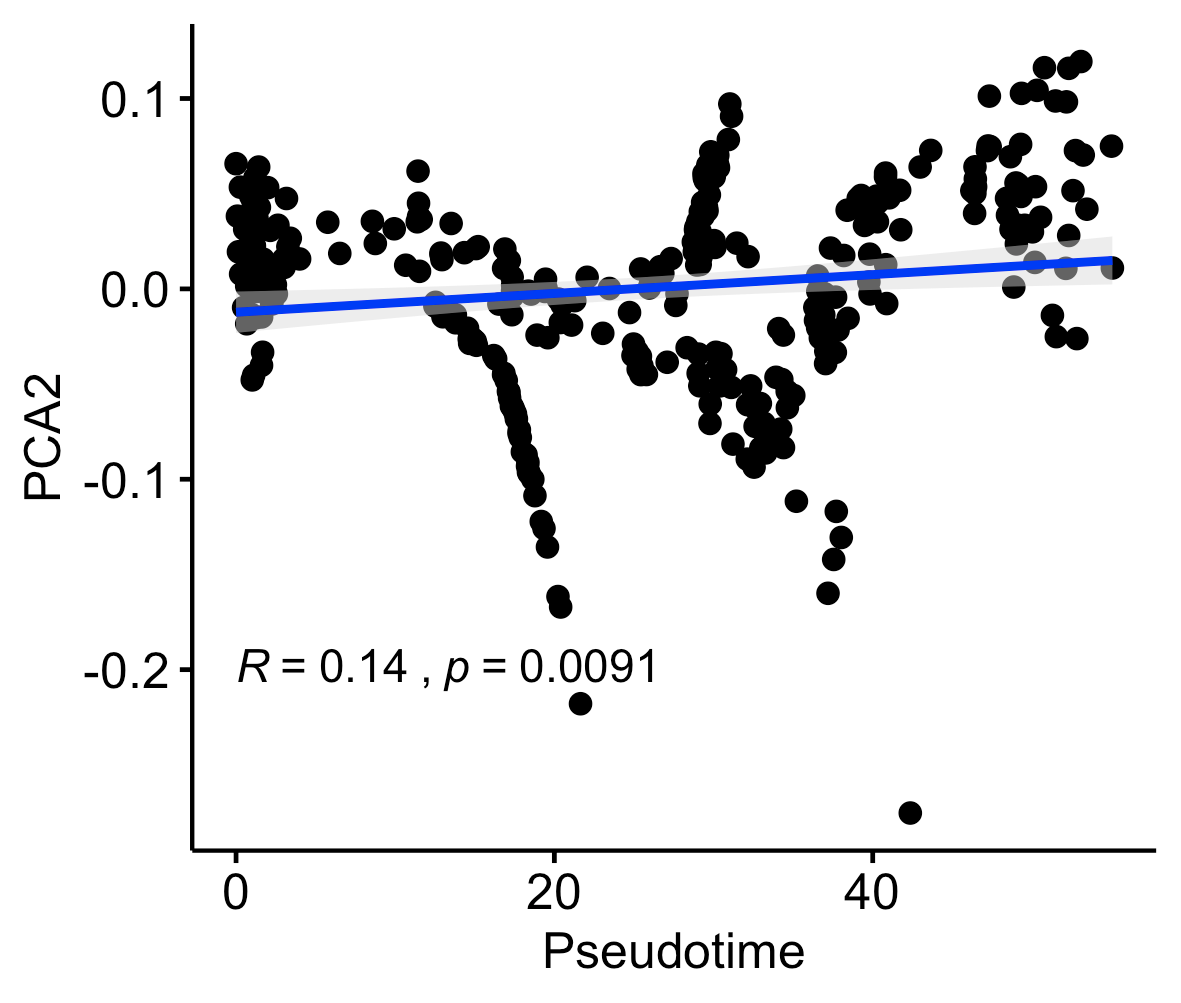

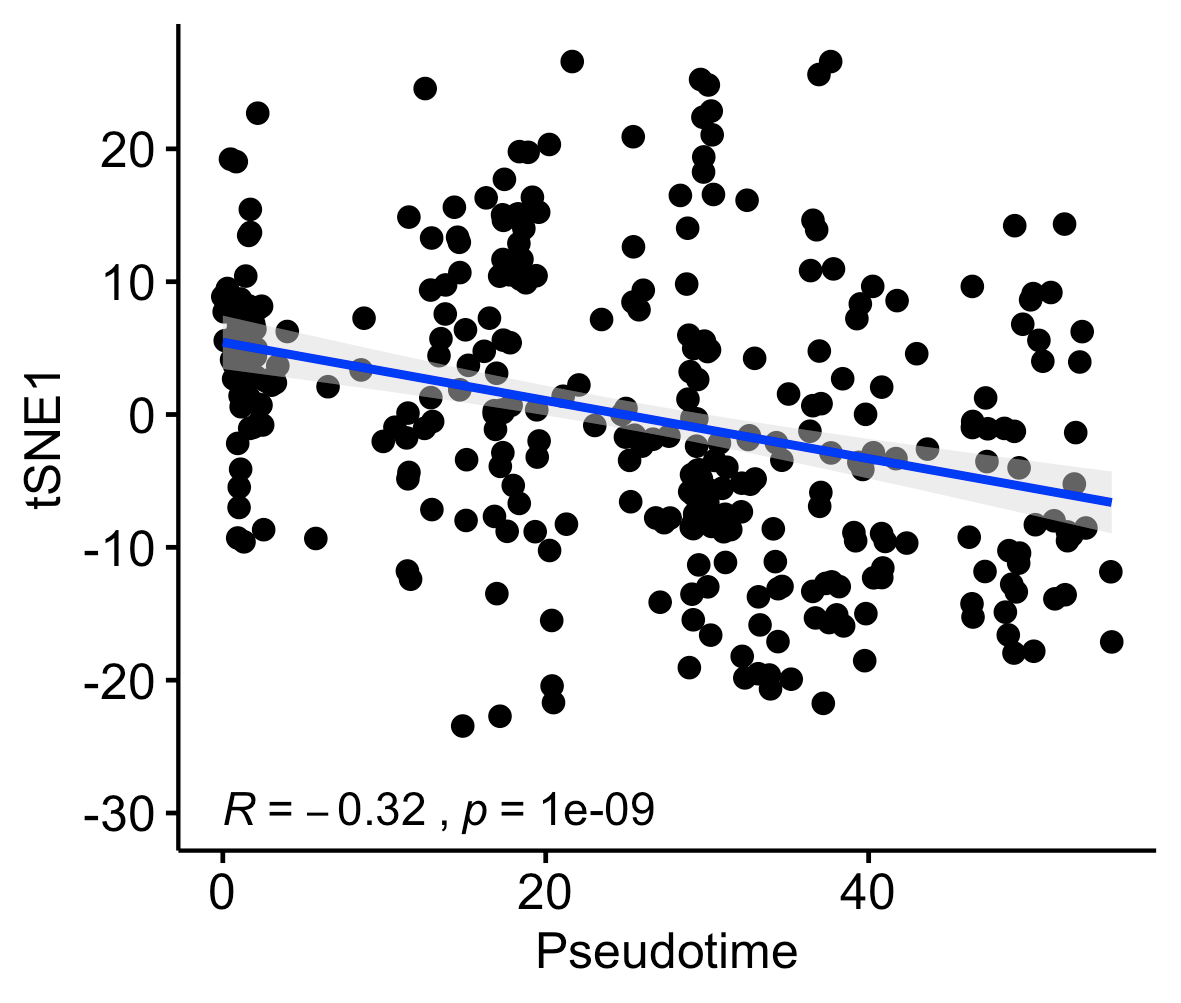

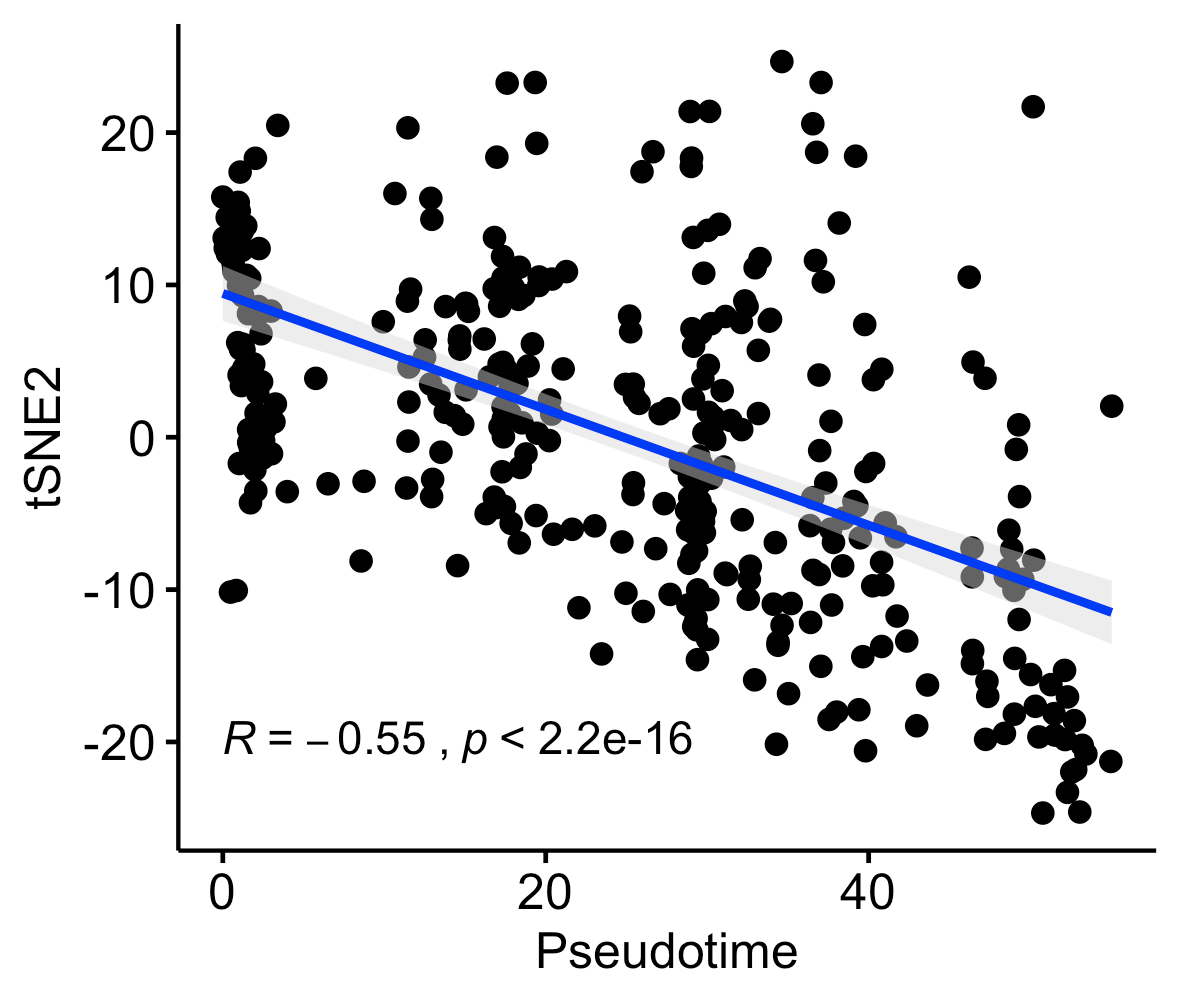

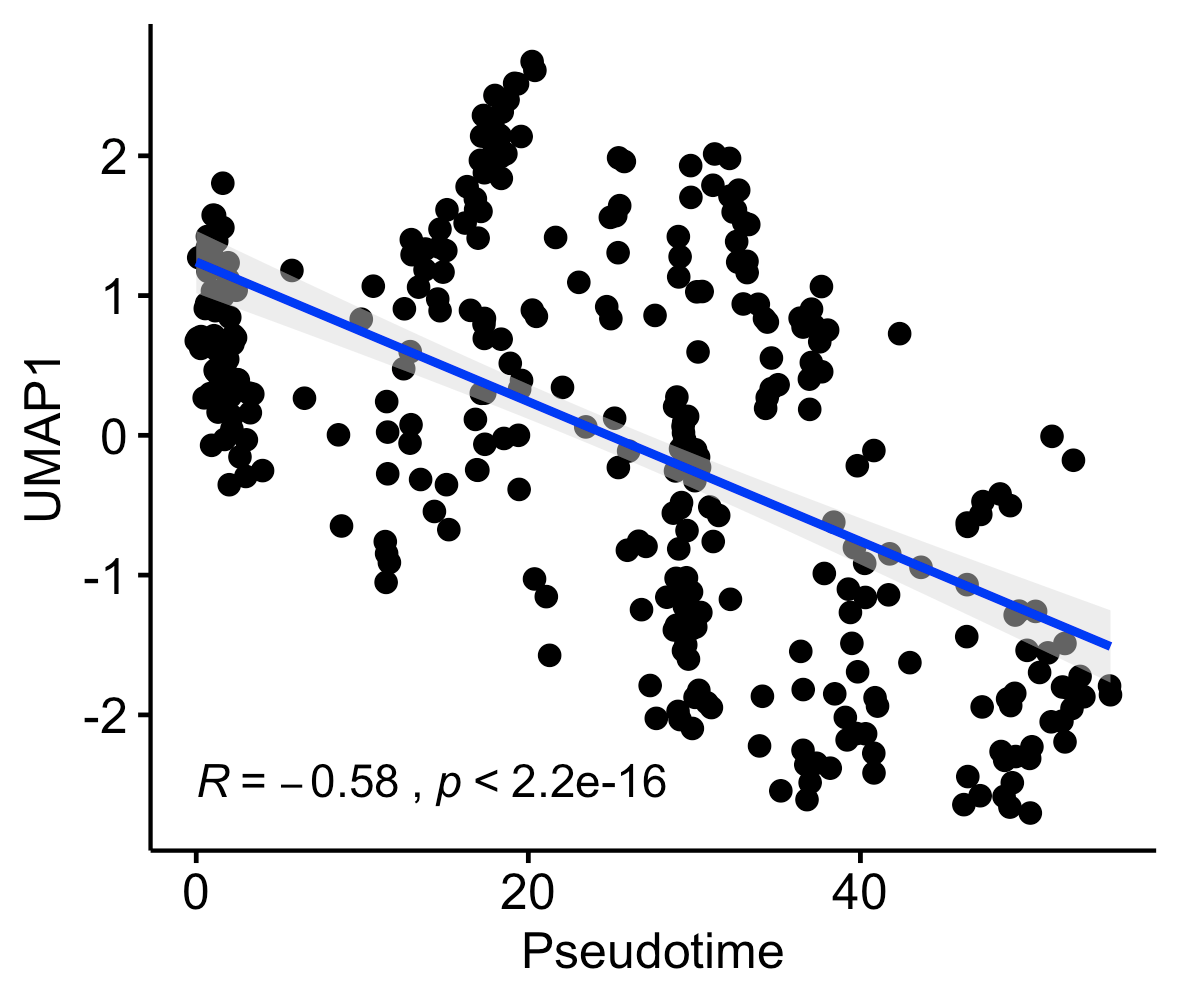

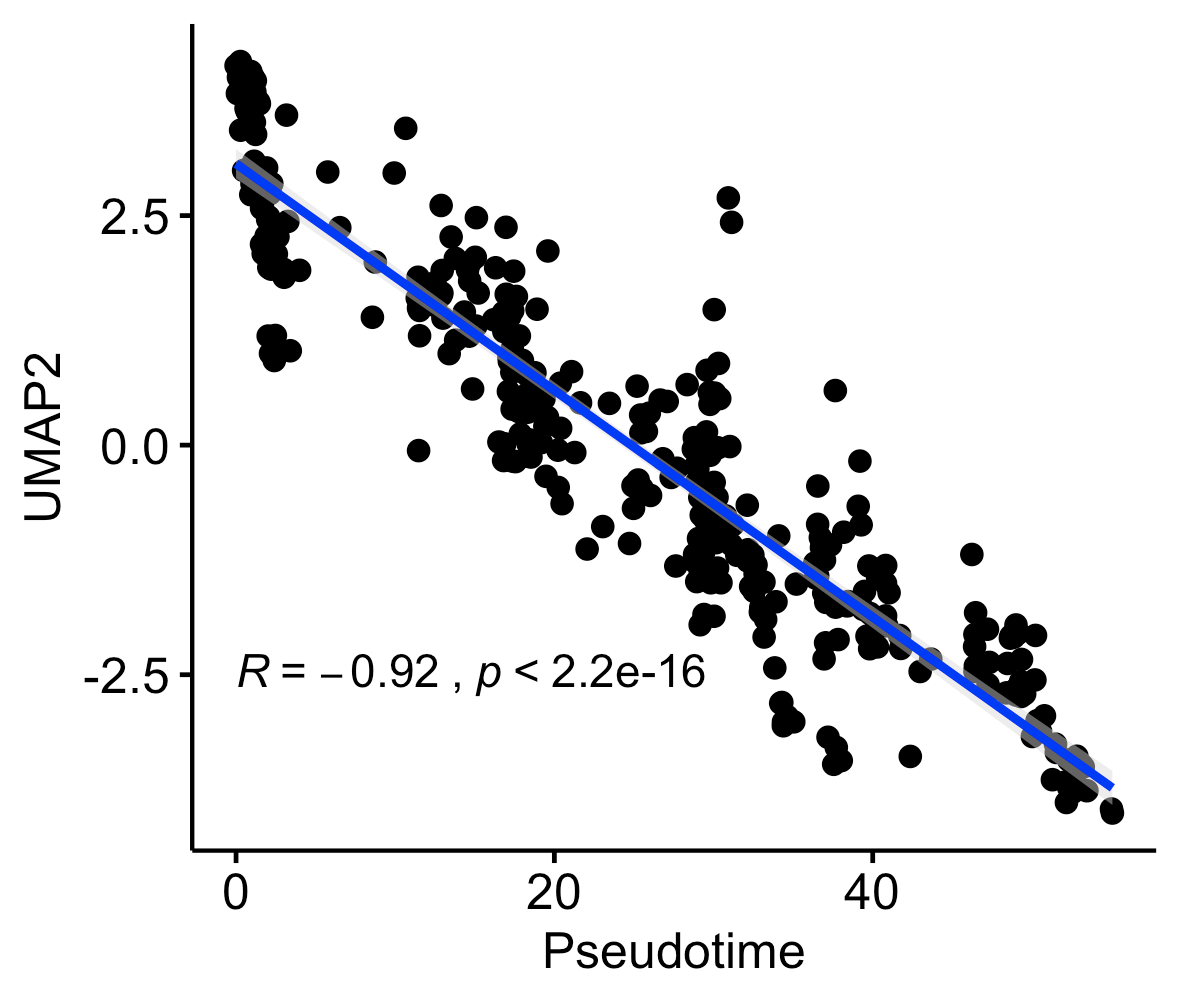

A

D

C

B

F

E

**Figure S16 -** Pearson correlation between Pseuodtime and principal component 1 (A), 2 (B), tSNE component 1 (C), 2 (D), and UMAP component 1 (E), and 2 (F) for Mayo RNAseq (TCX).

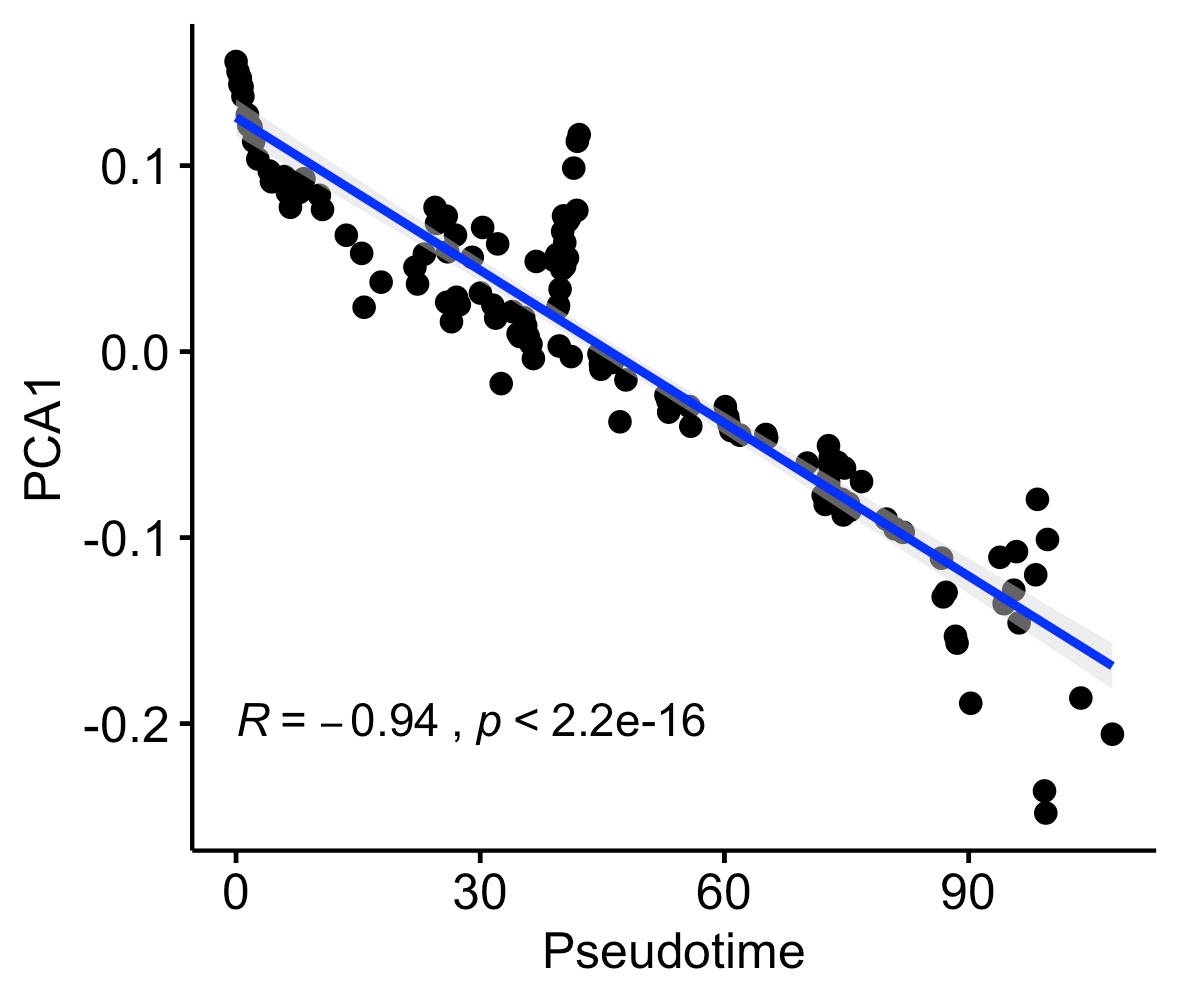

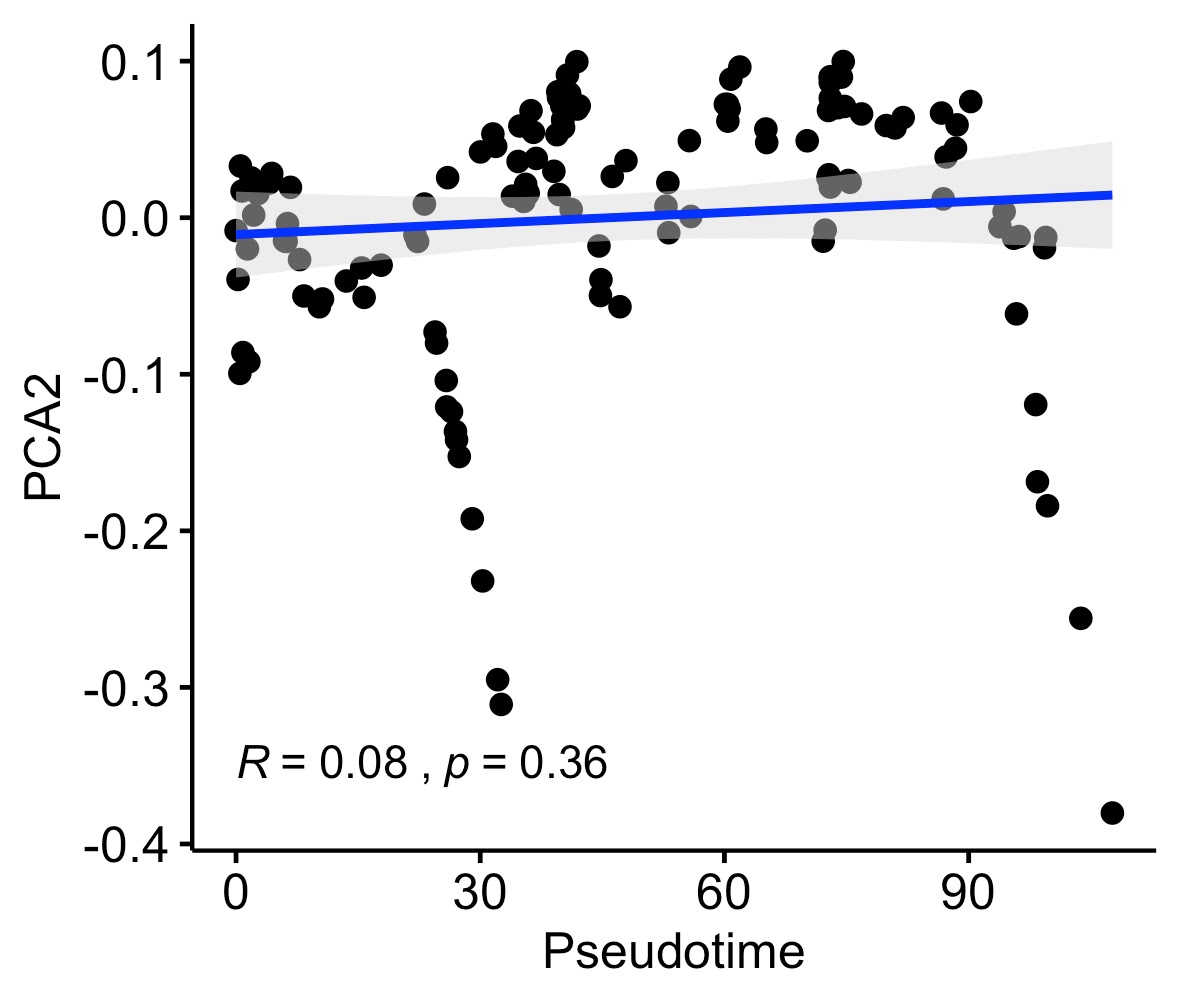

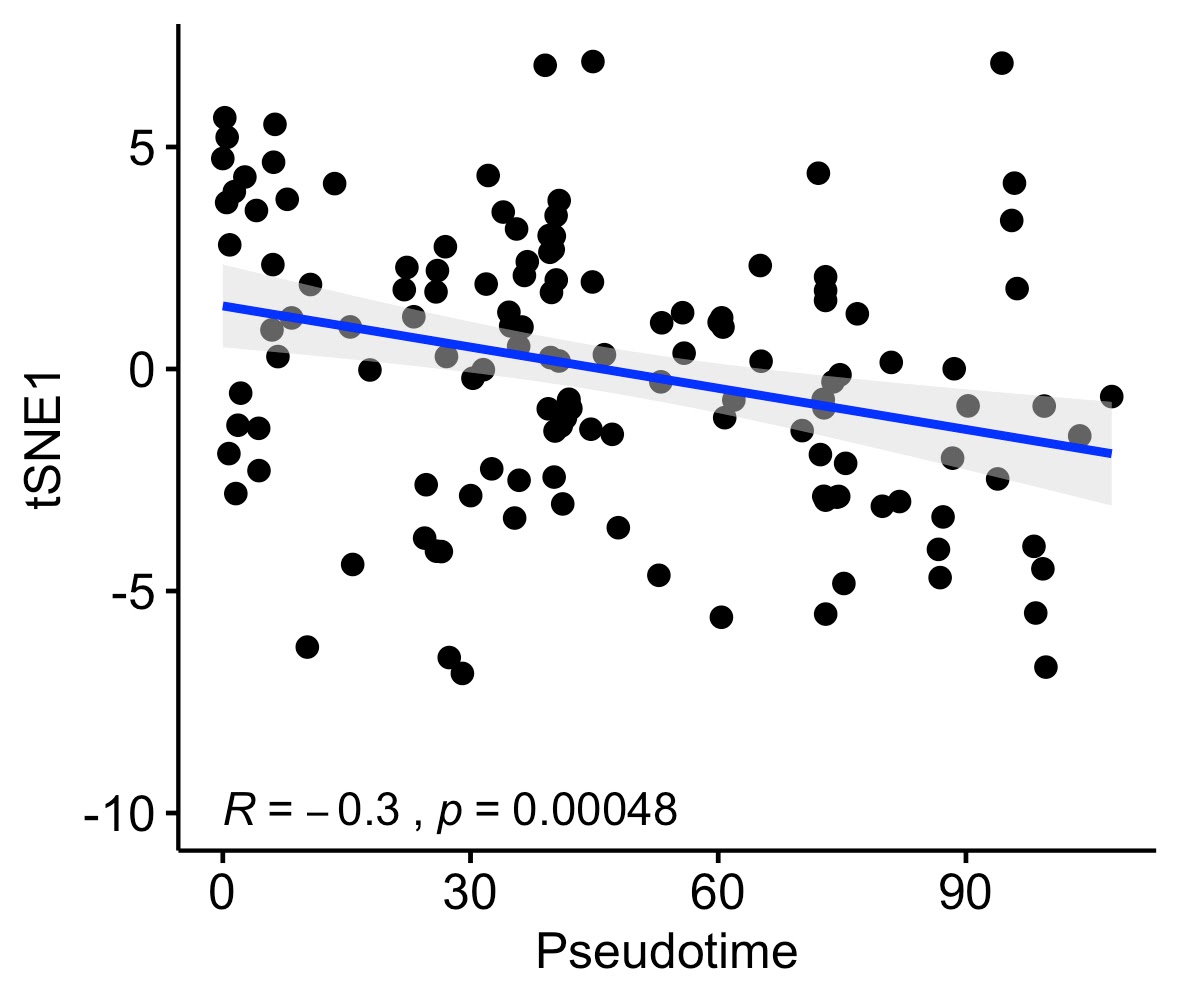

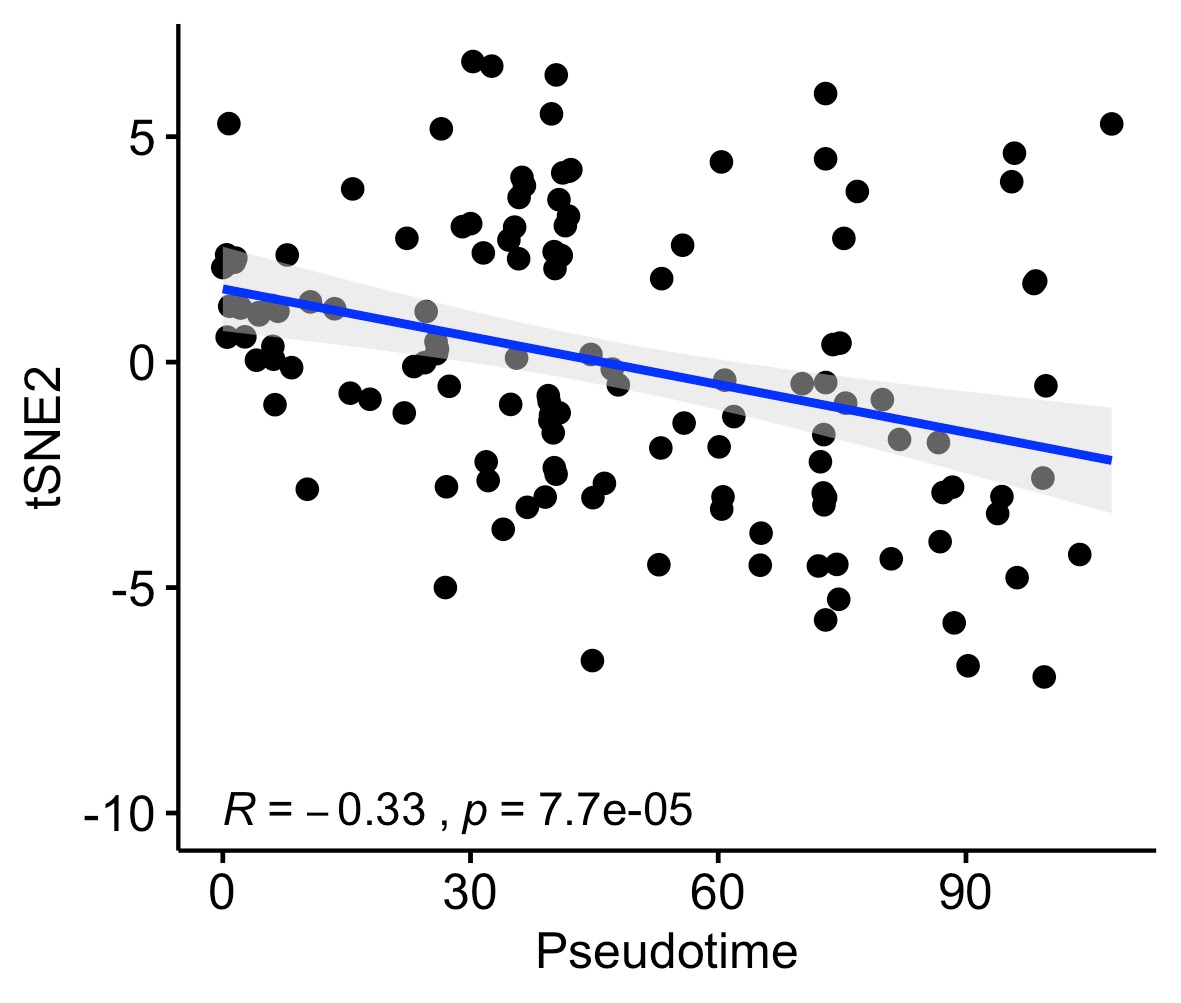

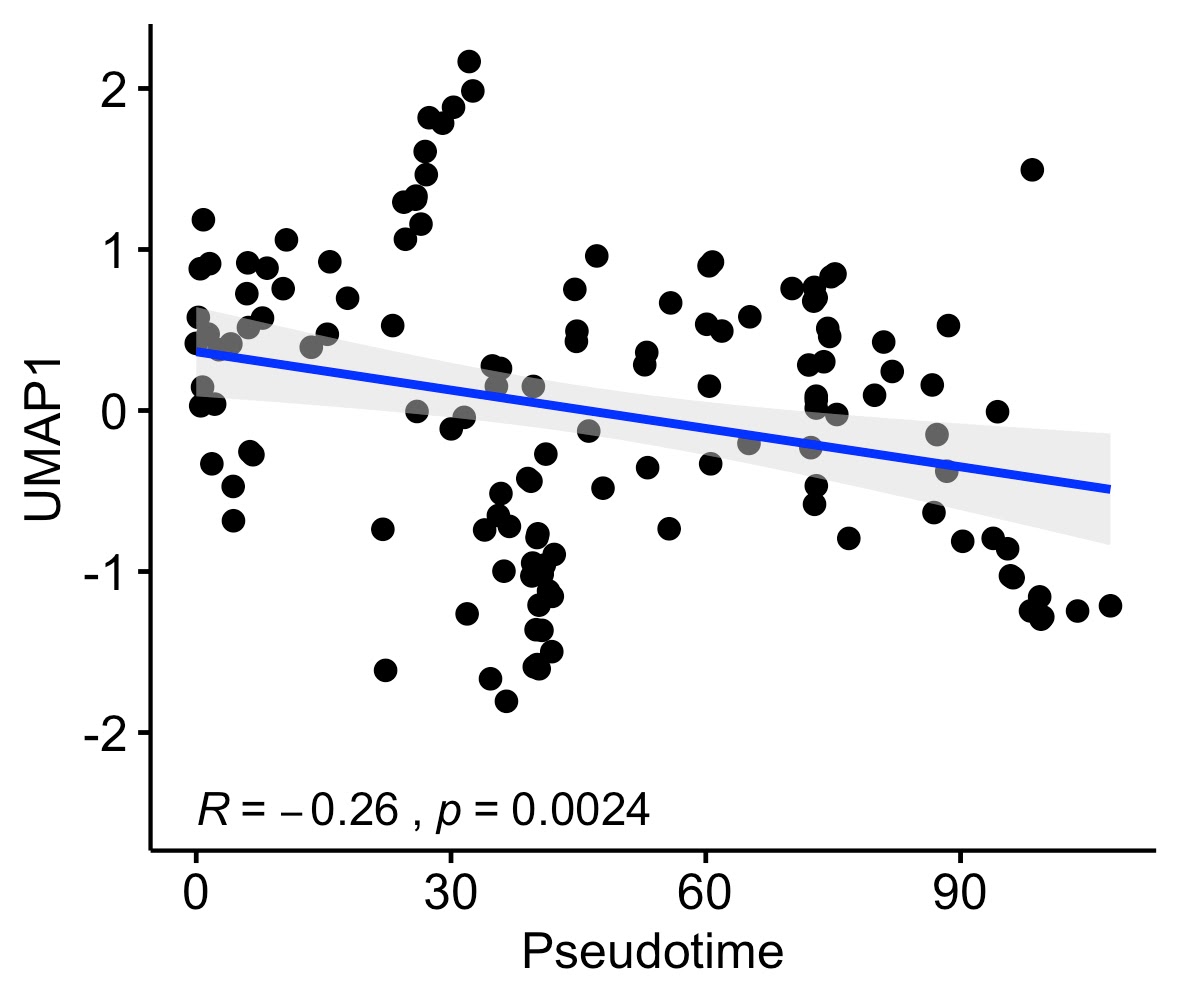

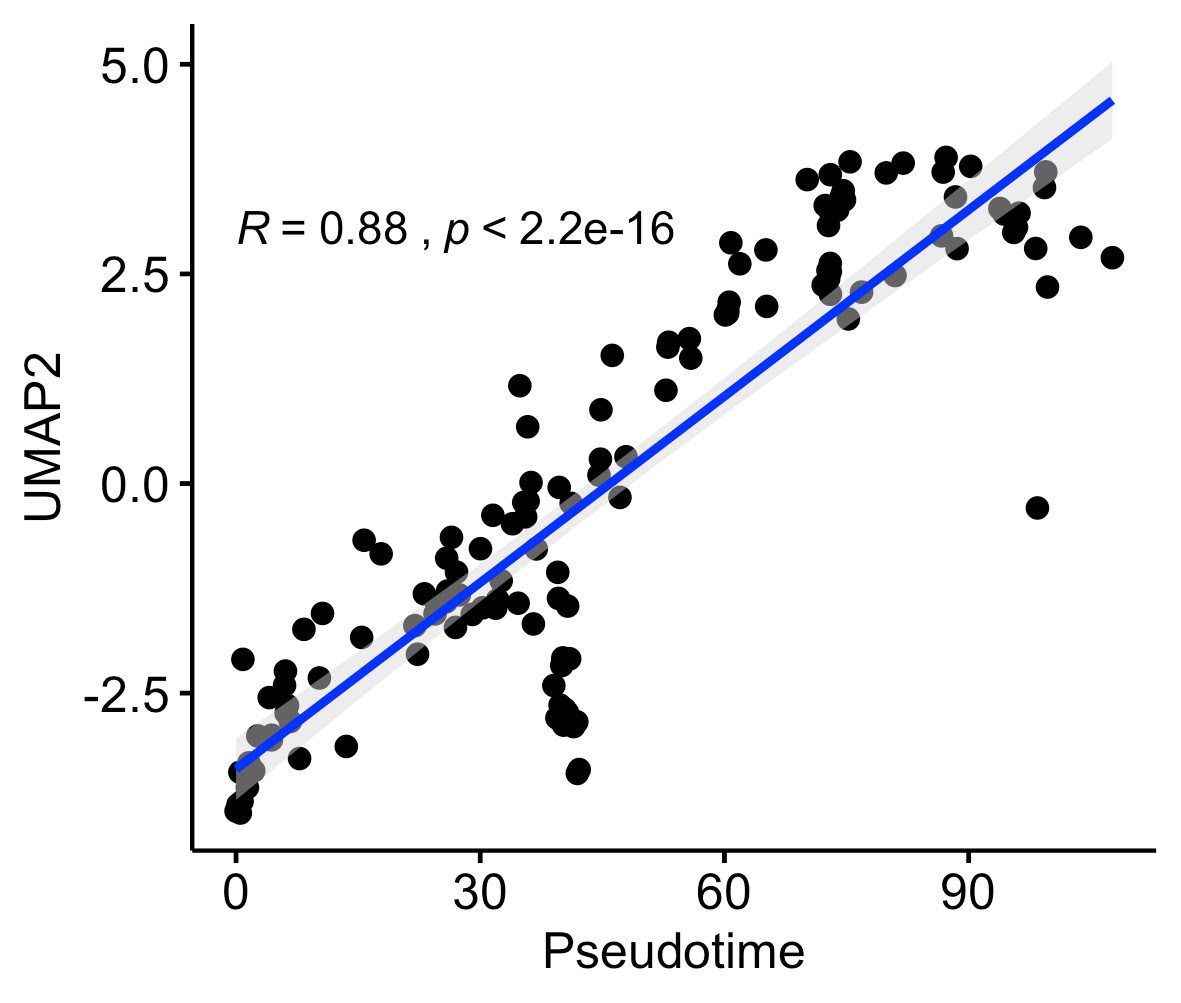

A

D

C

B

F

E

**Supplementary Figure 17 –** Monocle 3 trajectories and associations: UMAP method (Monocle3) for 218 independent samples from two independent studies, A) Lineage learned in TCX, B) Lineage learned in DLPFC, C) Association between disease pseudotime and diagnosis, D) correlation between pseudotime and IGAP GWAS genes. Box plots have lower and upper hinges at the 25th and 75th percentiles and whiskers extending to at most 1.5xIQR (interquartile range).

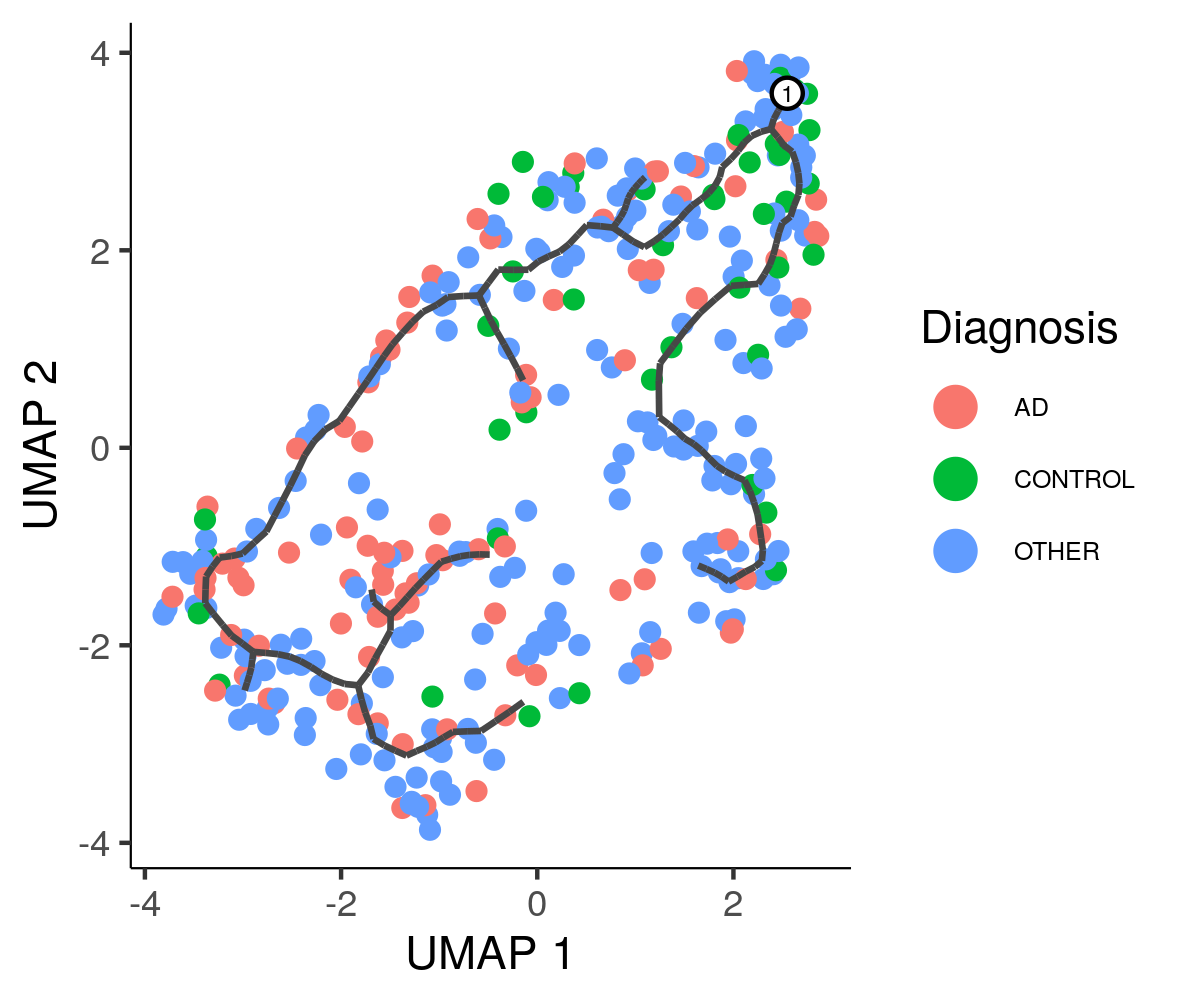

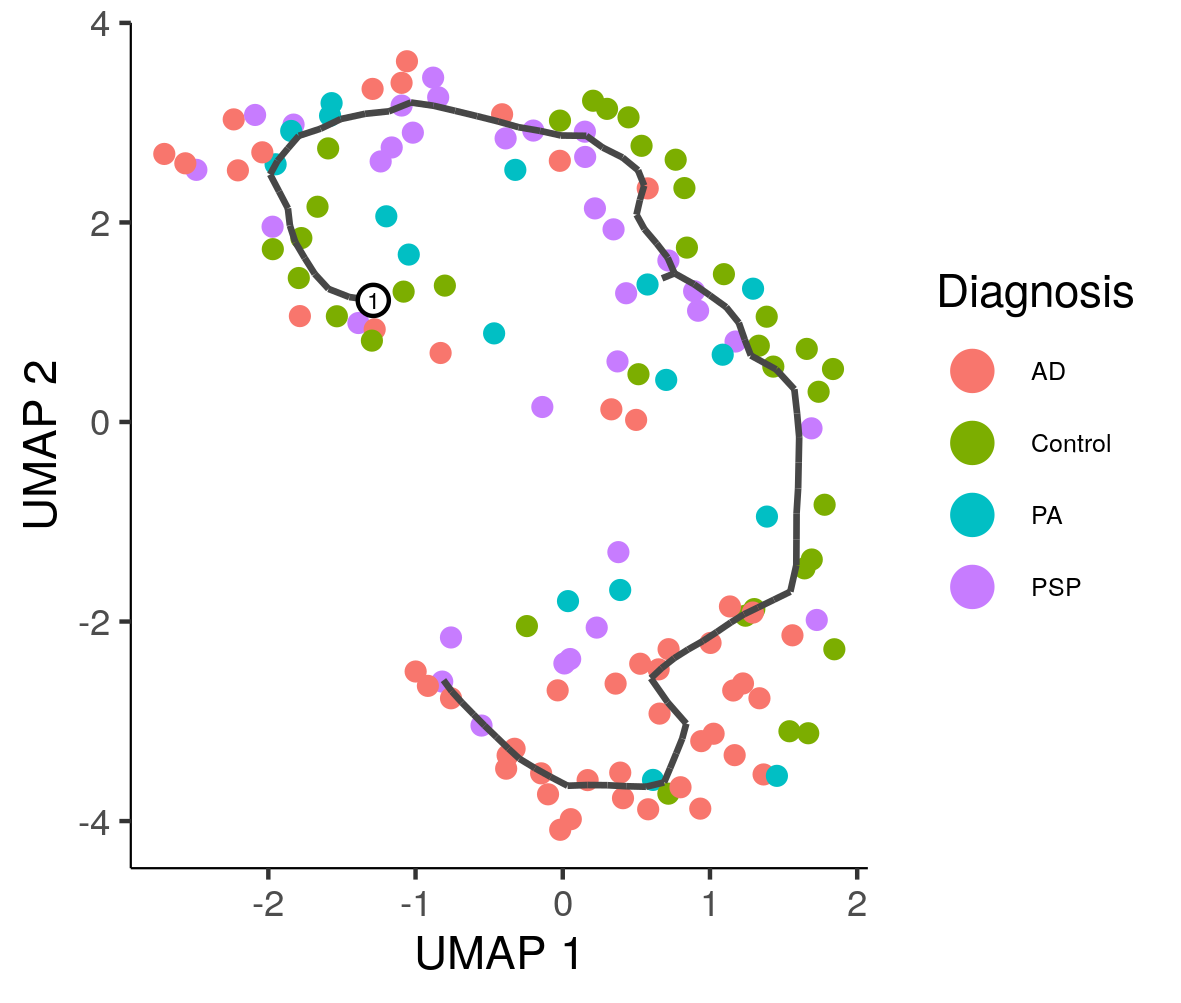

TCX

DLPFC

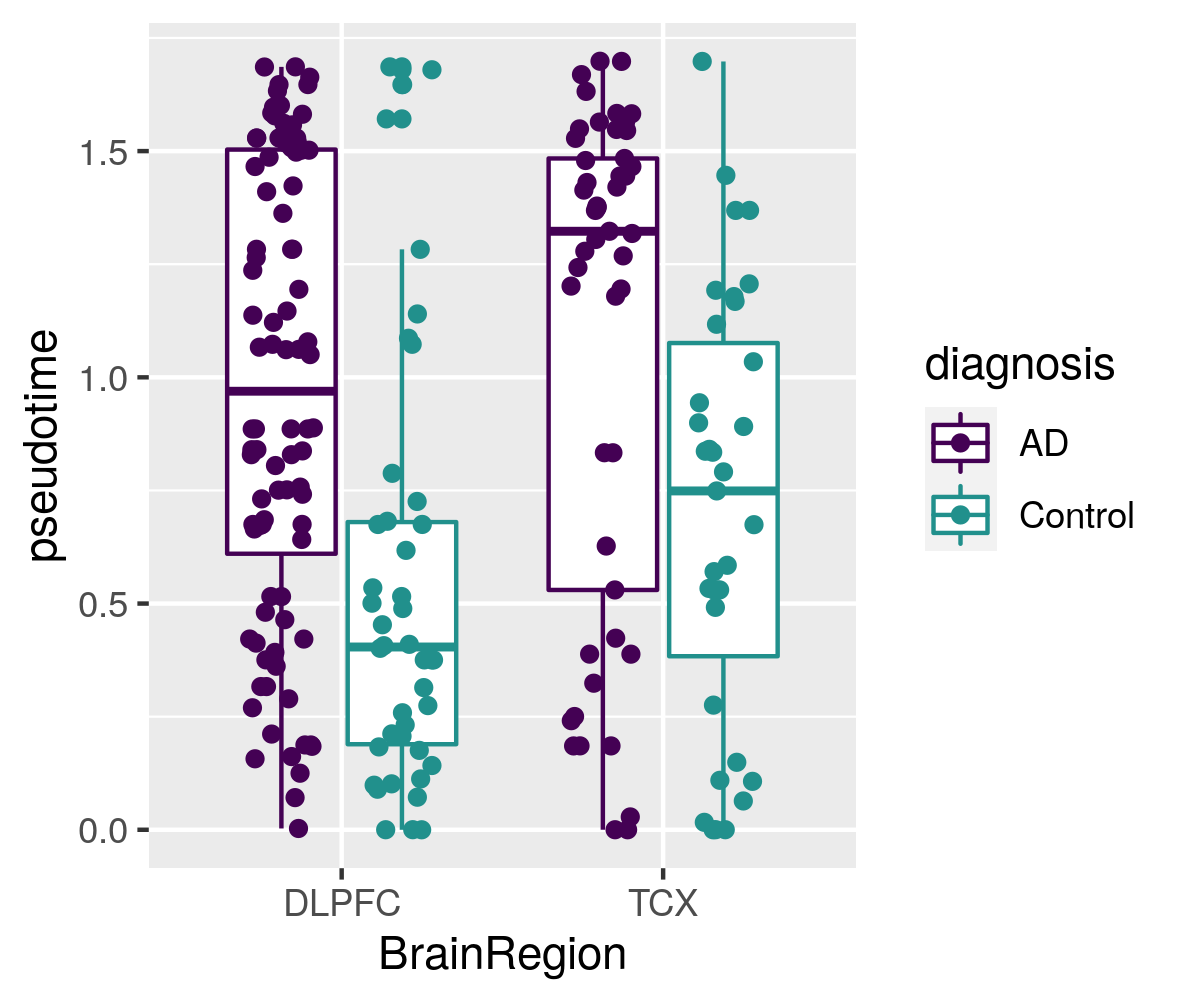

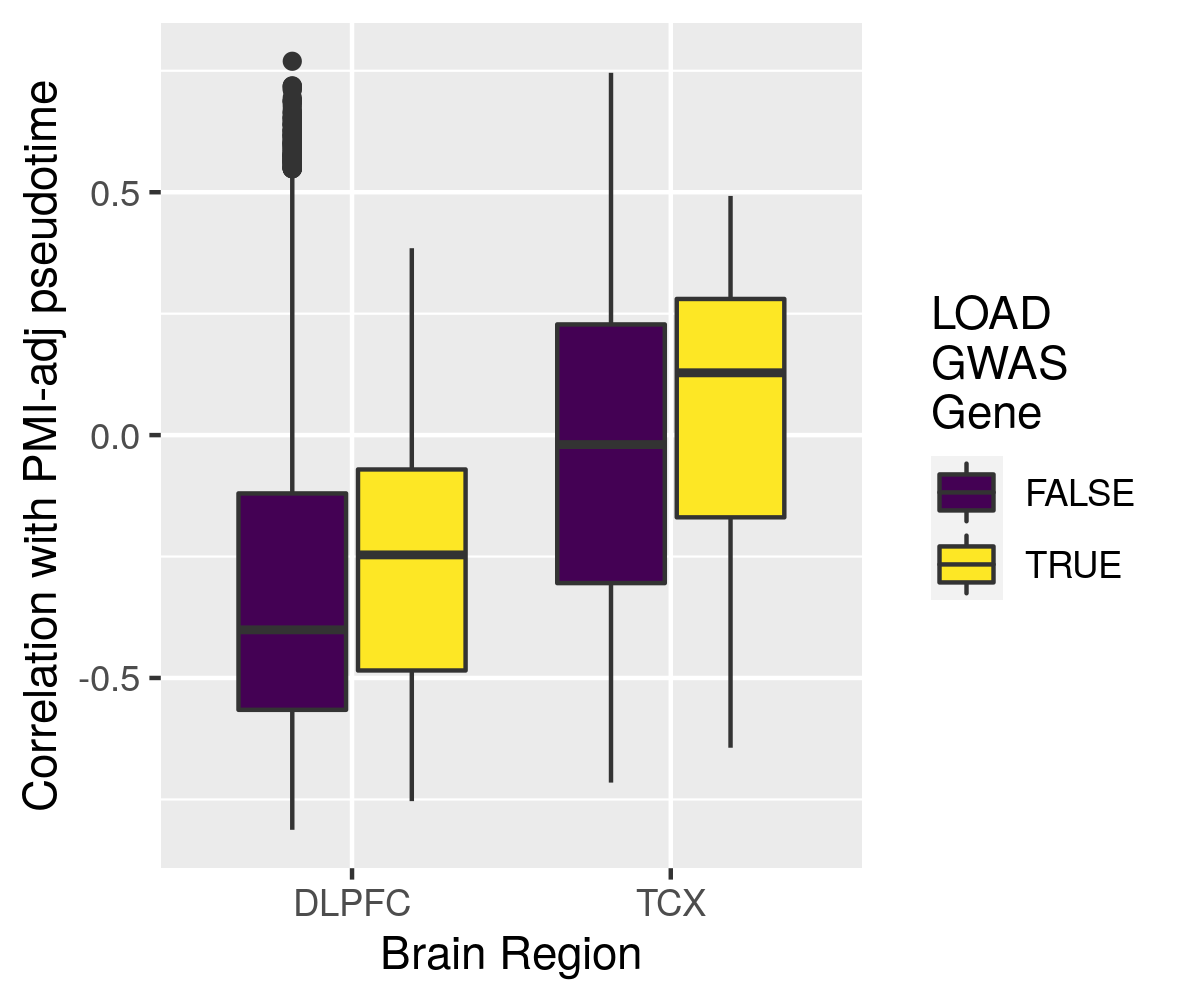

A

B

C

D

**Supplementary Figure 18 –** Monocle 3 trajectories and neuropath associations in DLPFC: UMAP method (Monocle3) for 338 independent samples from one study. A-C) Samples colored by three external measures of LOAD staging: Braak score, CERAD score, and cognitive diagnosis. D-F) Distribution of samples by inferred stage for different distinct stages in each of the three methods of measuring LOAD severity. Box plots have lower and upper hinges at the 25th and 75th percentiles and whiskers extending to at most 1.5xIQR (interquartile range).

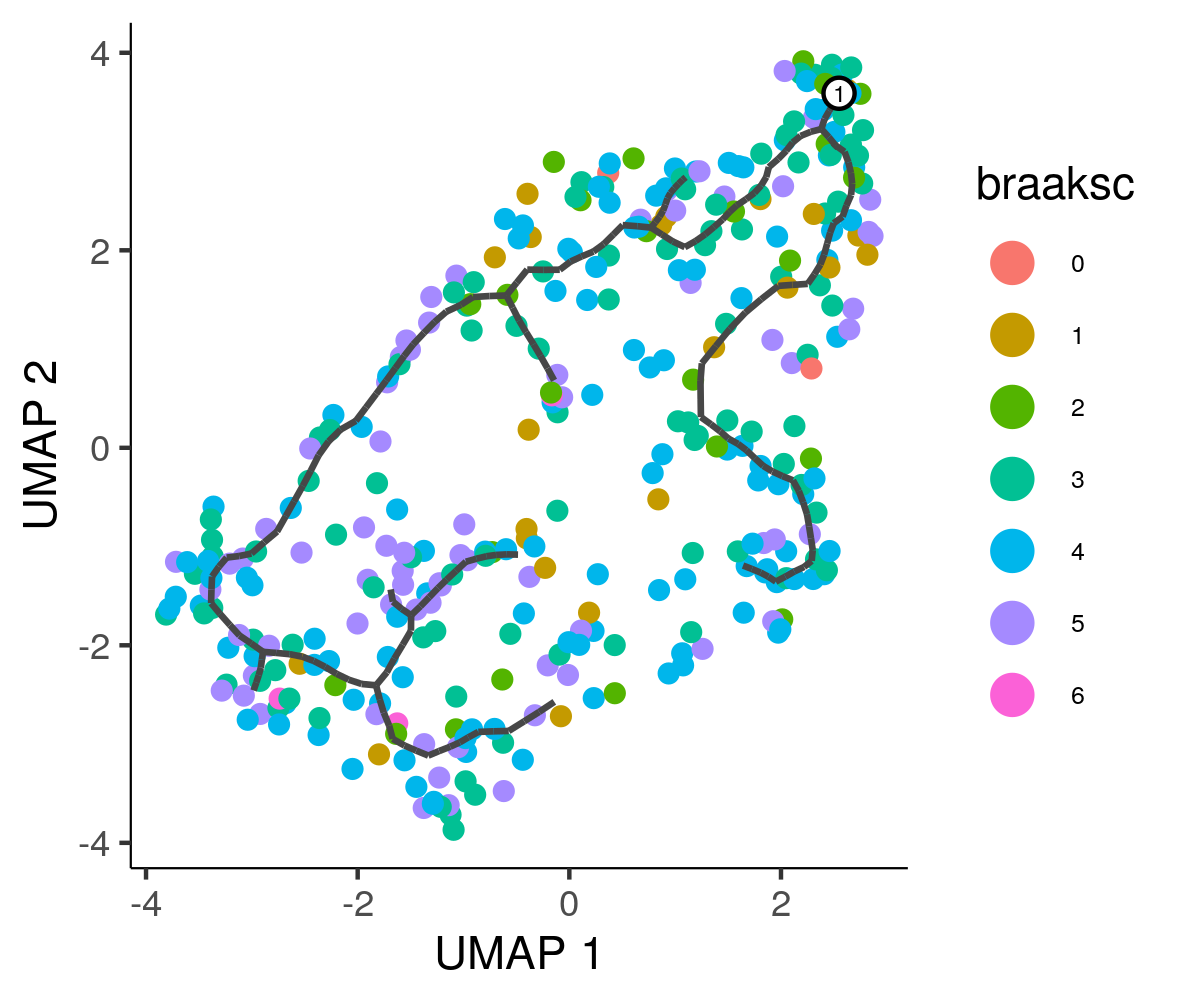

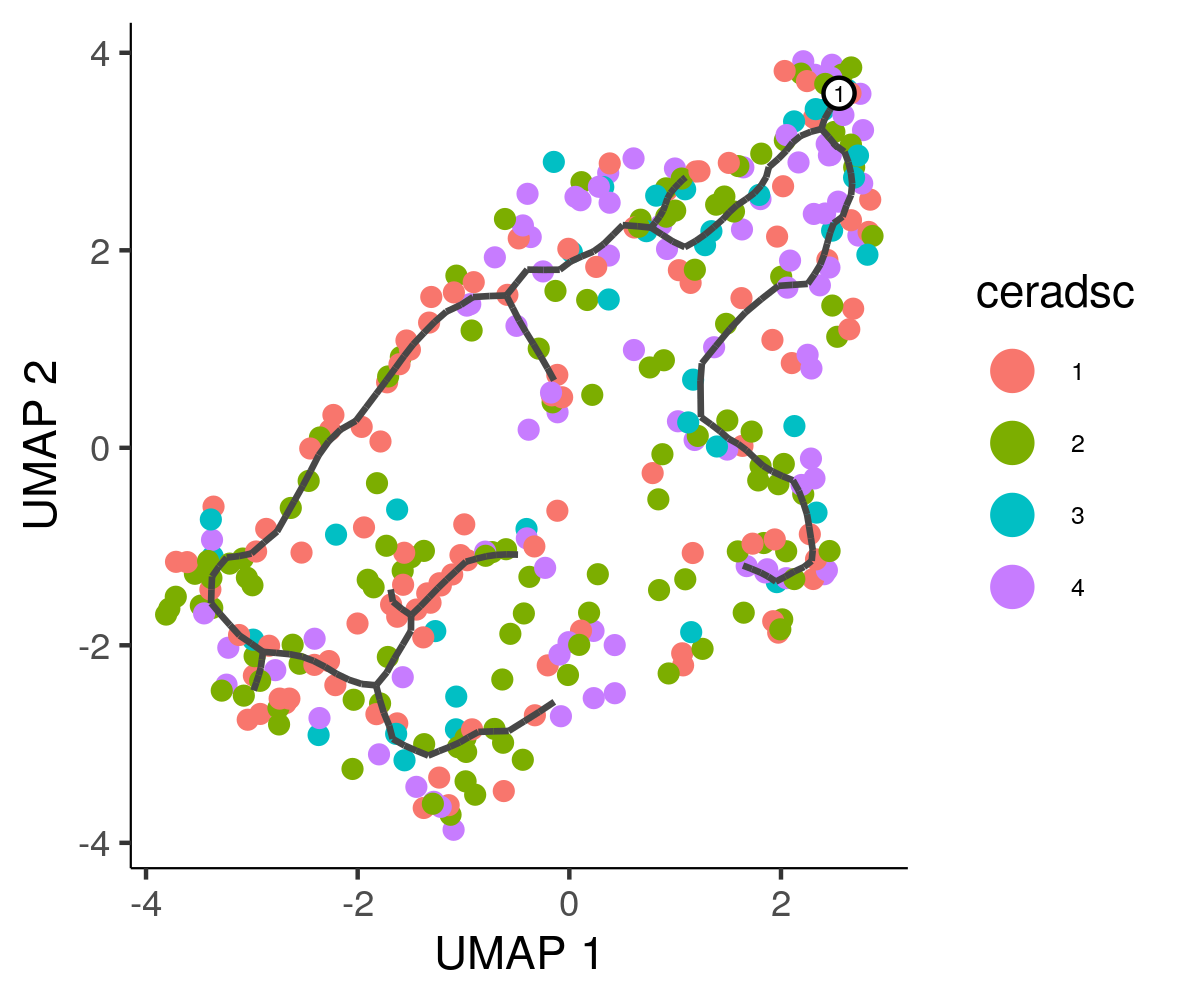

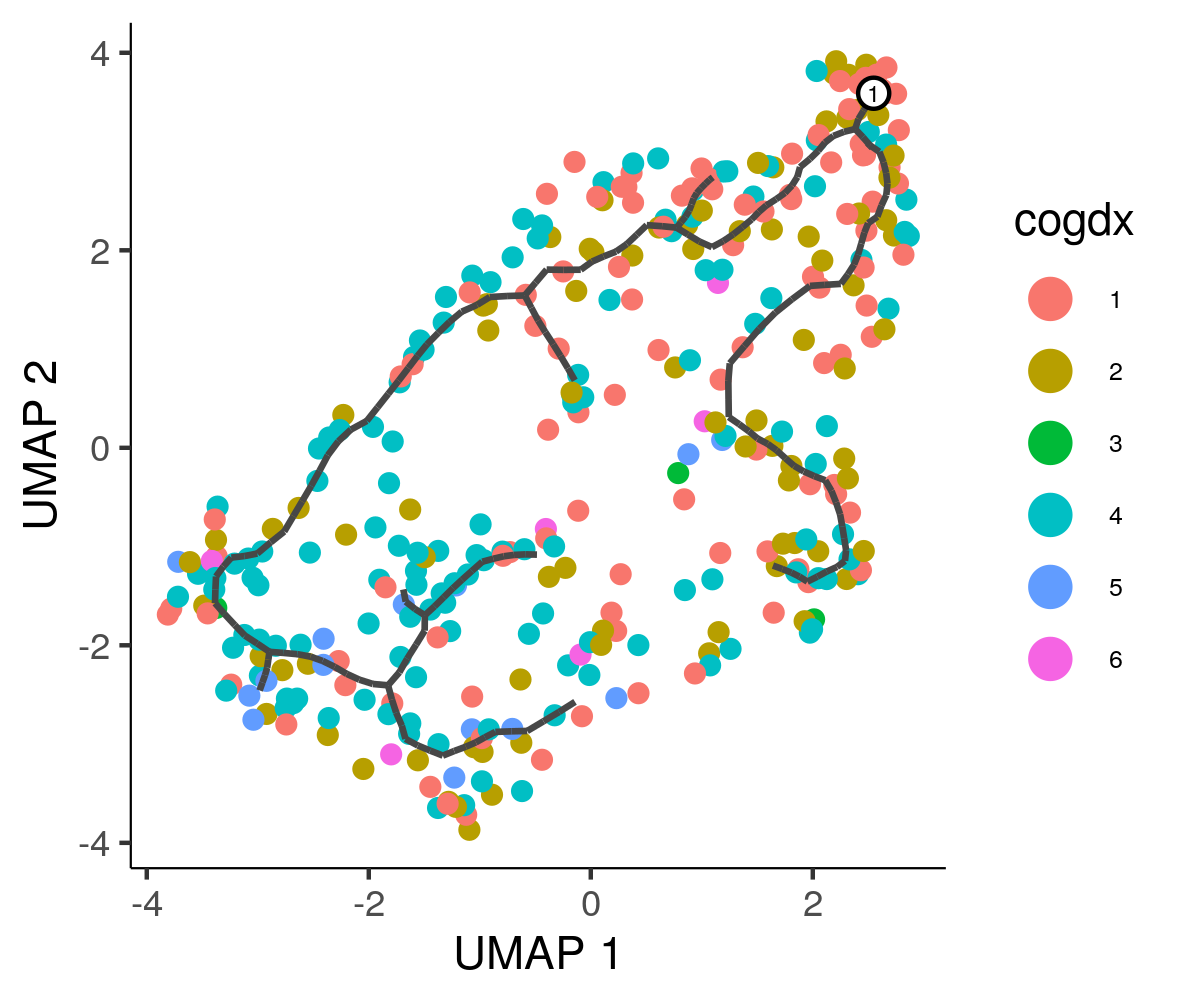

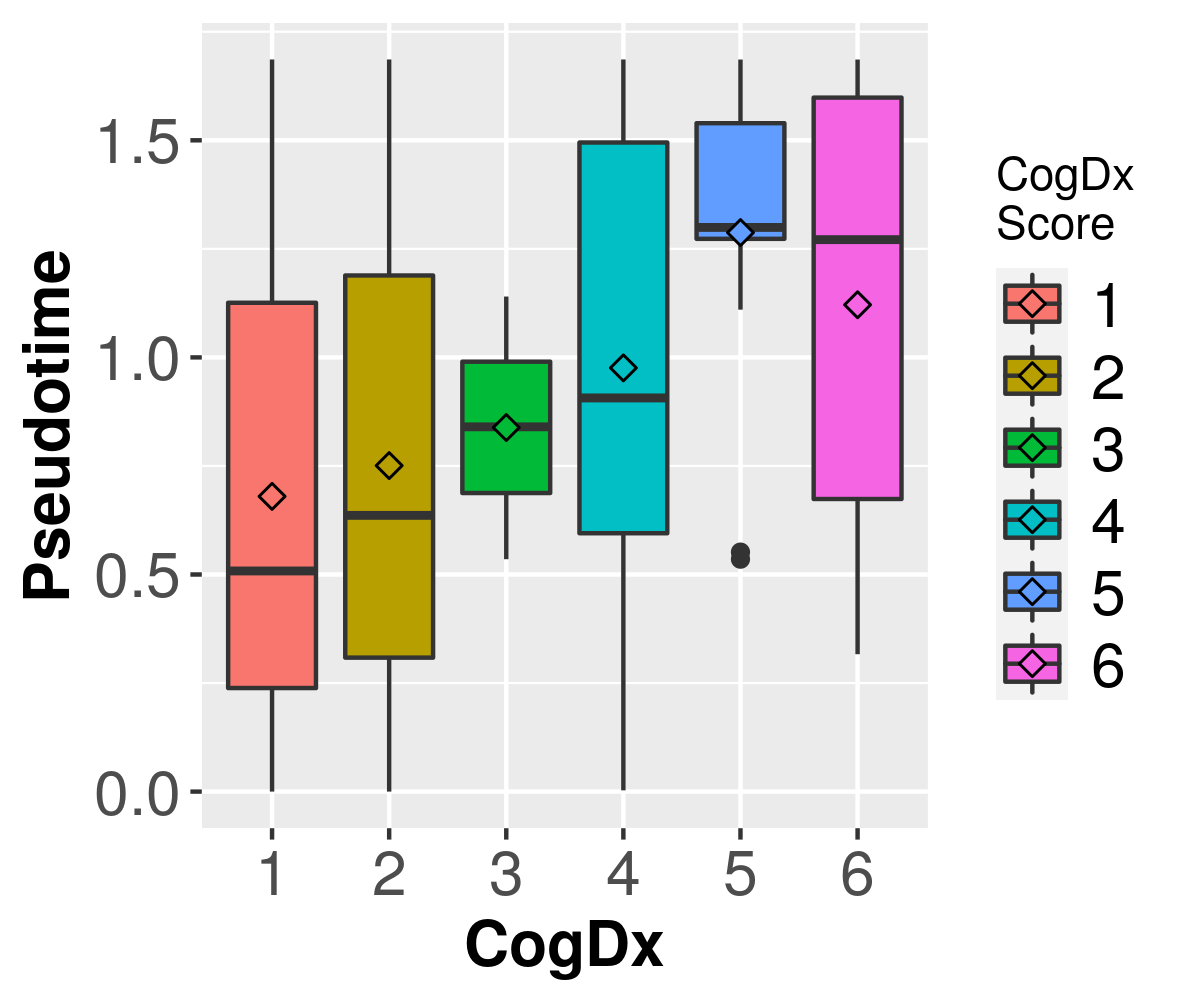

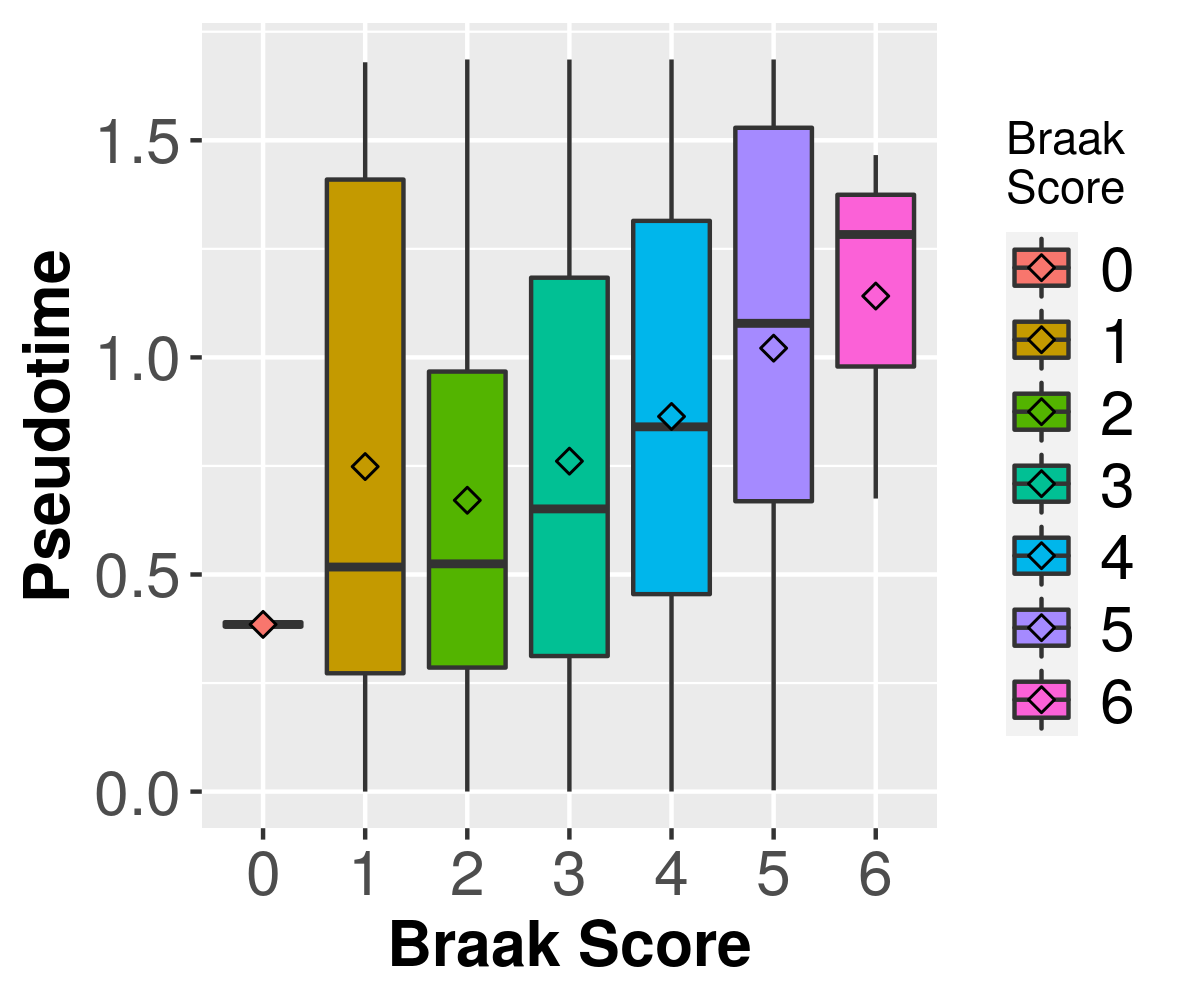

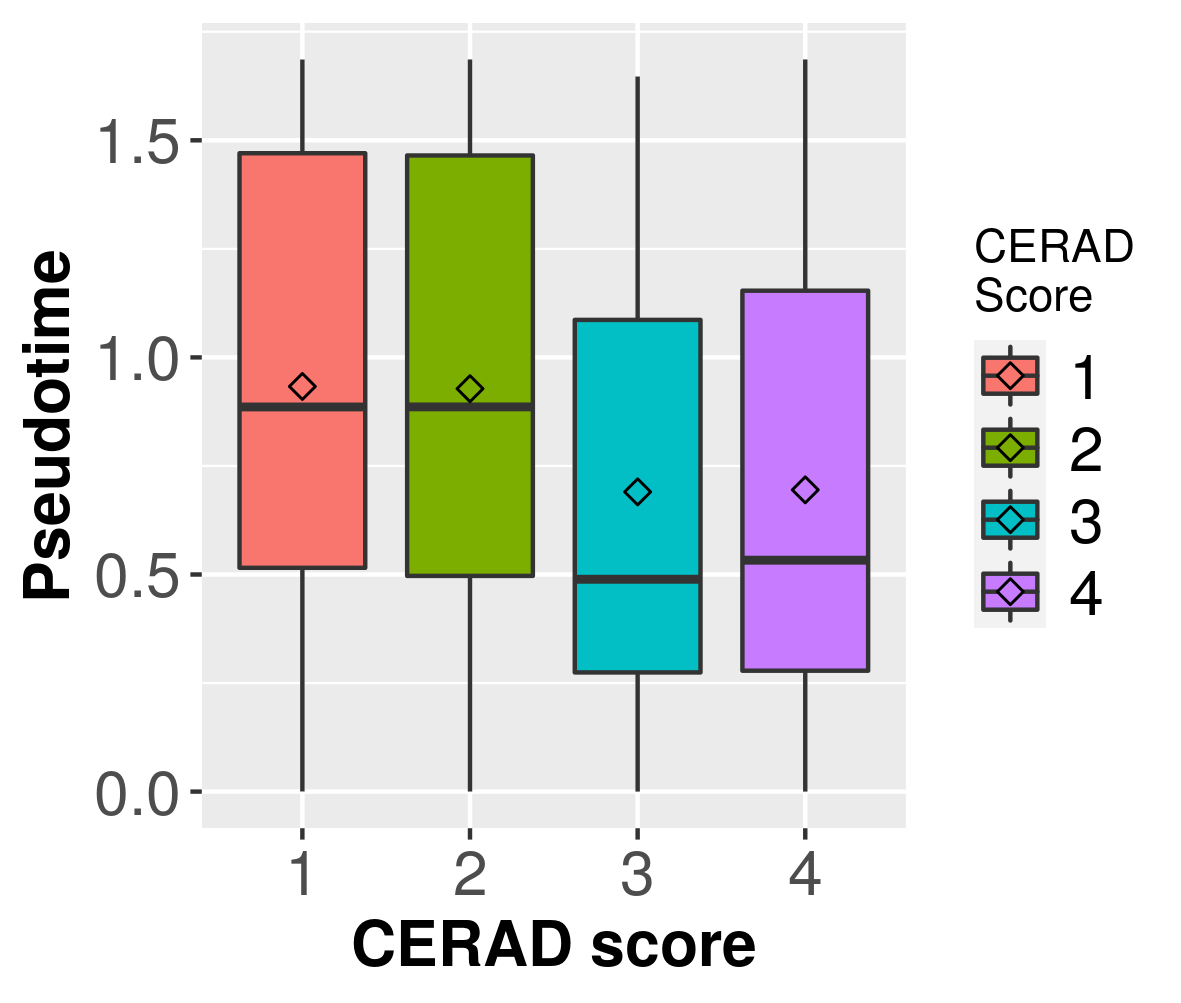

A

B

C

D

E

F

**Supplementary Figure 19 -** APOE e4 status of samples overlaid on inferred manifolds for both TCX and DLPFC brain regions.

**DLPFC**

**TCX**

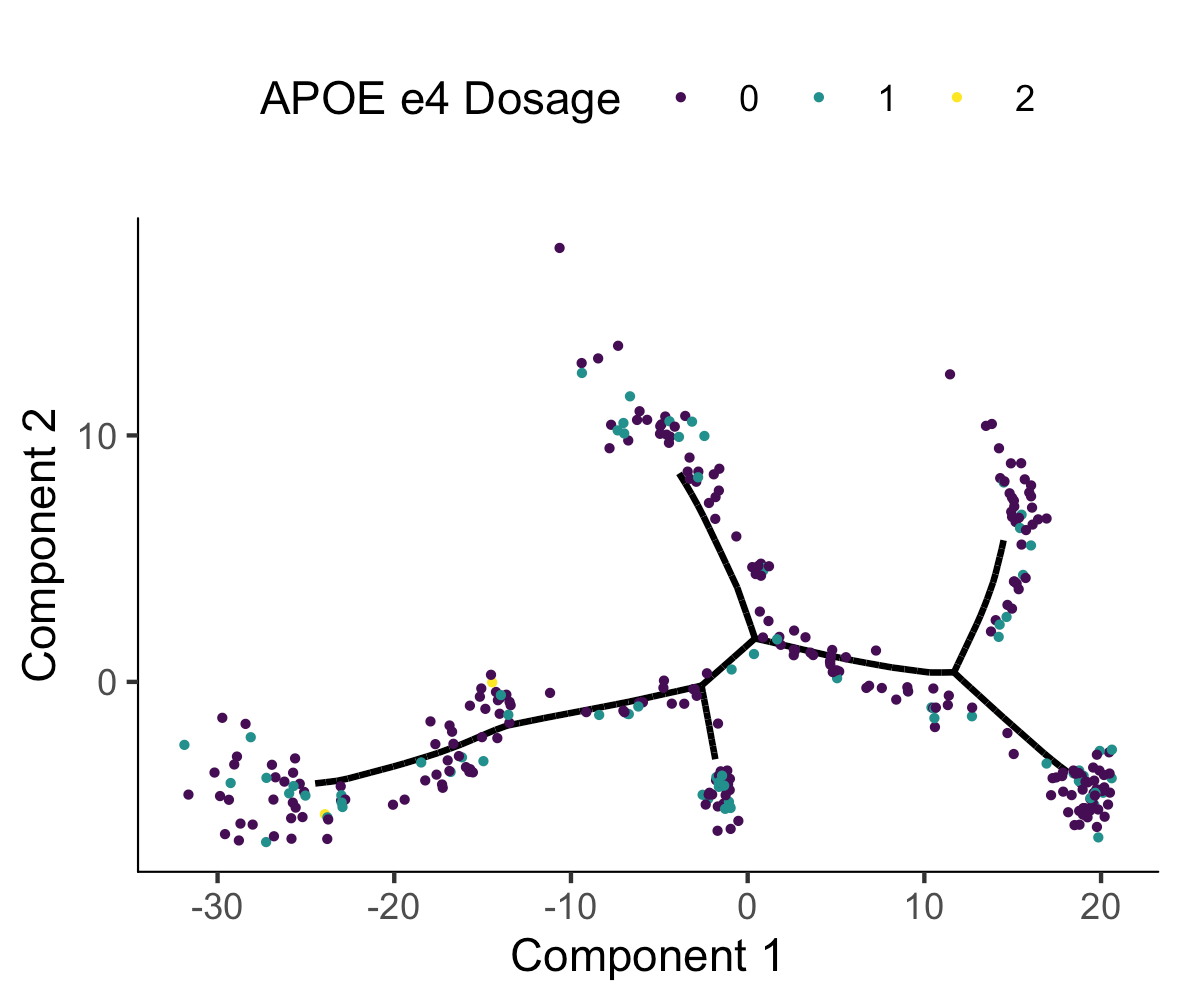

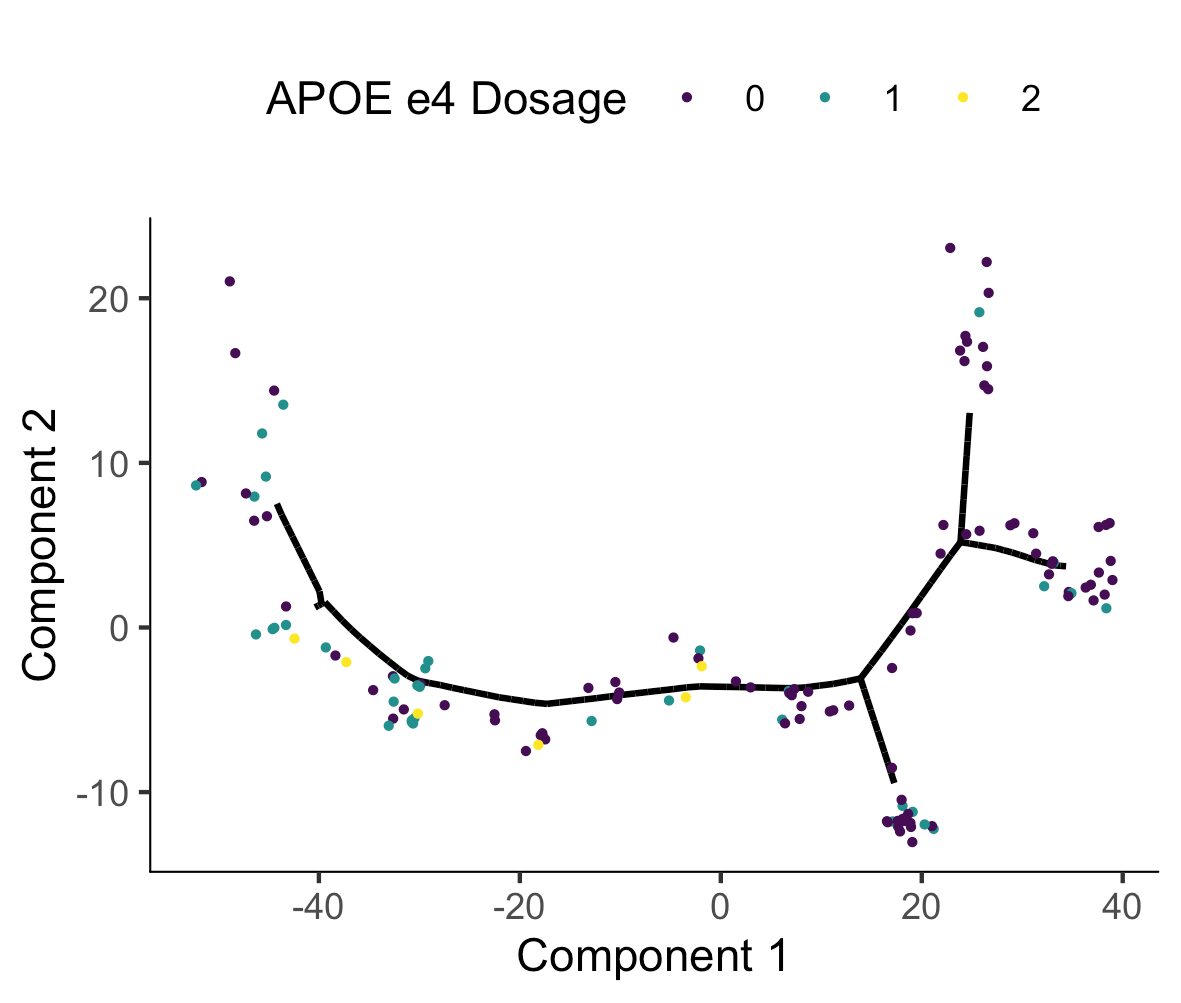

**Supplementary Figure 20 -** DLPFC manifolds with samples colored by inferred disease state.

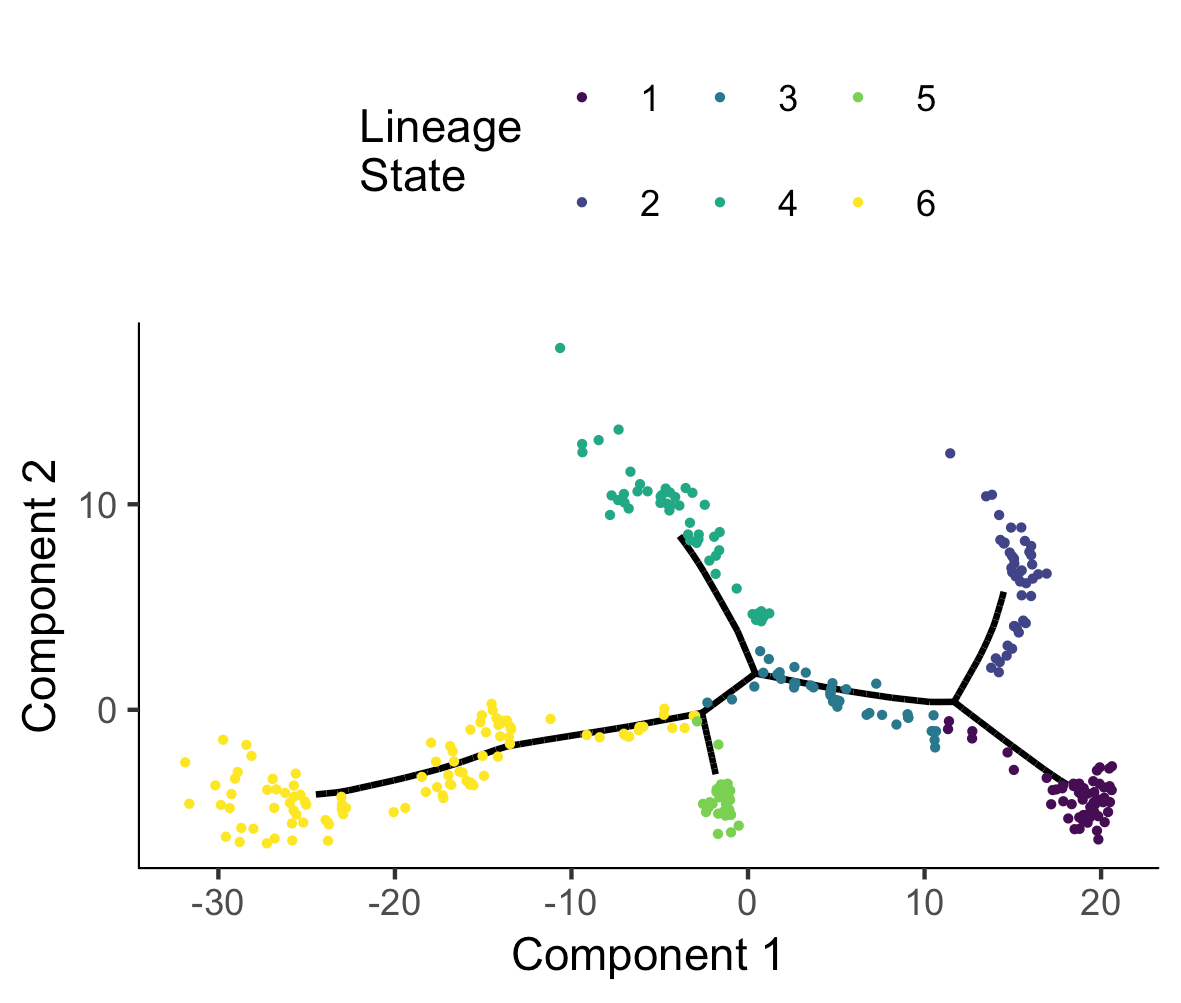

**Supplementary Figure 21 -** Quantile-quantile plot for the association with pseudotime in 305 female patients in the ROS/MAP cohort. The graph shows the Q-Q plot for GWAs of pseudotime in the ROS/MAP cohort with a genomic Inflation factor (lambda) of 0.981.

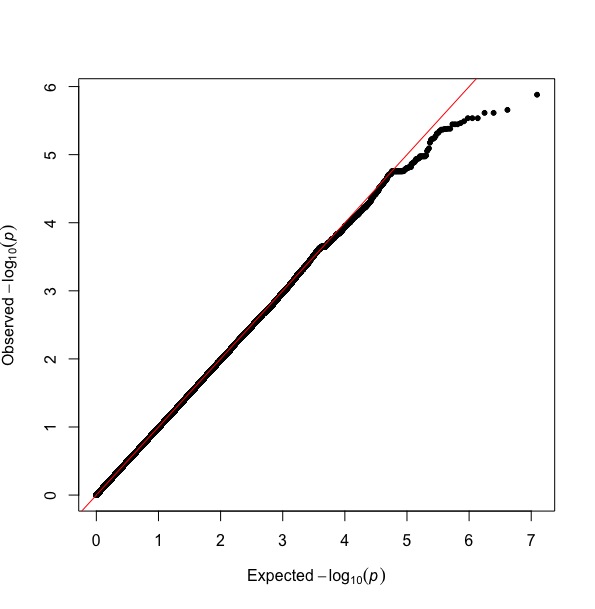

**Supplementary Figure 22 -** Quantile-quantile plot for the association with pseudotime in 131 female patients in the Mayo cohort.

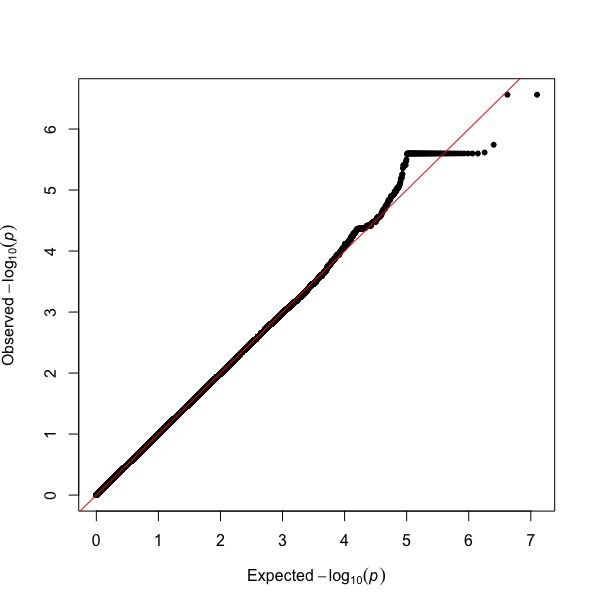

**Supplementary Figure 23 -** Manifold learning identified potential genetic factors of stage progression and subtypes of LOAD. A-B) GWA analysis was performed on the Mayo (A) and ROSMAP (B) cohorts using whole genome sequenced data and LOAD pseudotime as the phenotype. Despite the small sample sizes of both analyses (N = 131 in Mayo, N = 306 in ROSMAP), several genomic loci were identified harboring SNPs with a genome wide suggestive p-value (p < 1x10^-5^). These include several loci that were previously associated with LOAD or LOAD related endophenotypes (red labels; see also **Table S5**)

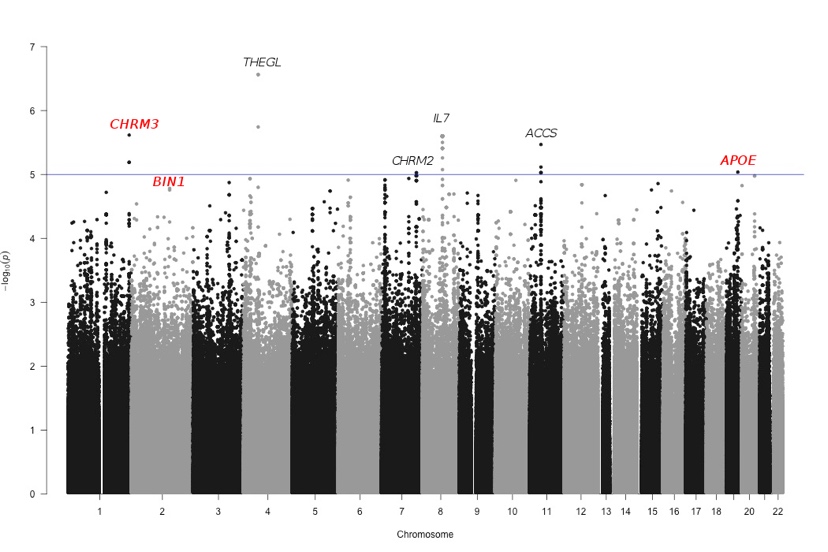

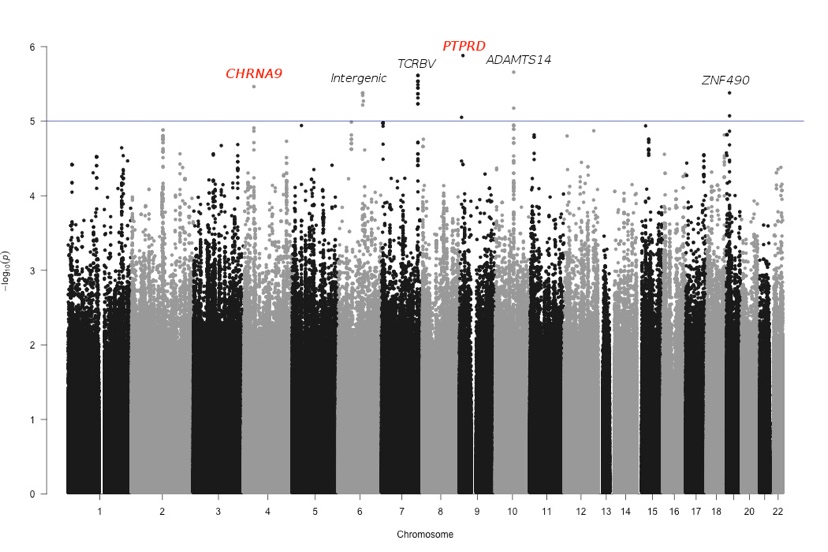

**A**

**B**

Chromosome

Chromosome

-Log_10_(p-value)

-Log_10_(p-value)

TCX

DLPFC

**Supplementary Figure 24 -** UpSet plot of branch differentially expressed genes from a two-sided Tukey’s honest significant test (FDR < 0.05) with branch one as reference branch in both studies respectively.

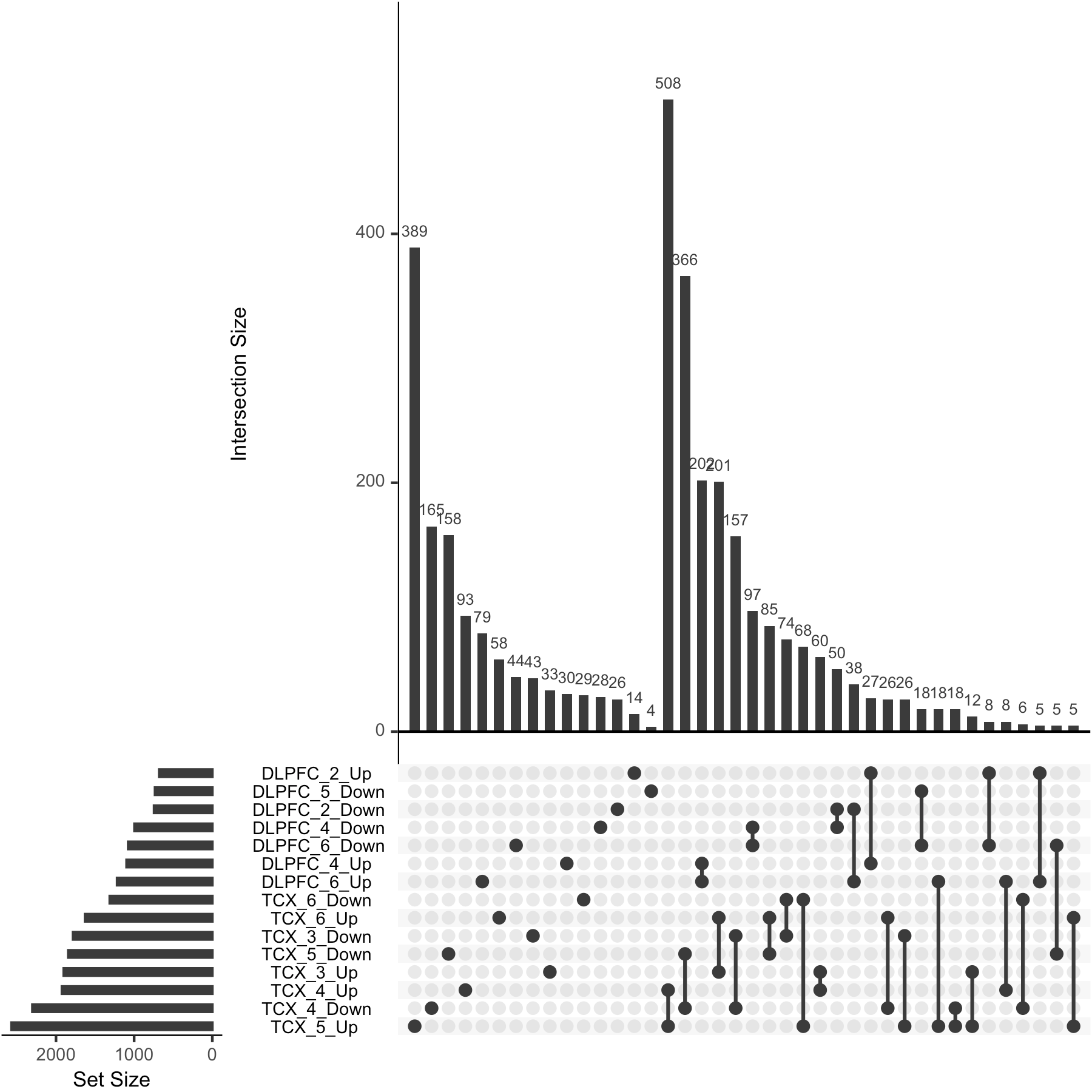

**Supplementary Figure 25 –** UpSet plot of comparison of clusters from Figure 4b from Mayo RNAseq lineage and differentially expressed genes from resistant individuals from the Mayo eGWAS study.

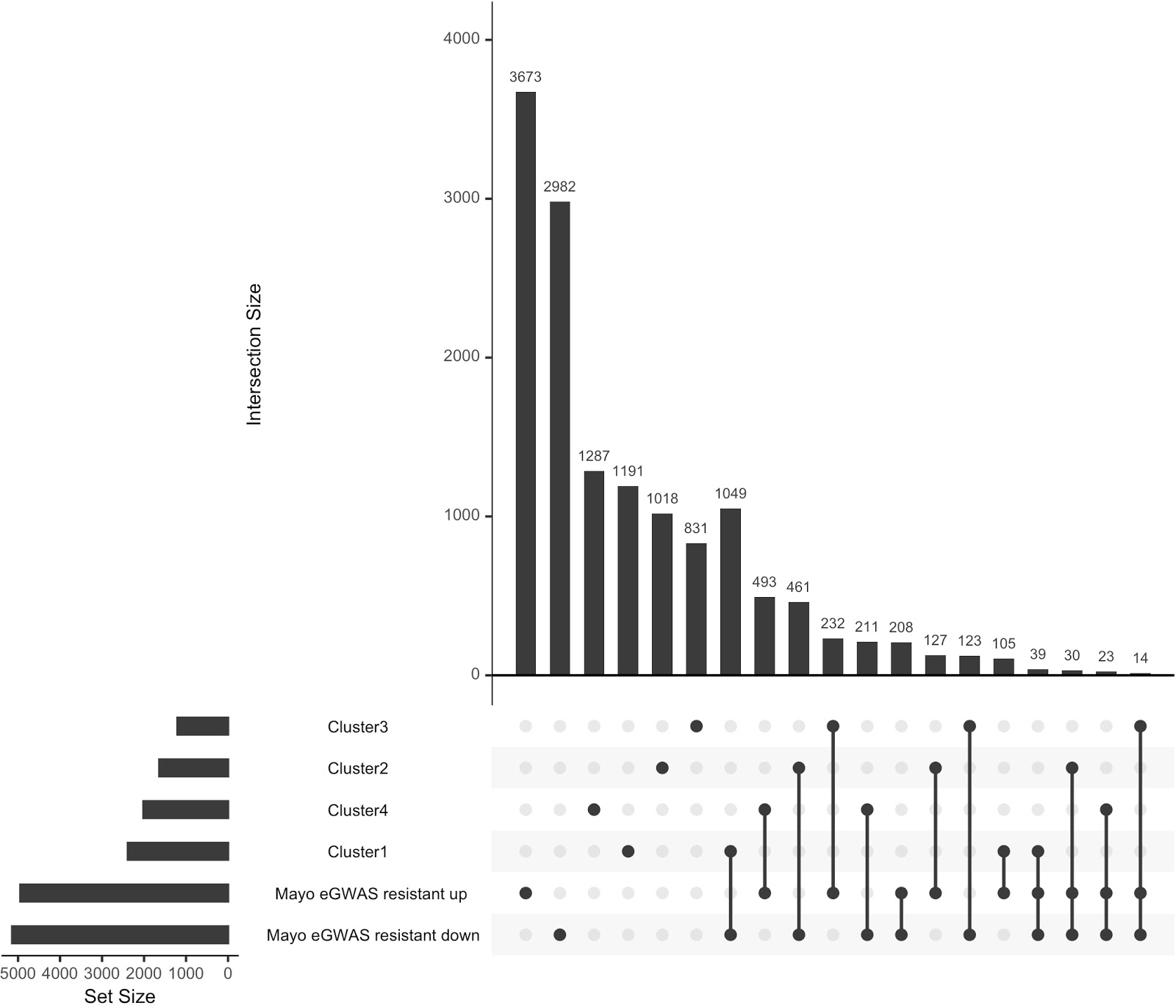

**Supplementary Figure 26 -** Comparison of different manifold learning methods for TCX brain region.

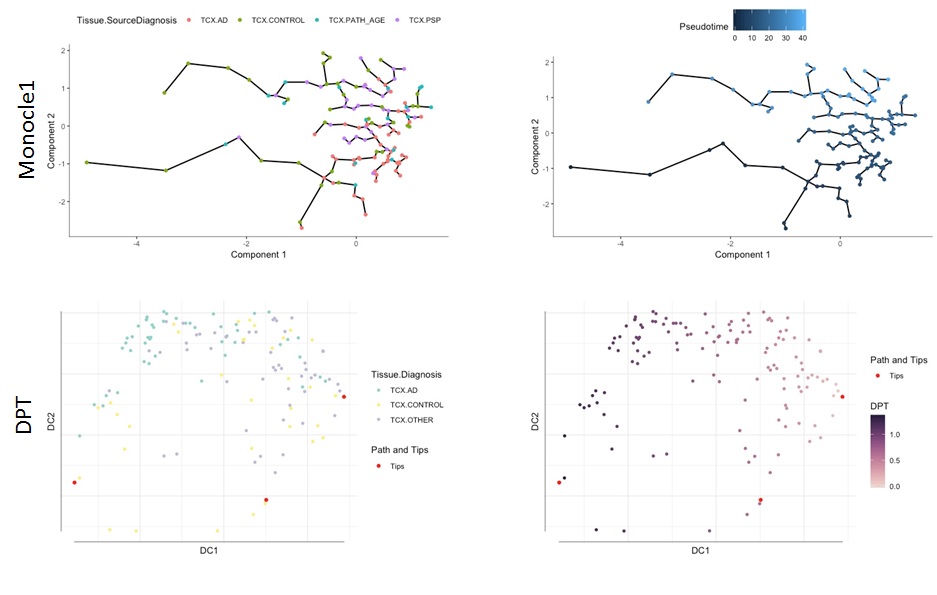

**Supplementary Figure 27 -** Comparison of different manifold learning methods for DLPFC brain region.

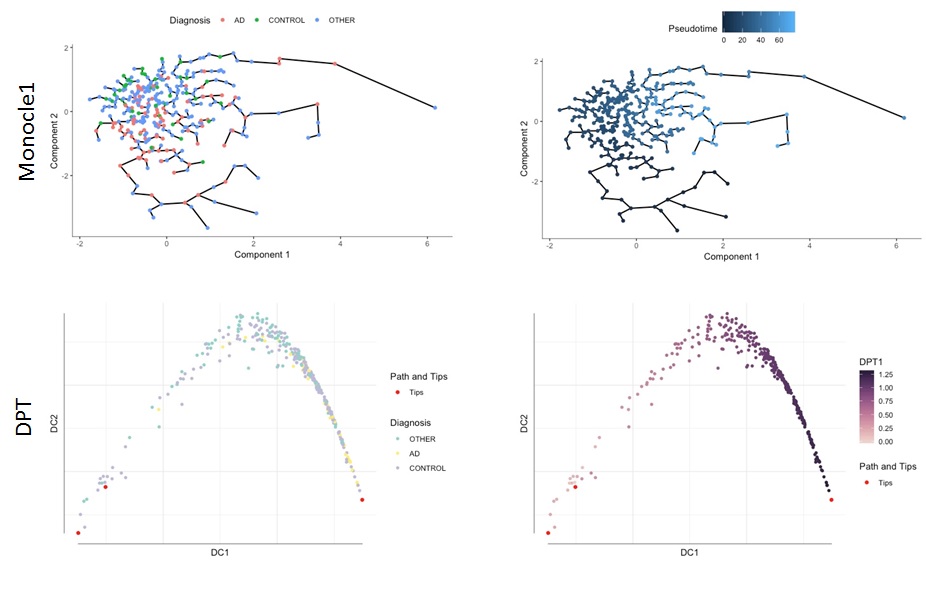

**Supplementary Figure 28 -** Correlation between pseudotimes estimated by different manifold learning approaches on both TCX and DLPFC brain region.

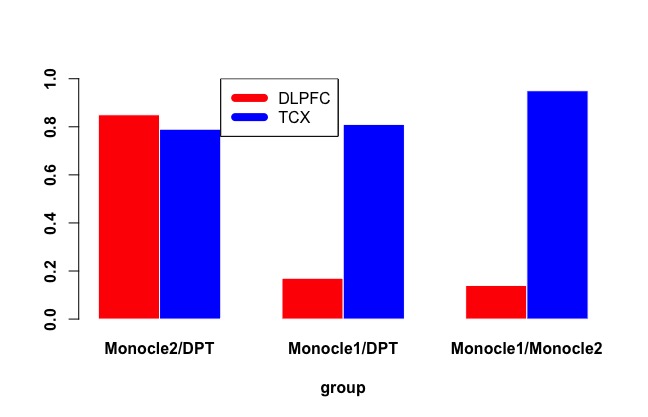

**Supplemental Tables**

**Supplementary Table 1**

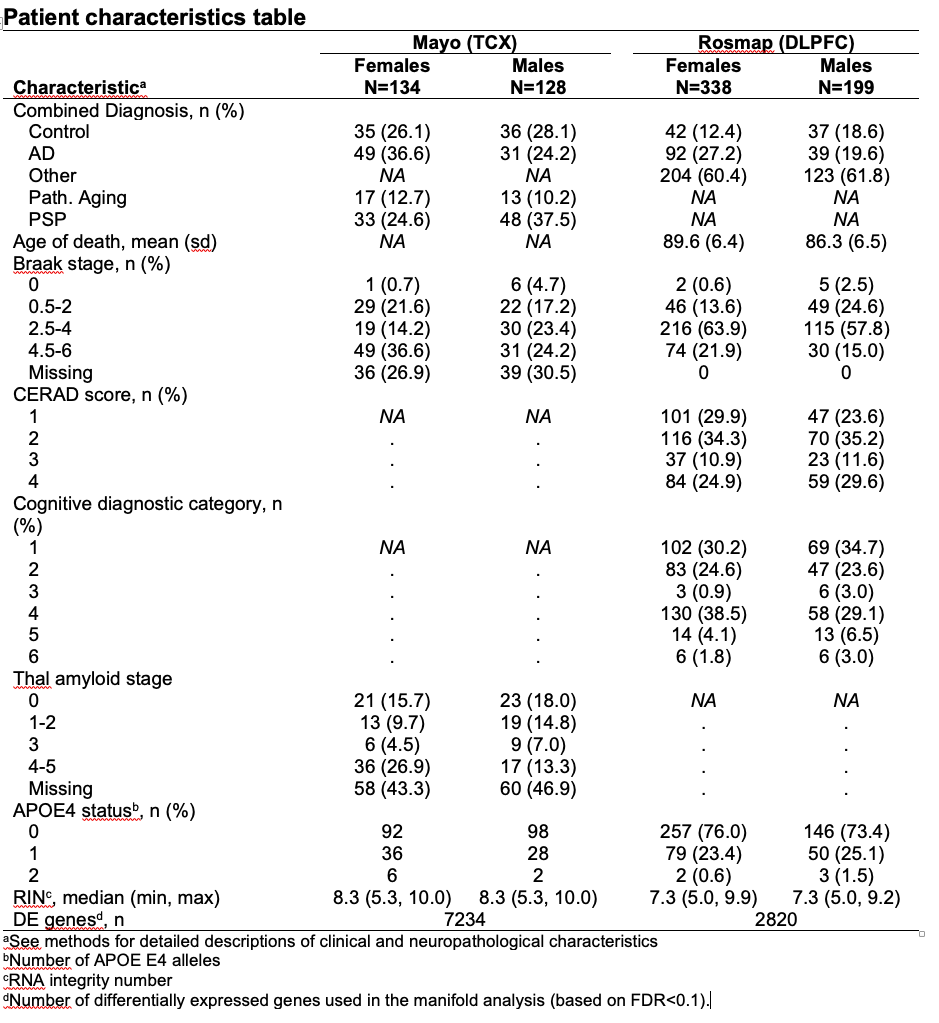

**Supplementary Table 2 -** Results of logistic regression for the association between unadjusted and adjusted pseudotime calculations (scaled) and AD case-control status.

| DLPFC | | | | |
| --- | --- | --- | --- | --- |
| PS adjustment | Coefficient | Std. Error | z value | P value |
| None | 1.7398 | 0.4075 | 4.27 | 1.96E-05 |
| RIN number | 1.3975 | 0.5732 | 2.438 | 0.0148 |
| PMI | 1.4923 | 0.3721 | 4.01 | 6.06E-05 |
| 1st 10 PCs | 1.1616 | 0.4485 | 2.59 | 0.00961 |
| ALL | 1.5486 | 0.5223 | 2.965 | 0.00303 |
| TCX | | | | |
| PS Adjustment | Coefficient | Std. Error | z value | P value |
| None | 1.0118 | 0.435 | 2.326 | 0.02 |
| RIN number | 1.6223 | 0.5035 | 3.222 | 0.00127 |
| PMI | 1.6097 | 0.4865 | 3.309 | 0.000938 |
| 1st 10 PCs | 1.545 | 0.498 | 3.103 | 0.00192 |
| ALL | 2.0967 | 0.5278 | 3.973 | 7.10E-05 |

**Supplementary Table 4 –** Associations between Braak, CERAD, and cogdx with features from alternative dimensionality reduction approaches.

| DLPFC (P-value from logistic ordinal regression) | | | |
| --- | --- | --- | --- |
| Feature | Braak | CERAD | cogdx |
| PCA1 | 1.91x10^-4^ | 4.51x10^-4^ | 3.97x10^-5^ |
| PCA2 | 0.800 | 0.201 | 1.07x10^-2^ |
| tSNE1 | 0.857 | 0.979 | 0.461 |
| tSNE2 | 1.02x10^-3^ | 1.11x10^-4^ | 1.23x10^-5^ |
| UMAP1 | 0.0219 | 9.63x10^-4^ | 1.34x10^-4^ |
| UMAP2 | 1.77x10^-6^ | 2.98x10^-6^ | 3.27x10^-7^ |
| Pseudotime (DDRTree) | 1.01x10^-5^ | 1.77x10^-5^ | 3.48x10^-7^ |
| Pseudotime (UMAP) | 1.34x10^-4^ | 6.64x10^-4^ | 2.18x10^-5^ |

**Supplementary Table 6 -** Association between mean expression of cell specific signatures and inferred disease severity (pseudotime).

| Study (Brain Region) | Cell Signature | P-value | R^2^ |
| --- | --- | --- | --- |
| Mayo RNAseq (TCX) | Neuronal | 3.6x10^-42^ | 0.76 |
|  | Microglial | 9.1x10^-29^ | 0.61 |
|  | Oligodendroglial | 6.7x10^-11^ | 0.28 |
|  | Astrocytic | 6.7x10^-22^ | 0.51 |
| ROS/MAP (DLPFC) | Neuronal | 1.6x10^-78^ | 0.65 |
|  | Microglial | 1.5x10^-31^ | 0.33 |
|  | Oligodendroglial | 1.4x10^-44^ | 0.44 |
|  | Astrocytic | 1.0x10^-50^ | 0.48 |

| SNP (dbSNP 150) | Location (hg19) | Nearest Gene(s) | region | A1 (Effect Allele) | A2 | Allele Freq.  (A1) | Beta  (Pseudotime) | SE (beta) | P | Cohort | Previous Association |
| --- | --- | --- | --- | --- | --- | --- | --- | --- | --- | --- | --- |
| rs4421019 | 4:40309851 | *CHRNA9* | *intergenic* | T | A | 0.35 | -6.18 | 1.31 | 3.44E-06 | ROS/MAP | LOAD |
| rs12216400 | 6:96292130 | *intergenic* | *intergenic* | A | G | 0.24 | 6.86 | 1.46 | 4.17E-06 | ROS/MAP | / |
| rs1573618 | 7:142244415 | *TCRBV* | *intronic* | T | C | 0.44 | -6.22 | 1.29 | 2.43E-06 | ROS/MAP | / |
| rs7870388 | 9:8660693 | *PTPRD* | *intronic* | G | C | 0.21 | -6.40 | 1.42 | 1.32E-06 | ROS/MAP | Tangle burden |
| rs4746059 | 10:72465488 | *ADAMTS14* | *intronic* | G | A | 0.42 | 5.85 | 1.21 | 2.20E-06 | ROS/MAP | / |
| rs55786848 | 19:12669655 | *ZNF490;*  *ZNF564* | *intergenic* | C | T | 0.15 | 8.01 | 1.71 | 4.16E-06 | ROS/MAP | / |
| rs12136200 | 1:240138130 | *CHRM3* | *intergenic* | C | T | 0.39 | -16.61 | 3.36 | 2.42E-06 | Mayo | Plaque burden |
| rs73818121 | 4:57397157 | *THEGL* | *exonic* | G | C | 0.07 | 33.19 | 6.63 | 1.81E-06 | Mayo | / |
| rs7809318 | 7:136419969 | *CHRM2* | *intergenic* | C | T | 0.07 | -34.03 | 7.37 | 9.41E-06 | Mayo | / |
| rs3808616 | 8:79868493 | *IL7* | *intergenic* | G | A | 0.35 | -17.70 | 3.59 | 2.51E-06 | Mayo | / |
| rs11037791 | 11:44022056 | *ACCS;*  *ACCSL* | *intergenic* | A | G | 0.49 | -16.41 | 3.38 | 3.39E-06 | Mayo | / |
| rs6857 | 19:45392254 | *PVRL2;*  *TOMM40;*  *APOE* | *intronic* | C | T | 0.17 | -18.23 | 3.95 | 9.18E-06 | Mayo | LOAD,  Tangle burden, Plaque burden |

**Supplementary Table 7 -** Overview of suggestive (p < 10^-5^) results from single variant association with pseudotime. Unadjusted p-values for a two sided likelihood ratio test in a linear regression model are shown.

**Supplementary Table 8 -** Associations of known AD variants associated with pseudotime in the IGAP cohort. Unadjusted p-values for a two sided likelihood ratio test in a linear regression model are shown for pseudotime.

| Chr. | Position (hg19) | SNP | Minor Allele Frequency | IGAP p- value  (Stage1+2) | Pseudotime  Cohort | Pseudotime p-value | Gene |
| --- | --- | --- | --- | --- | --- | --- | --- |
| 2 | 127887750 | rs62158731 | 0.26 | 3.41E-13 | Mayo | 4.68E-05 | *BIN1* |
| 3 | 151018968 | rs66927386 | 0.24 | 1.40E-04 | ROS/MAP | 0.0090 | *MED12L* |
| 6 | 32570051 | rs9270823 | 0.25 | 5.77E-10 | ROS/MAP | 0.0068 | *HLA-DRB1* |
| 7 | 99809921 | rs1727128 | 0.48 | 4.43E-06 | ROS/MAP | 0.0029 | *STAG3* |
| 9 | 129197516 | rs887656 | 0.11 | 1.40E-04 | ROS/MAP | 0.0079 | *MVB12* |
| 10 | 72524413 | rs2688767 | 0.36 | 1.39E-04 | ROS/MAP | 0.0078 | *ADAMTS14* |
| 11 | 85862728 | rs72962020 | 0.13 | 8.09E-06 | Mayo | 0.0075 | *PICALM* |
| 16 | 11199352 | rs12929596 | 0.13 | 6.43E-05 | ROS/MAP | 0.0067 | *CLEC16A* |
| 19 | 45392254 | rs6857 | 0.17 | 1.06E-15 | Mayo | 9.18E-06 | *APOE* |
| 20 | 55020557 | rs16979933 | 0.09 | 1.08E-07 | Mayo | 0.0054 | *CASS4* |

**Supplementary Table 11** - Number of genes differentially expressed at an FDR of 0.05 between the control branch (Branch 1) and other branches based on a two sided Tukey’s Honest significant difference test in an ANOVA model.

| Study (Brain Region) | Change in expression | Branch 2 | Branch 3 | Branch 4 | Branch 5 | Branch 6 |
| --- | --- | --- | --- | --- | --- | --- |
| ROSMAP (DLPFC) | Increased | 718 | 468 | 1121 | 662 | 1239 |
|  | Decreased | 781 | 611 | 1017 | 783 | 1094 |
| MayoRNAseq (TCX) | Increased | 506 | 2067 | 2034 | 2733 | 1815 |
|  | Decreased | 699 | 1912 | 2441 | 1966 | 1494 |

**Supplemental Table Legends**

**Supplementary Table 3**: AD LOAD GWAS genes^23^. Genes are from Tables 1-3 from previously published work^23^.

**Supplementary Table 5:** Cell specific gene sets used to compute mean expression of cell signatures across the lineages, as previously described^32^.

**Supplementary Table 9**: Summary statistics from differential expression analysis in DLPFC. Tukey’s honest significant difference test with branch 1 as reference is used, where unadjusted p-values from a two sided t-test for the mean difference are shown.

**Supplementary Table 10**: Summary statistics from differential expression analysis in TCX. Tukey’s honest significant difference test with branch 1 as reference is used, where unadjusted p-values from a two sided t-test for the mean difference are shown.

**Supplementary Table 12**: Significant GO pathway enrichments (FDR < 0.05) for DLPFC differential expressed gene sets.

**Supplementary Table 13**: Significant GO pathway enrichments (FDR < 0.05) for TCX differential expressed gene sets.

**Supplementary Table 14**: Significant GO pathway enrichments from biclustering analysis of mean expression of six branches (states) in TCX with four clusters.
